# Discovering Haematoma-Stimulated Circuits for Secondary Brain Injury after Intraventricular Haemorrhage by Spatial Transcriptome Analysis

**DOI:** 10.1101/2022.12.06.519018

**Authors:** Le Zhang, Jiayidaer Badai, Guan Wang, Xufang Ru, Wenkai Song, Yujie You, Jiaojiao He, Suna Huang, Hua Feng, Runsheng Chen, Yi Zhao, Yujie Chen

## Abstract

Central nervous system (CNS) diseases, such as neurodegenerative disorders and brain diseases caused by acute injuries, are important yet challenging to study due to disease leisions’ locations and other complexities. Utilizing the powerful spatial transcriptome analysis together with novel algorithms we developed for the study, we report here for the first time a 3D trajectory map of gene expression changes in the brain following acute neural injury using a mouse model of intraventricular haemorrhage (IVH). IVH is a common and representative complication after various acute brain injuries with severe mortality and mobility implications. Our data identified three main 3D global pseudospace-time trajectory bundles which represent the main neural circuits from the lateral ventricle to the hippocampus and primary cortex induced by experimental intraventricular haematoma stimulation. Further analysis indicated a rapid response in the primary cortex, as well as a direct and integrated effect on the hippocampus after IVH. These results are informative in understanding the pathophysiological changes, including the spatial and temporal patterns of gene expression changes, in IVH patients after acute brain injuries and strategizing more effective clinical management regimens. The bioinformatics strategies will also be useful for the study of other CNS diseases.

## 1. Introduction

Intraventricular haemorrhage (IVH) refers to bleeding occurring inside the ventricles of the brain or haematoma breaking from the periventricular brain parenchyma, which is usually secondary to traumatic brain injury, subarachnoid haemorrhage, or spontaneous intracerebral haemorrhage. IVH is one of the common complications after various acute brain injuries^1–3^, leading to severe mortality and mobility, with great burdens on economics and society. To date, the gold standard for the treatment of IVH is haematoma removal and hydrocephalus prevention^4^ due to the pathophysiological mechanisms of relieving mass effects and elevated intracranial pressure towards the lateral walls of ventricles^3^, as well as obstructions and inflammatory responses from blood metabolic stimulations^5^. However, a recently completed large-scale randomized clinical trial (CLEAR III, Trial No. NCT00784134) employing alteplase to effectively clear intraventricular haematoma did not demonstrate adequate neurobehavioral benefits for IVH patients^6^. With the development of neurosurgery and neurocritical care, the direct mortality from IVH has gradually decreased, leaving us to understand and alleviate the neurological dysfunction after IVH for better outcomes of these patients. Nevertheless, the pathophysiology of how these haematoma stimulations in ventricles cause neurological dysfunction remains unknown.

For this reason, this study investigated the pathophysiological mechanism and related intervention strategies for secondary neurological dysfunction after IVH by developing a bioinformatics analysis workflow based on spatial transcriptome sequencing^7,8^. First, by constructing 3D pseudospace-time trajectories^9^, we identified the cell subtypes generated after IVH and their potential cell types. Then, we explored and validated the important pathophysiological mechanisms for these cell subtypes by carrying out cell–cell communication strength ^9^ and pathway analyses^10–14^.

In summary, our study not only identified the brain regions affected by secondary neurological dysfunction after IVH^15–17^ from a 3D perspective but it also has the potential for us to construct corresponding intervention strategies for acute traumatic brain trauma at the molecular and cellular levels in the distant future.

## 2. Results

### 2.1 Developing a spatial transcriptome sequencing-based bioinformatics analysis workflow to investigate IVH

In order to comprehensiavely dissect the complex gene expression changes following acute brain injury, using the powerful spatial transcriptomics technology and IVH as an example here, we first developed a spatial transcriptomics sequencing-based bioinformatic analysis workflow (Figure 1).

**Figure 1:**
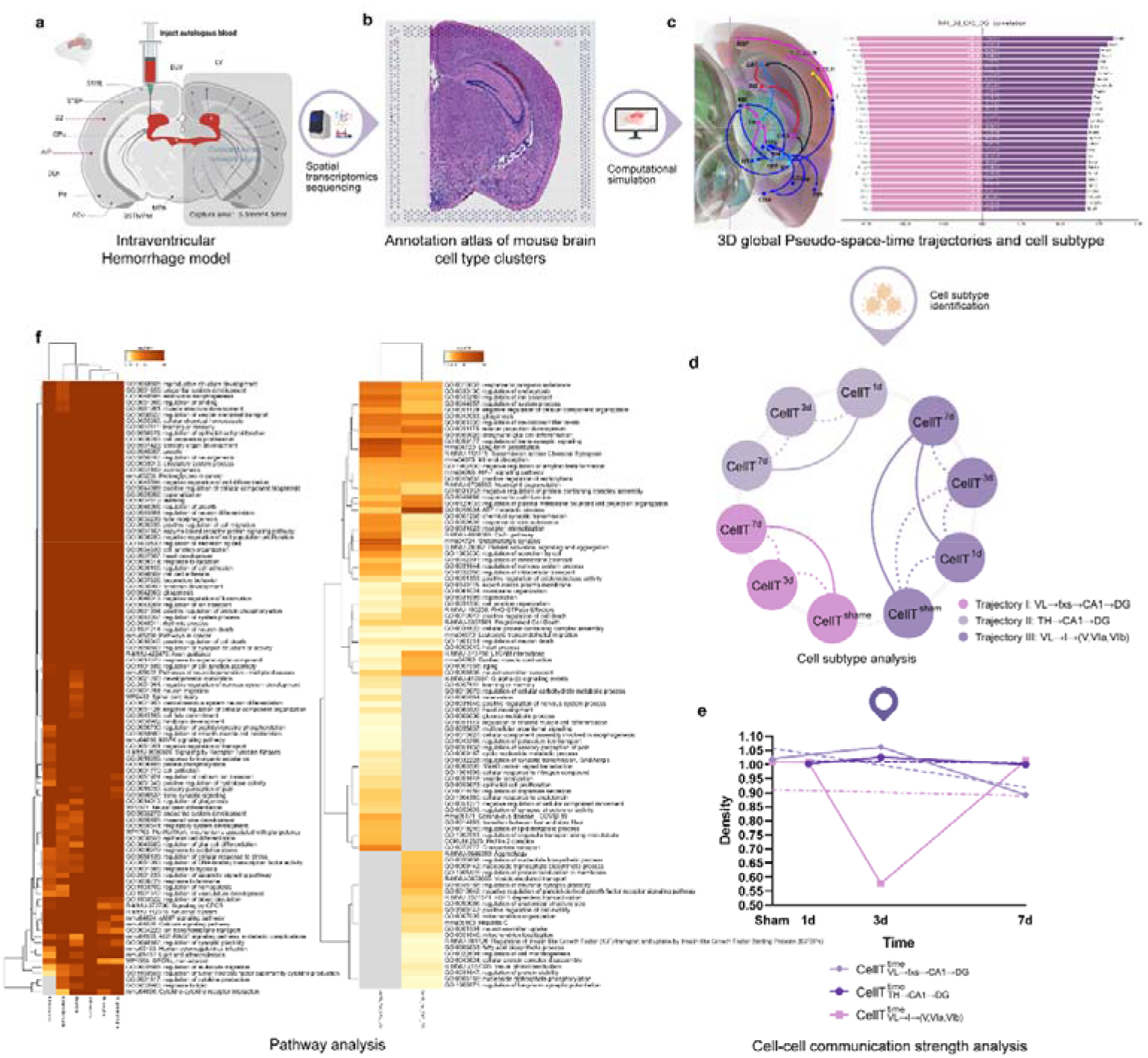
The workflow of the study. **a,** Secondary intraventricular haemorrhage experiment. **b,** Annotation atlas of mouse brain cell type clusters. **c,** 3D global pseudospace-time trajectories and the transition gene sets. **d,** Cell subtype analysis. **e,** Cell-cell communication strength analysis. **f,** Pathway analysis.

First, given the difficulty in perfoming the analysis using human patient samples, we constructed an IVH model (Figure 1a) in mice by injecting autologous blood into the lateral ventricle (for technical details please refer to Experimental Procedures).

Spatial transcriptomics sequencing^7,8^ and the SME clustering algorithm^9^ were subsequently used to analyze frozen sections of IVH mouse brain to obtain the annotation atlas of cell type clusters (Figure 1b and Visium spatial transcriptome sequencing of Methods section) and to construct 3D global pseudospace-time trajectories (Figure 1c) and transition gene sets. More extensive cell subtype characterizations were performed on the three 3D global pseudospace-time trajectories identified and at different times (Figure 1d). We further explored the molecular-level changes of these cell subtypes (Figure 1e) by using a newly-developed cell–cell interaction intensity density algorithm and interrogated the biological characteristics of these cell subtypes using pathway analysis (Figure 1f).

### 2.2 3D global pseudospace-time trajectory reconstruction for mouse brain tissue after lateral ventricle haemorrhage

Initially, we took 19 original frozen sections of the brains of 5 mice with IVH as the input. After carrying out spatial transcriptome sequencing^7,8^, we obtained H&E (haematoxylin and eosin) staining slices and examined spatial gene expression (Figure 2a, SR1. Figure 1 and SR1. Table 1).

**Figure 2:**
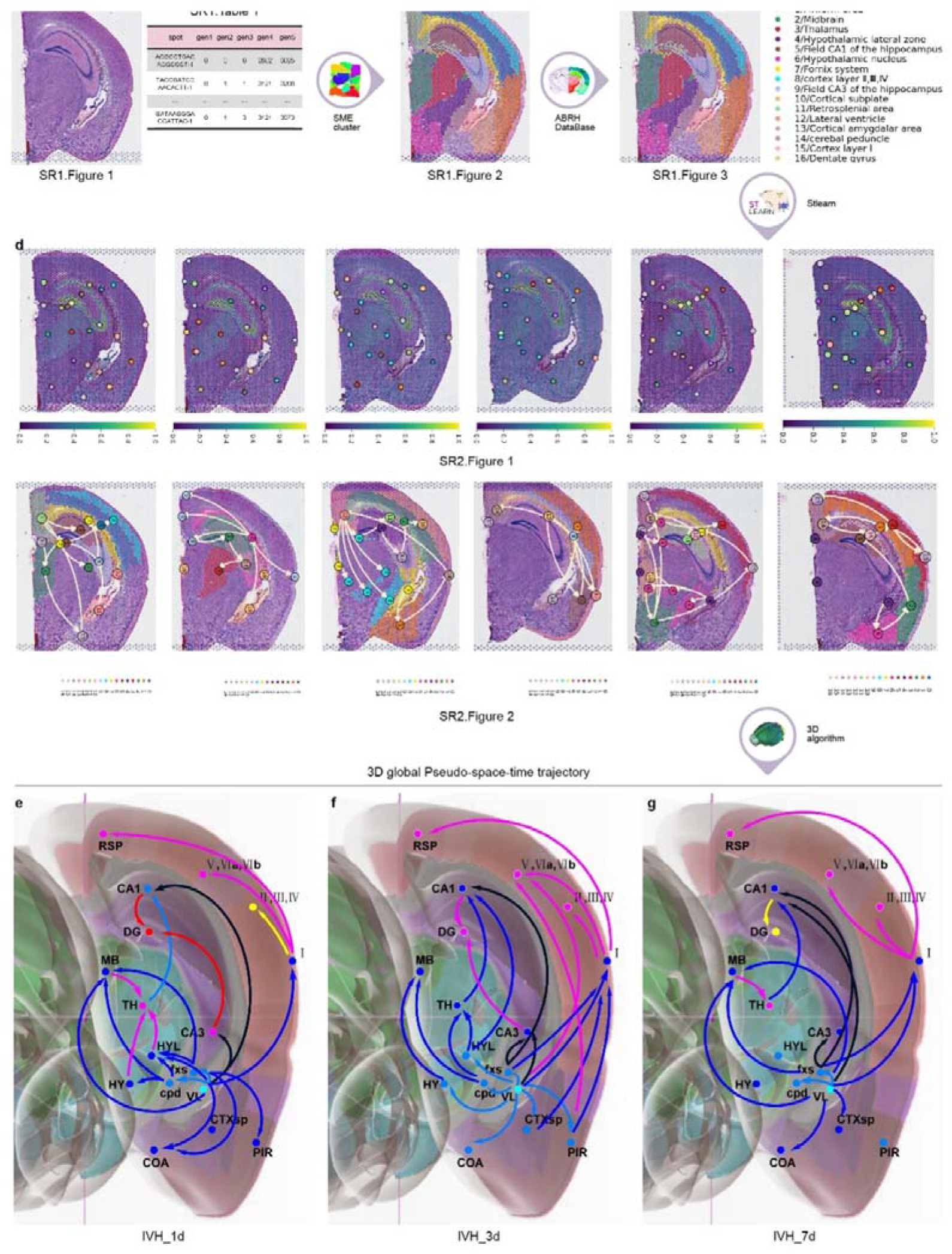
The process to obtain the 3D global pseudospace-time trajectory after IVH. **a,** H&E (haematoxylin and eosin stain) slices and spatial gene expression. **b,** Normalization and clustering of the cell types with similar gene expression. **c,** Annotation atlas of cell type clusters. **d**, Planar diffusion pseudotime plots and planar global pseudospace-time trajectories. **e**, The 3D global pseudospace-time trajectories after IVH on the 1^st^ day. **f**, The 3D global pseudospace-time trajectories after IVH on the 3^rd^ day. **g**, The 3D global pseudospace-time trajectories after IVH on the 7^th^ day.

Next, we employed the SME algorithm^9^ to normalize and cluster the cell types with similar gene expression (Figure 2b and SR1. Figure 2) by inputting SR1. Figure 1 and SR1. Table 1.

After that, we obtained the annotation atlas of cell type clusters (Figure 2c) in different brain regions by comparing the clustering results with the anatomical brain regions on the Allen Brain Reference Atlas map^18^, which describes the whole transcriptional signal in the brains of mice after IVH (SR1. Figure 3).

Then, we employed the stLearn algorithm^9^ to obtain the planar diffusion pseudotime plots and the planar global pseudospace-time trajectories (Figure 2d and SR2. Figure 1 &2) by inputting SR1. Figure 3.

Last, we employed our developed 3D global pseudospace-time trajectory reconstruction algorithm (SM. Tables 2 & 3) to obtain 3D global pseudospace-time trajectories (Figure 2e, 2f and 2g) by inputting SR2. Figures 1 & 2.

In addition to consciousness disturbance, it has been commonly observed that short-term motor dysfunction and long-term cognitive disorders occur after a single intraventricular haemorrhage. Table 1 classifies 3D global pseudospace-time trajectories(Figure 2e,2f and 2g)into three major trajectory bundles after haematoma stimulation based on their pivotal effect and their representative value for the neural circuits of motor dysfunction and cognitive disorder.

**Table 1.**
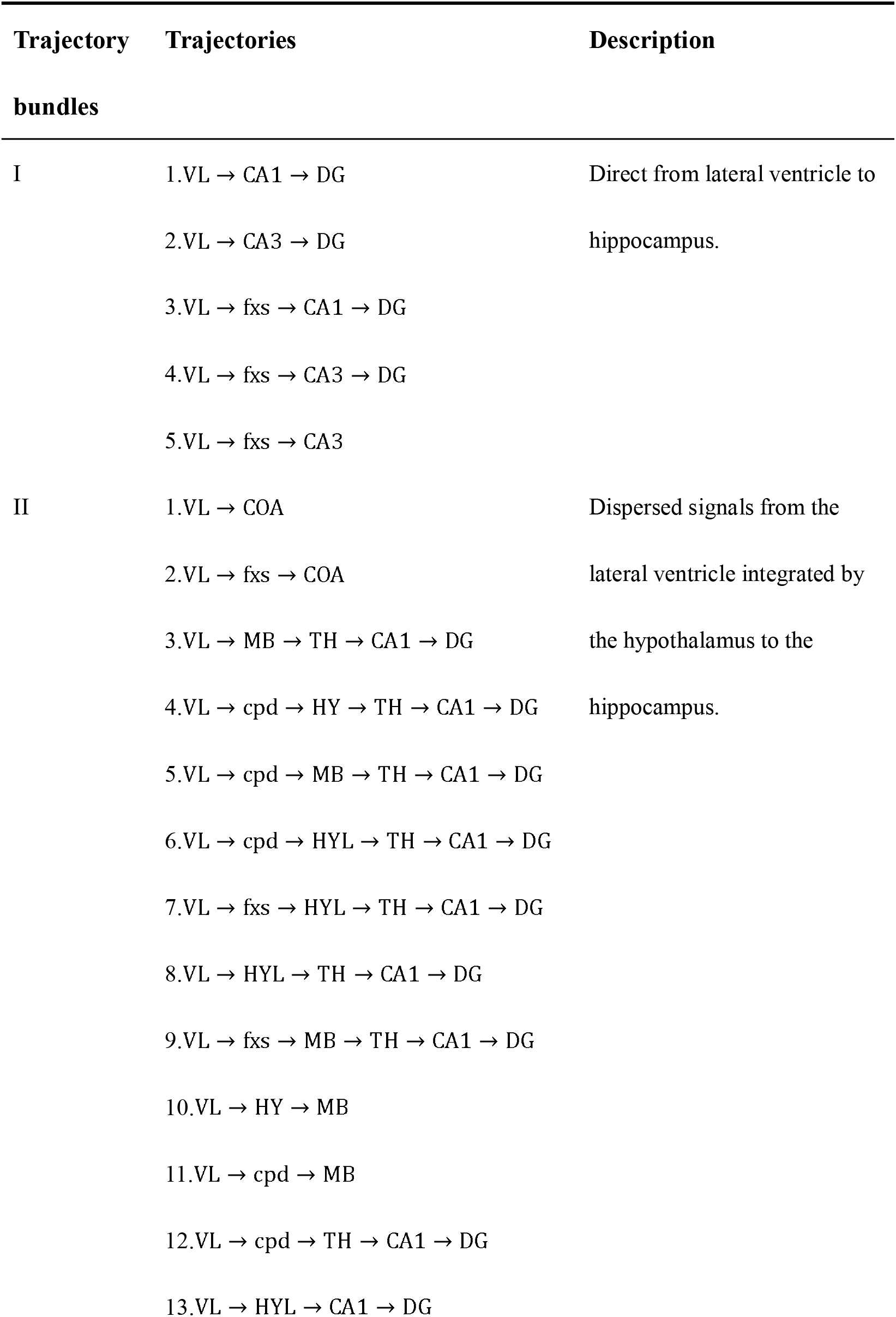
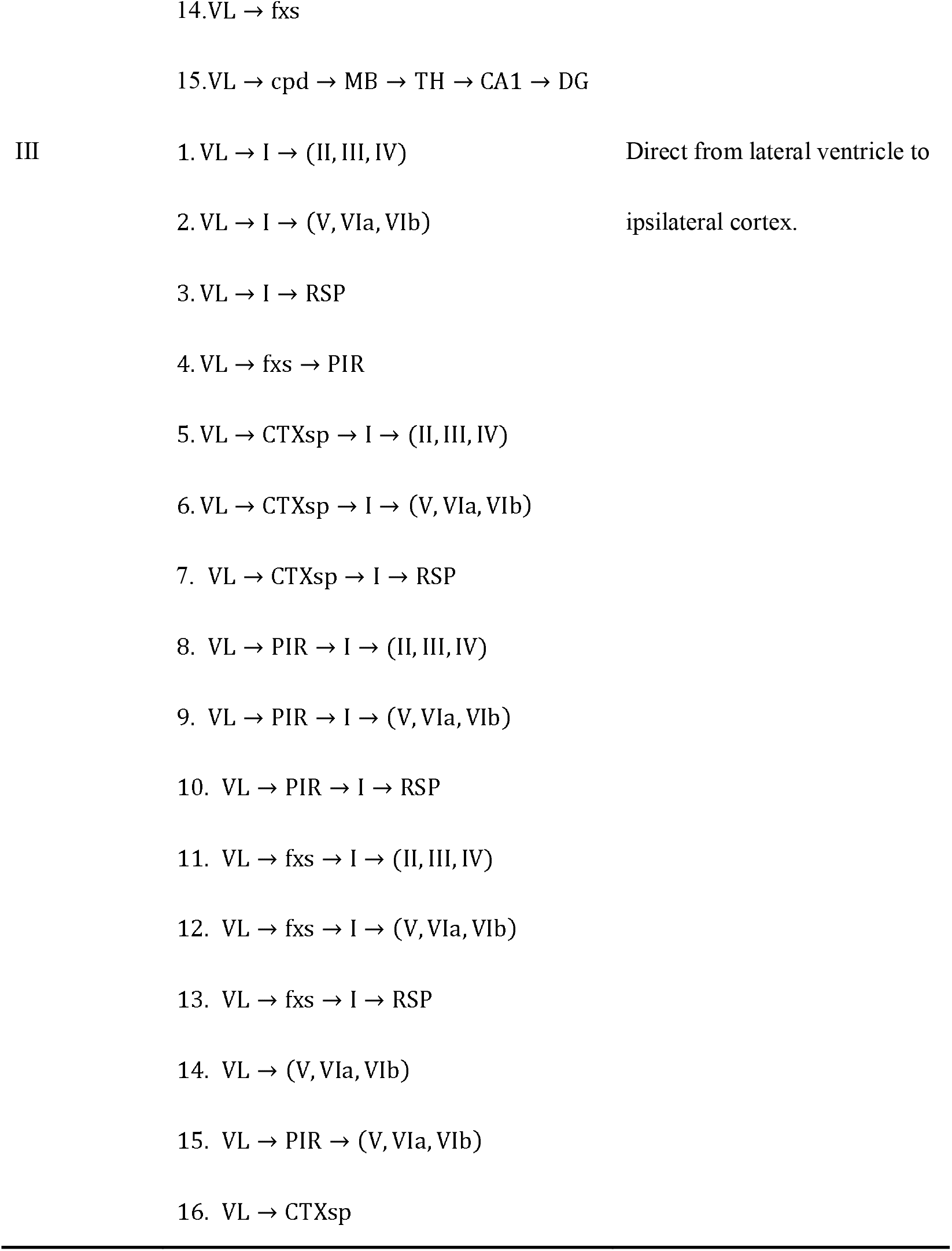
Three major trajectory bundles from the lateral ventricle to the hippocampus and primary cortex after haematoma stimulation.

Here, trajectory bundles I and II are described in SR2. Figure 3 b, e, h and SR2. Figure 3 c, f, i for different times, which extend from the lateral ventricle to the hippocampus. Trajectory bundle III is described by SR2. Figure 3 d, g, j for different times, which extend from the lateral ventricle to primary cortex.

The following analysis of trajectory bundle III explores how haematoma stimulation in the lateral ventricle affects the cortex and causes motor dysfunction, while that of trajectory bundles I and II illustrates direct pathophysiological mechanisms of cognitive dysfunction after IVH, with unique neural circuits for haematoma stimulation in the lateral ventricle.

### 2.3 The trajectory-based transition gene set for 3D global pseudospace-time trajectories

Initially, we employed the Spearman correlation analysis of stLearn^9,19–26^ to obtain the top 30 planar upregulated/downregulated transition genes (Figure 3a and SR3. Table 1) for each planar subtrajectory by inputting the planar global pseudospace-time trajectory (SR2. Figure 2).

**Figure 3:**
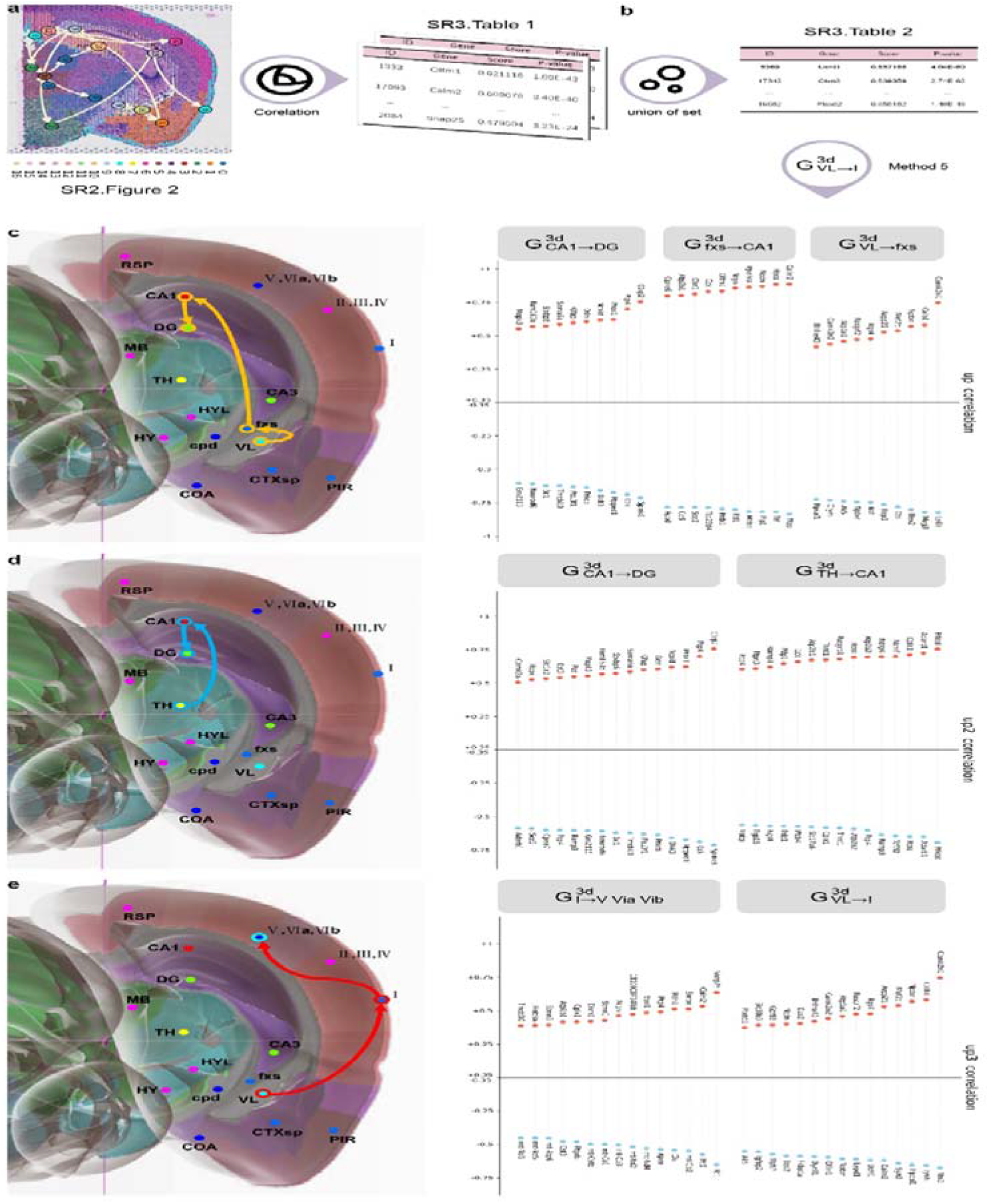
The process to obtain trajectory-based transition gene sets for 3D global pseudospace-time trajectories. **a,** The top 30 planar upregulated/downregulated transition genes for each planar subtrajectory. **b**, The top 30 3D upregulated/downregulated transition genes for each 3D subtrajectory. **c**, The top 30 3D trajectory-based transition gene set of VL→fxs→ CA1→DG. **d**, The top 30 3D trajectory-based transition gene set of TH→CA1→DG. **e**, The top 30 3D trajectory-based transition gene set of VL→I→(V,VIa,VIb).

Among the three major global pseudospace-time trajectory bundles (Table 1), we chose frequently appearing trajectories from each trajectory bundle at different time for further analysis, which are VL→fxs→CA1→DG from trajectory bundle I, TH→CA1 →DG from trajectory bundle II and VL→I→(V,VIa,VIb) from trajectory bundle III.

VL→fxs→CA1→DG from trajectory bundle I could be the representative mechanism of how haematoma stimulation in the lateral ventricle directly induces hippocampal activities and cognitive dysfunction.

TH→CA1→DG from trajectory bundle II could transmit multiple signals from the lateral ventricle and integrate them via the hypothalamus to the hippocampus, which might be a relatively long-term effect of haematoma stimulation.

The changes in VL→I→(V,VIa,VIb) from trajectory bundle III might reflect the acute and direct effect of haematoma stimulation on primary cortex.

Next, we carried out a union operation for the top 30 planar upregulated/downregulated transition genes for the same subtrajectory in different plane slices at the same time to obtain the top 30 3D upregulated/downregulated transition genes (Figure 3b and SR3. Table 2) for each 3D subtrajectory by inputting SR3. Table 1. When we performed a union operation for the top 30 planar upregulated/downregulated transition genes, if different plane slices had the same gene with a different sign (upregulated or downregulated), we chose the gene whose absolute value was the greatest.

Here, we use Figure 3c, 3d and 3e as examples to describe the top 30 3D upregulated/downregulated transition genes for each subtrajectory in VL→fxs→CA1 →DG from trajectory bundle I, TH→CA1→DG from trajectory bundle II, and VL→I→ (V,VIa,VIb) from trajectory bundle III.

### 2.4 Cell subtype analysis

As discussed in section 2.3, we chose VL→fxs→CA1→DG from trajectory bundle I, TH→CA1→DG from trajectory bundle II and VL→I→(V,VIa,VIb) from trajectory bundle III for further analysis.

Initially, we employed our developed algorithm (SM. Tables 4 & 5) to identify a cell subtype on the selected trajectory at different times (Figure 4a and SR4. Table 1) by inputting the top 30 upregulated/downregulated 3D transition genes for each 3D subtrajectory (SR3. Table 2), the marker gene sets of which are listed in Figure 4b and SR4. Table 2. This process is detailed in cell subtype identification of Methods section.

**Figure 4:**
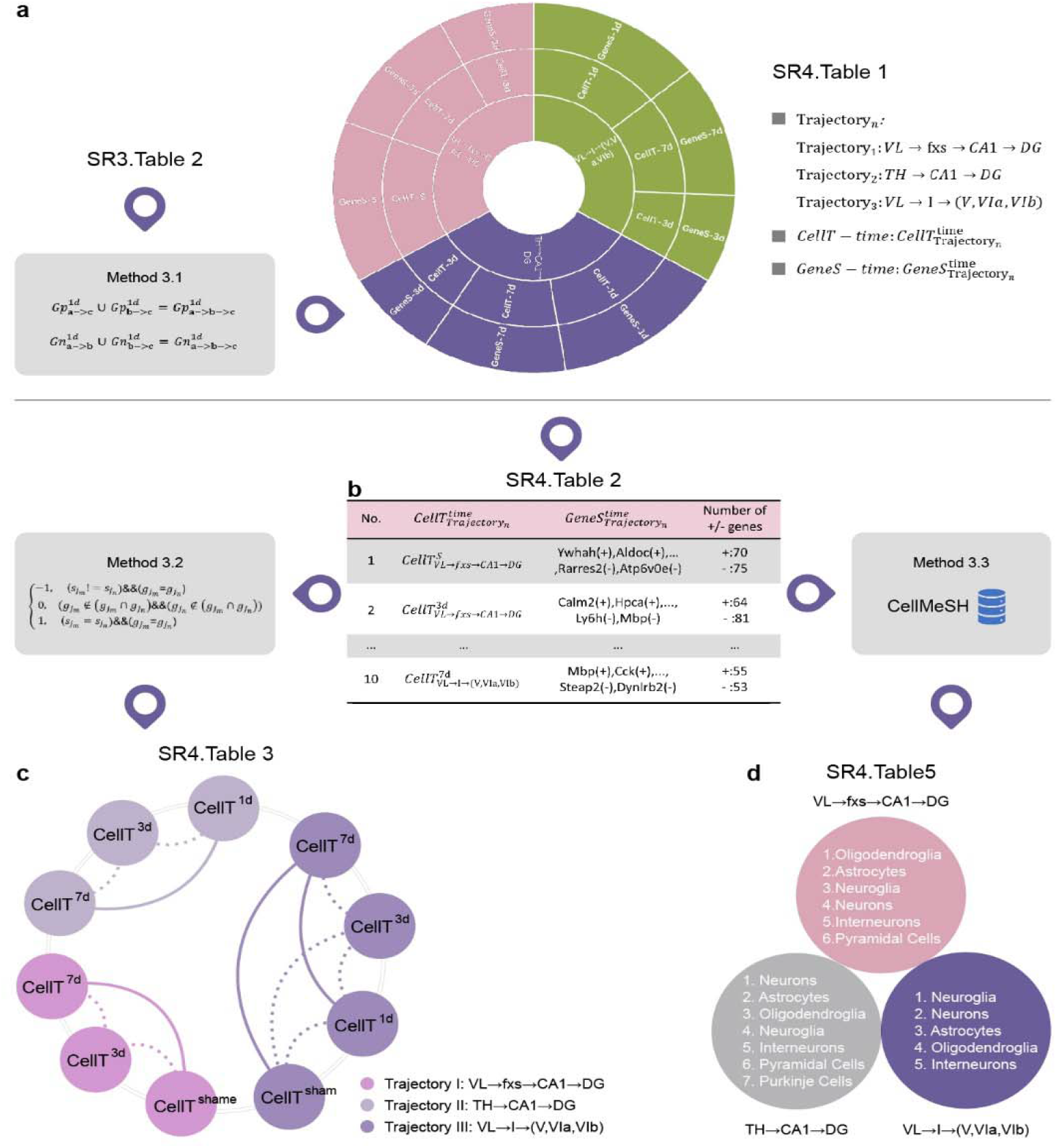
Cell subtype analysis. **a,** The cell subtypes corresponding to the selected trajectory at different times. **b,** The marker gene sets of cell subtypes corresponding to the selected trajectory at different times (SR4. Table 2). **c,** Similarity analysis of cell subtypes between different times on the same trajectory (SR4. Table 3). Among them, the thick line represents a high similarity between two time points, and the thin line represents a low similarity between two time points. **d,** The cell types shared by our identified cell subtypes at different times on the same trajectory (SR4. Table 5).

Then, we built up a similarity algorithm (SM. Tables 4 & 6) to describe the similarity between cell subtypes on the same trajectory at different times (SR4. Table 3) by inputting the marker gene sets (SR4. Table 2). This process is detailed in similarity algorithm for cell subtypes of Methods section.

Figure 4c shows that the cell subtype similarity between 3^rd^ day and 7^th^ day is small for each selected trajectory. Additionally, the cell subtype similarity between sham and 3^th^ day are both small, but the cell subtype similarity between sham and 7^th^ day are both great for *VL* → *fxs* → *CA1* → *DG* and VL → I → (V, VIa, VIb).

These findings indicate a pattern in which not only are the cell subtypes similar for *VL* → *fxs* → *CA1* → *DG*, TH → CA1 → DG and VL → I → (V, VIa, VIb) after IVH, but also these cell subtypes are different from the cell subtypes of the sham group with only the acupuncture effect.

It is noted that we do not have a cell subtype for the 1^st^ day or the sham group for *VL* → *fxs* → *CA1* → *DG* and TH → CA1 → DG, respectively. Additionally, Figure 4c and SR4. Table 3 show that the pattern is not so obvious for VL → I → (V, VIa, VIb).

Last, we found cell types (SR4. Table 4) from a commonly used public single-cell sequencing database (CellMeSH ^27^) that are similar to our identified cell subtypes (SR4. Table 1) by using search for similar cell types by CellMeSH of Methods section. Additionally, Figure 4d and SR4. Table 5 describe the cell types that are most like our identified cell subtypes at different times on the same trajectory.

### 2.5 Cell-cell communication strength analysis

Initially, we employed the CellTalkDB database^28^ to locate ligand receptors by inputting the upregulated and downregulated transition gene sets (SR3. Table 2). Figure 5a and SR5. Table 1 show the upregulated/downregulated transition ligand-receptor (*LR_n_*) sets.

**Figure 5:**
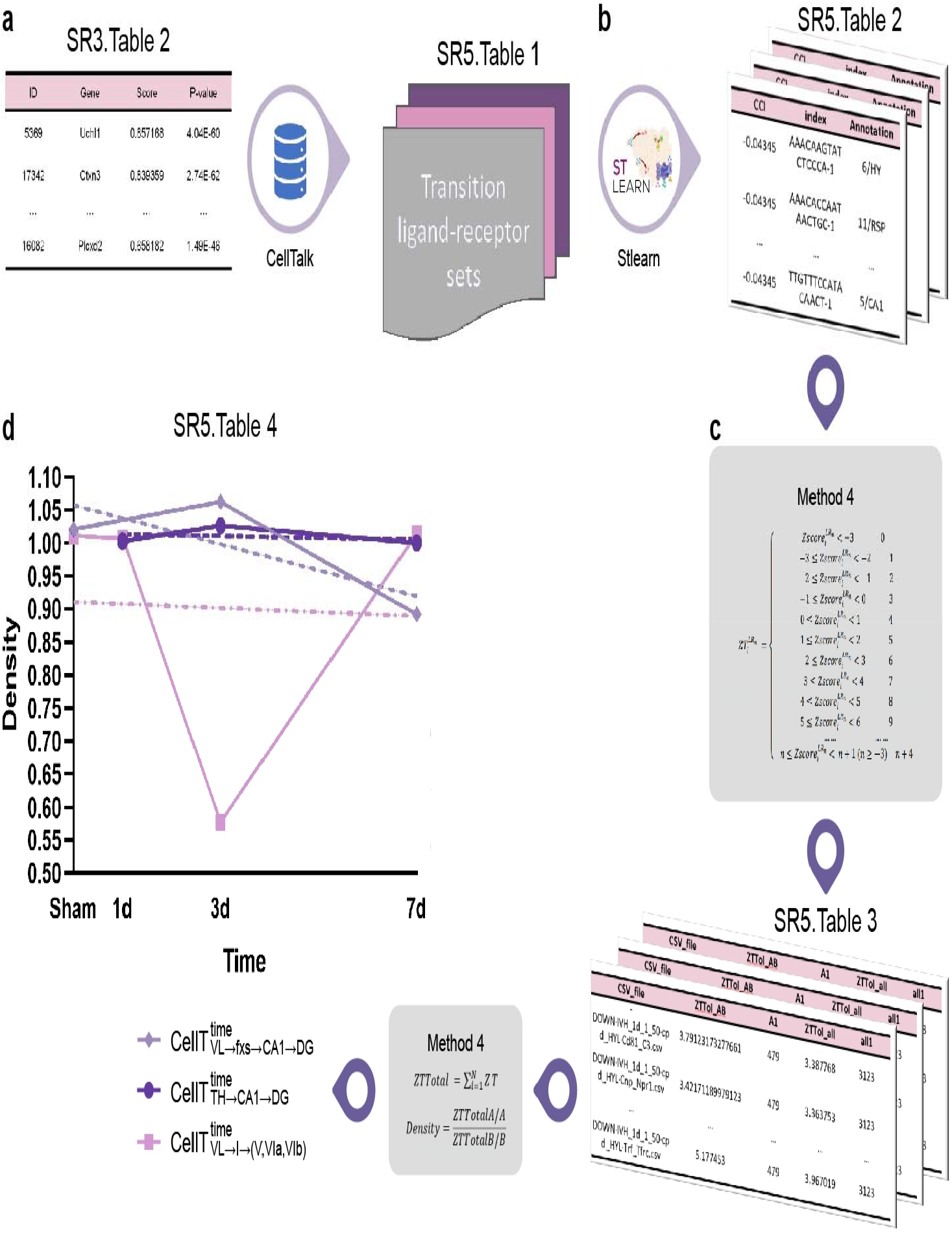
Cell-cell communication strength analysis. **a,** The transition ligand–receptor sets. **b,** The interaction intensity value 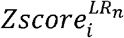. **c,** The discrete interaction intensity 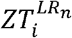. **d,** The cell-cell communication strength (*Density*).

Next, we employed stLearn^9^ to analyse their interaction intensity by using SR5. Table 1 as the input. Figure 5b and SR5. Table 2 show the interaction intensity values 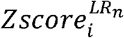 for the upregulated/downregulated transition ligand-receptor (*LR_n_*) sets. Then, we obtained the discrete interaction intensity 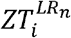(Figure 5c and SR5. Table 3) and the average *Density^LRn^* for each ligand receptor of the cell subtypes (Figure 5d and SR5. Table 4) by using Cell-cell communication analysis of Methods section.

Figure 5d shows that the *Density of* 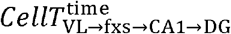 increased from 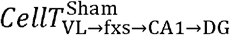 to 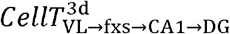 and decreased from 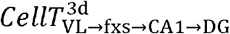 to 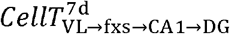; the *Density of* 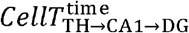 increased from 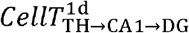 to 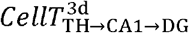 but decreased from 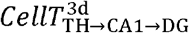 to 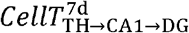; and the *Density of* 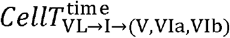 decreased from 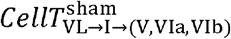 to 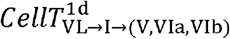 and from 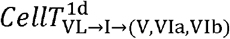 to 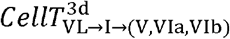 but increased from 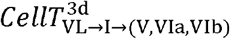 to 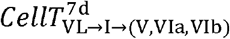.

Last, the Kruskal-Wallis test^29^ (SR5. Table 5) demonstrates that the density is significantly different among these cell subtypes on VL → fxs → CA1 → DG, TH → CA1 → DG and VL → I → (V, VIa, VIb).

Therefore, we hypothesized that the cell-cell communication strength would greatly change on the 3^rd^ day after IVH and return to the normal state on the 7^th^ day.

### 2.6 Pathway analysis

First, we employed CellMeSH ^27^ to identify the genes associated with similar cell types (Figure 6a and SR6. Table 1) by inputting their similar cell types (SR4. Table 5).

**Figure 6:**
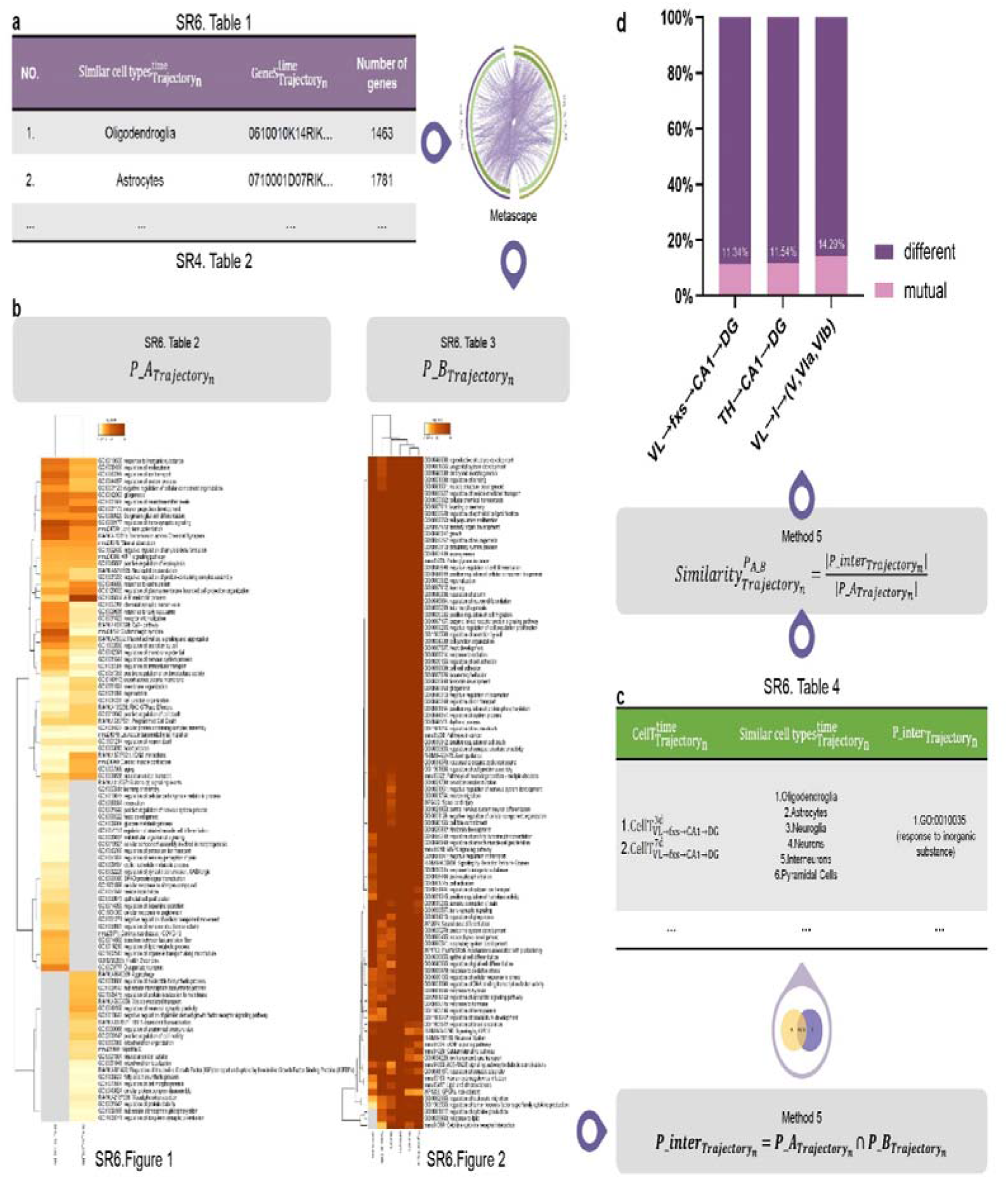
Pathway analysis. **a,** The marker genes of the similar cell type. **b,** The mutual pathway sets for our identified cell subtypes *P_A_Trajectory_n__* and similar cell types *P_B_Trajectory_n__*. **c,** The mutual pathway set *P_inter_Trajectory_n__* between our identified cell subtypes *P_A_Trajectory_n__* and similar cell types *P_B_Trajectory_n__*. **d,** The similarity between *P_A_Trajectory_n__* and *P_B_Trajectory_n__*.

After that, using SR6. Table 1 and marker genes of our identified cell subtypes (SR4. Table 2) as the input, we employed Metascape^10^ to carry out pathway analysis for each of them. Figure 6b shows mutual pathway sets for our identified cell subtypes *P_A_Trajectory_n__* (SR6. Table 2 and SR6. Figure 1) and similar cell types *P_B_Trajectory_n__* (SR6. Table 3 and SR6. Figure 2).

Figure 6c and SR6. Table 4 show the mutual pathway set *P_inter_Trajectory_n__* between our identified cell subtypes *P_A_Trajectory_n__* and similar cell types *P_Bτrajectory_n_*, the process of which is detailed in signalling pathway analysis of Methods section.

Fig. 6d shows that our identified cell subtypes and their similar cell types on *VL* → *fxs* → *CA1* → *DG, TH* → *CA1* → *DG* and *VL* → *I* → (*V*, *VIa*, *VIb*) have 11, 9 and 9 mutual pathways, respectively. Since they have mutual pathways (SR6. Table 4), and the proportion of these mutual pathways remains stable for each trajectory, we hypothesized that our identified cell subtypes and their similar cell types found at the cell level from CellMeSH should be the same at the molecular level.

## 3. Discussion

In the present study, we employed an intraventricular haemorrhage mouse model with haematoma in the lateral ventricle as primary stimulation for the ipsilateral hemisphere. Then, we developed spatial transcriptome sequencing-based bioinformatic algorithms to investigate the 3D spatial brain regions affected by the primary stimulation of haematoma in the ipsilateral ventricle after IVH (Figure 1). Based on this, we have the following important findings: (1) we found three 3D global pseudospace-time trajectories (Figure 2); (2) we not only identified the cell subtypes on these trajectories at different times after IVH (Figure 3) but we also revealed a similar pattern for these cell types on each 3D global pseudospace-time trajectory (Figure 4); (3) we observed that the cell-cell communication strength greatly changes after IVH and returns to the normal state later by analysing the cell-cell communication strength (Figure 5); and (4) our identified cell subtypes and their similar cell types found from the single-cell sequencing database (CellMeSH ^27^) should be similar as each other at the molecular level (Figure 6).

As SM. Figure 1 indicates, 25 μl of autologous blood was injected into the contralateral ventricle to establish the IVH model used in this study, which exhibited obvious haematoma drainage across the interventricular foramen and deposition in the ipsilateral ventricle but without direct damage to the ependyma barrier in the ventricles. We believe this volume of haematoma is the ideal model to investigate the secondary neurological dysfunction derived by haematoma stimulation after IVH. The 3D global pseudospace-time trajectories (Figure 2. e, f and g) indicated that haematoma stimulation in the lateral ventricle mainly affected the ipsilateral cortex and hippocampus. For the ipsilateral cortex, recent brain functional connectivity analysis indicated that IVH in infants born moderately preterm and later preterm was associated with the frontoparietal operculum and orbitofrontal cortex, which are related to language and cognition during development^30^. In addition, low-grade IVH after germinal matrix haemorrhage in preterm neonates was reported to cause lower cerebral blood flow in posterior cortical and subcortical grey matter regions^31^, all within the trajectory bundle III directly from the lateral ventricle to the ipsilateral cortex, suggesting regional vulnerability of these brain structures. However, intraventricular extension after intracerebral haemorrhage might not be associated with dysphagia, even with a space-occupying effect and midline shift^32^. As illustrated in Figure 4c and SR4. Table 3, we revealed a pattern in which not only are the cell subtypes after IVH similar for *VL* → *fxs* → *CA1* → *DG*, TH → CA1 → DG and VL → I → (V, VIa, VIb), but also these cell subtypes are different from the cell subtypes of the sham group (with only the acupuncture effect). However, SR4. Table 3 demonstrates that the pattern is not as obvious for VL → I → (V, VIa, VIb) compared to *VL* → *fxs* → *CA1* → *DG* and TH → CA1 → DG. We explain this phenomenon as follows: (1) motor dysfunction usually occurs in most clinical patients with isolated intraventricular haemorrhage; (2) our data show that lateral ventricle stimulation has only a transient effect on the ipsilateral cortex.

However, since most patients with intraventricular haemorrhage are in a supine resting state after onset, there is no clinical evidence to accurately describe the severity of motor dysfunction in its acute phase, and the effects of stimulation largely returned to normal levels after 7 days of intracerebral haemorrhage.

Relative to the limited evidence of haematoma stimulation in the lateral ventricle for the cortex, we also illustrated two major 3D global pseudospace-time trajectory bundles for the hippocampus. Trajectory bundle I (SR2. Figure 3 b, e, h) is a direct neuronal circuit from the lateral ventricle to the hippocampus, which in our understanding is closely associated with white matter lesions after intraventricular extension in spontaneous intracerebral haemorrhage patients^33^. Trajectory bundle II (SR2. Figure 3 c, f, i) is a novel integrated signalling circuit from the hypothalamus to the hippocampus that flexibly modulates long-term potentiation^34^ and selected components of cognition^35^. In addition, corpus callosum injury is also reported to strongly correlate with the severity of IVH^36^ and serves as a prognostic factor for poor outcomes after brain trauma. Consistent with our previous rodent experiment, perihaematomal tissue injury and neurocognitive deficits are reported in intracerebral haemorrhage with ventricular extension^37^, and blood metabolites such as iron^38^, oxyhemoglobin^39^ and thrombin^39,40,^ are well considered in this pathophysiological process. Combined with the negative results of the CLEAR III trial, our data in Figure 4c indicate that blood metabolism rapidly stimulates the direct neuronal circuit from the lateral ventricle to the hippocampus, and clearing the blood clots in the ventricle might not successfully remove these stimulations in the cerebrospinal fluid. Subsequent secondary brain injury due to haematoma stimulation in the ventricles might be a promising target and therapeutic strategy for IVH patients. Figure 4d and SR4. Table 5 illustrate the most affected or motivated cell types by our identified cell subtypes (Figure 4b and SR4. Table 2), which could vividly refer to the potential pathophysiological changes in these main trajectories/neural circuits after IVH.

For VL→fxs→CA1→DG from trajectory bundle I, which might be tightly associated with white matter lesions, as we discussed above, the most motivated cell type is oligodendroglia, consistent with it being widely understood as the main participator in white matter and cognitive functions. Further pathway analysis indicated intense metabolic changes and synaptic activity, as well as gliogenesis and neural cell death in this trajectory. The cell-cell communication strength analysis found a stimulation after IVH which significantly turned into inhibition, as shown in Figure 5d. During the post-IVH period, these 11 mutual pathways (Figure 6d) reflect the core pathophysiological changes, which might be potential clues and therapeutic targets for further preclinical study.

Additionally, in TH→CA1→DG from trajectory bundle II, it seems that neuronal cells are the main participators, and cell population proliferation is the highest changing pathway, which is supported by a recent study^41^ that hypothalamic circuits could modulate adult hippocampal neurogenesis for memory regulation.

However, the pathway analysis in VL→G→(V,VIa,VIb) from trajectory bundle III indicates an acute response to hypoxia and oxidative stress, which is consistent with the significant inhibition in cell-cell communication strength on this trajectory on the 3^rd^ day after IVH(Figure 5d). Nevertheless, the communication strength recovered to a normal level on the 7^th^ day after IVH, which might be an explanation for the short-term motor dysfunction of IVH patients for clinical attention in the future.

Our proposed bioinformatics analysis workflow has the following innovations and limitations. First, we developed a 3D global pseudospace-time trajectory reconstruction algorithm, which can not only investigate the genetic changes in 3D global pseudospace-time trajectories but also identify cell subtypes on these trajectories at different times. However, due to the limited slices and the conflict between planner trajectories, this algorithm cannot obtain high accuracy.

Second, since our proposed similarity algorithm considers the up/down regulation for marker genes of cell subtypes, we can accurately compute the similarity between cell subtypes at different times on the same trajectory. However, since spatial transcriptome data only provide discrete up/downregulation data, the accuracy of our similarity algorithm is limited.

Third, our proposed cell-cell communication intensity density algorithm can uniformly measure cell-cell communication intensity for multiple spatial transcriptome slices at different times for each ligand paired with cell subtype compared to a previous method^9^ that analysed cell-cell communication intensity for a single slice. However, since we do not have an automatic algorithm to match cell subtype ligand receptors in the CellTalkDB database^28^, it is time-consuming for us to process such cell subtypes that have many paired ligands.

In conclusion, we innovated a bioinformatics algorithm to discover three main 3D global pseudospace-time trajectory bundles representing the main neural circuits from the lateral ventricle to the hippocampus and primary cortex induced by experimental intraventricular haematoma stimulation. Further analysis indicated a rapid response in the primary cortex, as well as a direct and integrated effect on the hippocampus after IVH. To the best of our knowledge, this is the first study to investigate secondary brain injury after IVH by using spatial transcriptome sequencing and bioinformatics analysis. These data could provide more helpful information for elucidating the pathophysiological mechanism of IVH in patients after acute brain injury, as well as the analysis of algorithm strategies for similar studies in the future.

## Supporting information

Supplementary of Results and Supplementary of Methods

## Acknowledgements

This work was supported by the State Key Laboratory of Trauma, Burn and Combined Injury [No. SKLYQ202002 to Yujie Chen], National Key Research and Development Program of China [Nos. 2021YFF1201200 and 2018ZX10201002 to Le Zhang], National Natural Science Foundation of China [No. 82030036 to Hua Feng], Sichuan Science and Technology Program [No. 2022YFS0048 to Le Zhang].

## Author contributions

Le Zhang and Yujie Chen conceptualized and designed experiments and supervised the research. Xufang Ru, Suna Huang and Yujie Chen performed animal experiments. Le Zhang, Jiayidaer Badai, Guan Wang, Wenkai Song, Yujie You, Jiaojiao He and Yujie Chen acquired and analyzed the data. Le Zhang, Jiayidaer Badai, Guan Wang, Wenkai Song, Hua Feng, Runsheng Chen, Yi Zhao, and Yujie Chen interpretated the data. Le Zhang, Jiayidaer Badai, Wenkai Song, Yi Zhao and Yujie Chen drafted the manuscript. All authors have read and approved the current version of the manuscript.

## Data availability

All data generated or analysed during this study are included in this manuscript and its supplementary information files. The raw sequence data reported in this paper have been deposited at Gene Expression Omnibus (https://www.ncbi.nlm.nih.gov/geo/, No. GSE214349).

## Methods

### 1. Experimental setup

Five wild-type C57/BL6 male mice (weighing 25±5 g, 8 weeks old, Experimental Animal Center of Third Military Medical University, Chongqing, China) were used in this study. These mice were housed in a temperature-controlled room under specific-pathogen-free conditions and a standard 12-h light/dark cycle, with ad libitum access to food and water. All experiments were reported in compliance with the Animal Research: Reporting in vivo Experiments (ARRIVE) guidelines. The experimental protocols were approved by the Laboratory Animal Welfare and Ethics Committee of Third Military Medical University (AMUWEC2020762) and performed according to the Guide for the Care and Use of Laboratory Animals.

An intraventricular haemorrhage model was established according to previously described methods with modified coordinates. Briefly, mice in each group were deeply anaesthetized by intraperitoneal injection of 2 mg ketamine and 0.4 mg xylazine in 0.9% saline. A feedback-controlled heating pad was used to maintain body temperature at 37.0 °C. A small cranial burr hole was drilled, and a 32-gauge needle was inserted stereotaxically into the right lateral ventricle (coordinates: 1.0 mm lateral, 0.24 mm posterior, and 2.5 mm ventral to bregma) to establish the intraventricular haemorrhage model under stereotactic guidance. Then, 25 μl of autologous blood was injected at a rate of 5 μl/min using a microinfusion pump (Harvard Apparatus, Holliston, MA). The burr hole was sealed with bone wax, and the skin incision was closed with sutures after the needle was removed. Sham-operated mice underwent the same surgery without blood injection. The brains of the mice were harvested at different time points after IVH after deep anaesthesia for further study.

### 2. Visium spatial transcriptome sequencing

We collected wild-type C57/BL6 mouse brains, which were classified into the no-surgery group (control), the needle-puncture group without blood injection (sham), and the needle-puncture group with blood injection at 1^st^, 3^rd^ and 7^th^ day. Then, these brains were frozen on dry ice in OCT.

Frozen brains were then sectioned coronally at 10 μm on a cryostat and mounted on 10x Genomics Visium spatial transcriptome slides. The tissue locations of the library patches started from the emergence of the lateral ventricle with 20 μm intervals and a 10 μm margin of error (SM. Table 1).

For morphological analysis and spatial alignment of sequencing data, sections were fixed in methanol at −20 °C for 30 min before H&E staining. Additionally, all fractions were measured using a bioanalyzer (Agilent 2100) meeting the RNA integrity number RIN>=7.

After brightfield imaging, we permeabilized 19 brain slices, the permeabilization time of which is listed in SM. Table 1. After tissue permeabilization, we captured poly-A mRNA at each point in the capture region and added spatial barcodes and unique molecular identities into reads for library construction.

Our library detection instrument employs a DNF-915 kit and QPCR for quantification. After the library was qualified, we performed PE150 paired-end sequencing using the Illumina NovaSeq 6000 S4 sequencing platform with 50,000-100,000 reads per locus (spot).

Finally, genomes and images were aligned by mouse reference genome mm10-3.0.0 and SpaceRanger (v1.1.0), respectively. The data were analysed and normalized by using Seurat (v3.1.2) and SCTransform, respectively.

### 3. Cell subtype analysis

#### 3.1 Cell subtype identification

We used the top 30 upregulated/downregulated 3D transition genes for each 3D subtrajectory (SR3. Table 2) as input. Then, we employed our developed algorithm (SM. Tables 4 & 5) to identify a cell subtype corresponding to the selected trajectory at different times (SR4. Table 1).

#### 3.2 Similarity algorithm for cell subtypes

We used the marker gene sets of cell subtypes corresponding to the selected trajectory at different times (SR4. Table 2) as input. Then, we employed a similarity algorithm for cell subtypes (SM. Tables 4 & 6) to describe the similarity between cell subtypes at different times on the same trajectory (SR4. Table 3).

#### 3.3 Search for similar cell types by CellMeSH

Here, we used marker genes of the cell subtypes as input. Then, we found the top 5 similar cell types for each cell subtype by using CellMeSH^27^. Next, we performed a union operation for these top five similar cell types.

### 4. Cell-cell communication analysis

We use 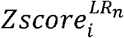 (SR5. Table 2) for each ligand-receptor (*LR_n_*) of cell subtypes and 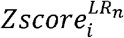 for all the brain regions as the input. Then, we employed the cell-cell communication strength (Density) algorithm (SM. Tables 7 & 8) to obtain the discrete interaction intensity 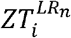 after conversion of the continuous interaction intensity 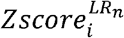 and the sum of the interaction intensity 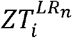 of N points in the corresponding position *ZTTotal* (SR5. Table 3), and the average *Density^LR_n_^* for each ligand receptor of the cell subtypes (SR5. Table 4).

### 5. Signalling pathway analysis

We used mutual pathway sets for our identified cell subtypes *P_A_Trajectory_n__* (*n* = 1…*N*) (SR6. Table 2) and similar cell types *P_B_Trajectory_n__* (*n* = 1…*N*) (SR6. Table 3) as the input. Then, we employed the similarity algorithm for mutual pathway sets (SM. Tables 9 & 10) to obtain the mutual pathway set (*P_inter_Trajectory_n__*) (SR6. Table 4) and the similarity 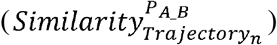 between *P_A_Trajectory_n__* and *P_B_Trajectory_n__* (*n* = 1…*N*) (SR6. Figure 3).

