## Supplementary of Results and Supplementary of Methods for "Discovering Haematoma-Stimulated Circuits for Secondary Brain Injury after Intraventricular Haemorrhage by Spatial Transcriptome Analysis"

This supplementary of results section consists of 6 supplementary files, which is listed as below.

1. **Supplementary of Result 1 (SR1)**

Figure 1 H&E (Hematoxylin and Eosin stain) slices of mouse brain after IVH

a. IVH_C1_50

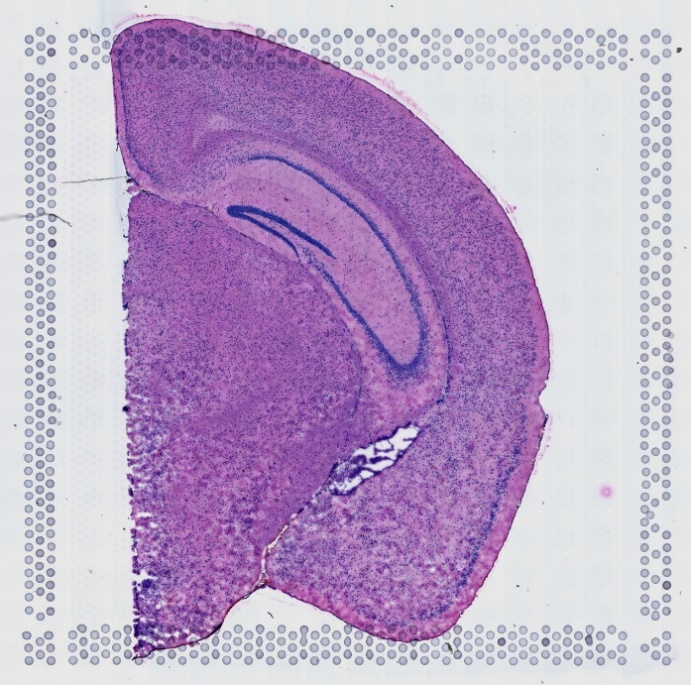

b. IVH_C1_110

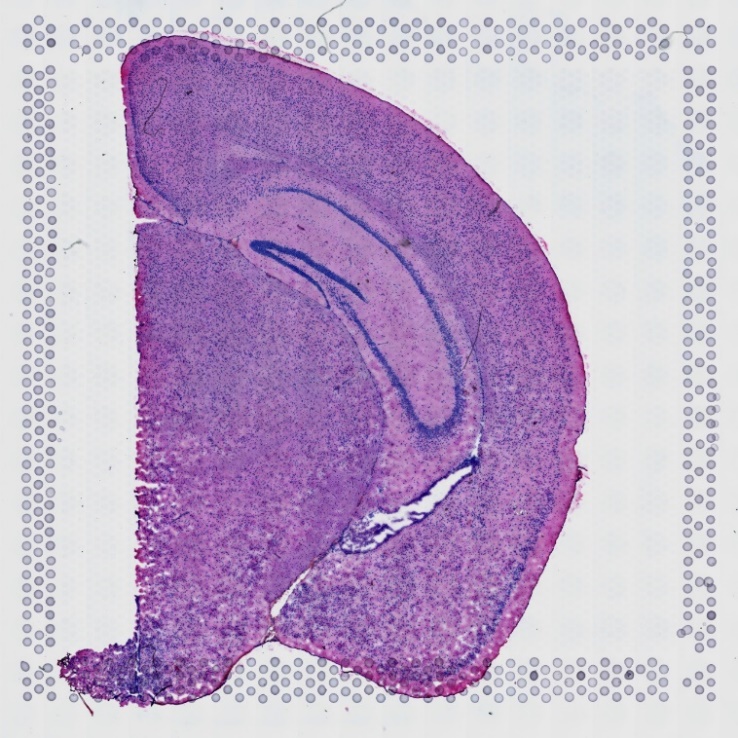

c. IVH_S1_50

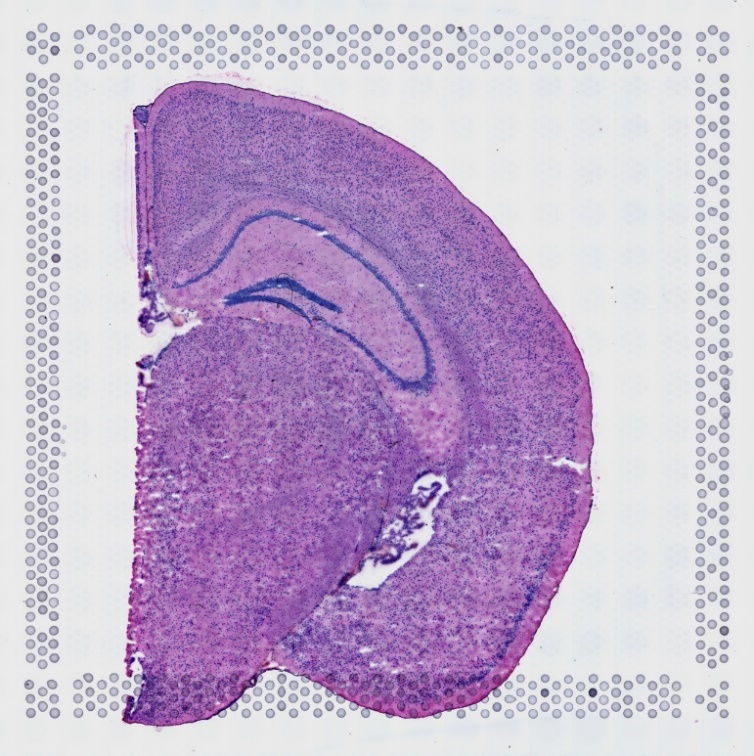

d. IVH_S1_80

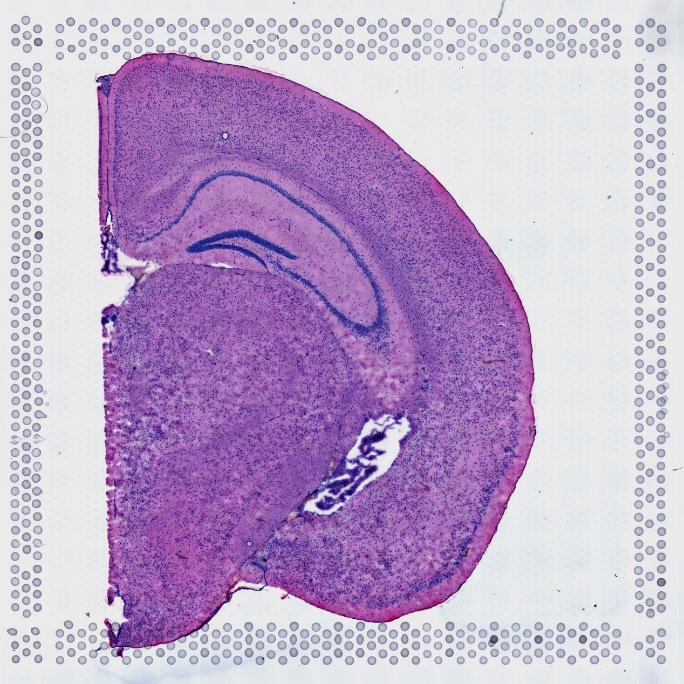

e. IVH_S1_110

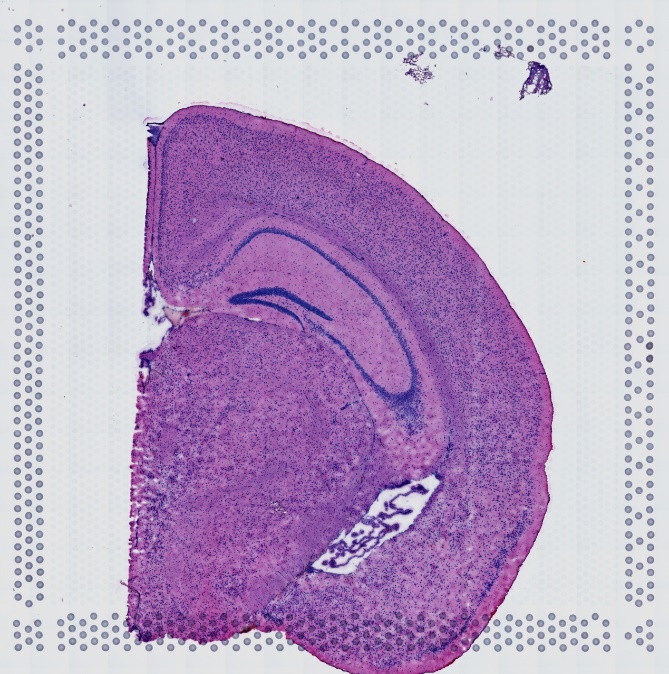

f. IVH_1d_20

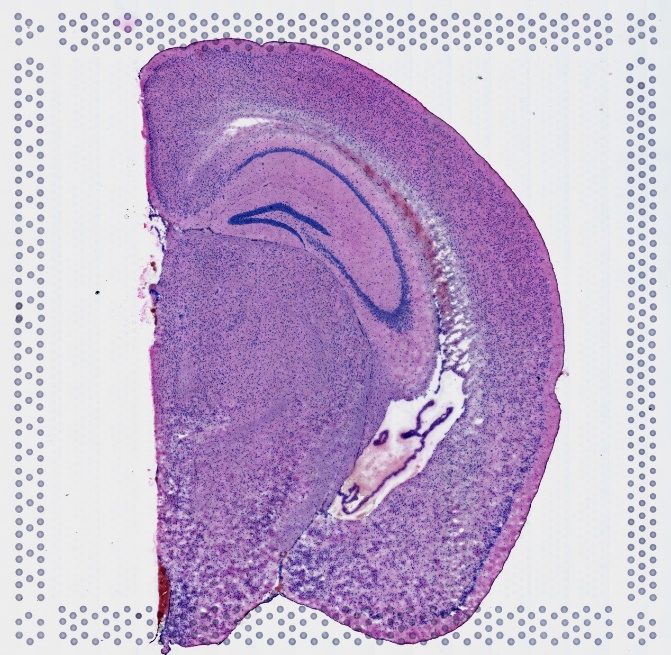

g. IVH_1d_50

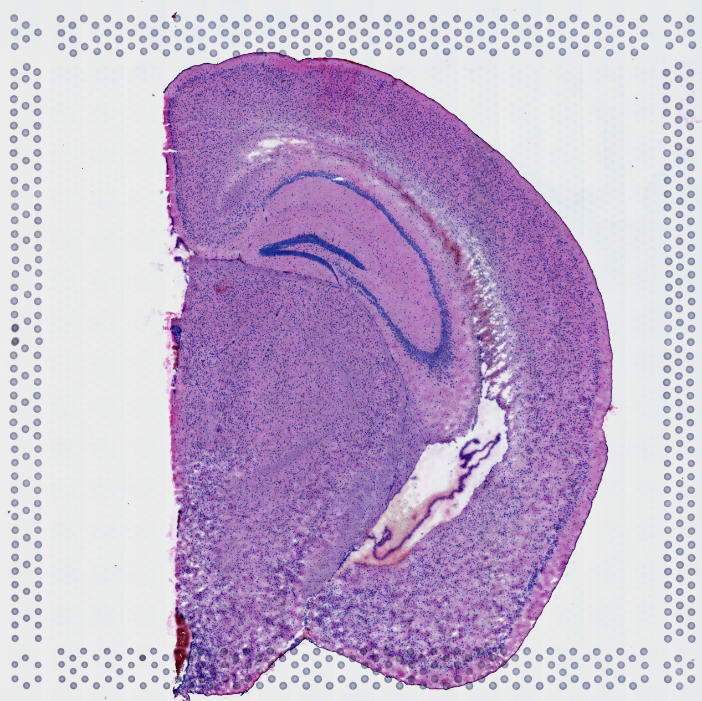

h. IVH_1d_80

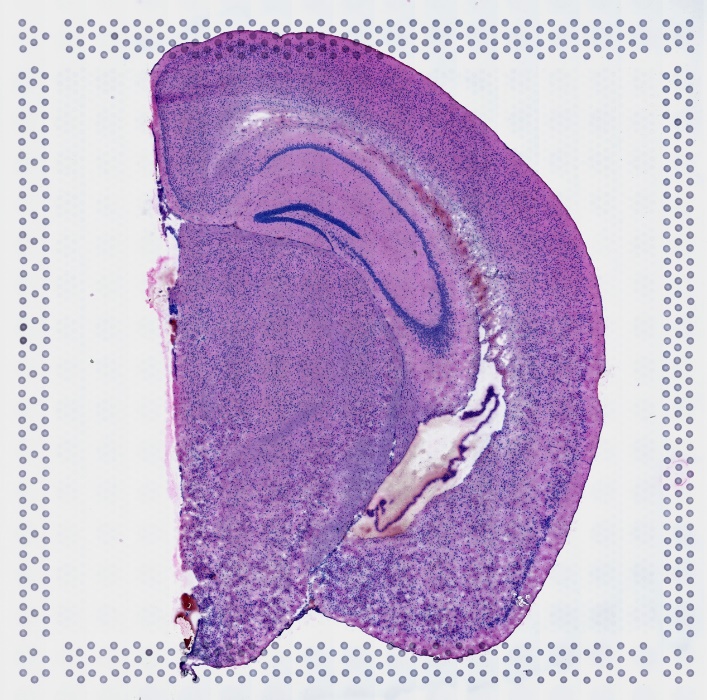

i. IVH_1d_110

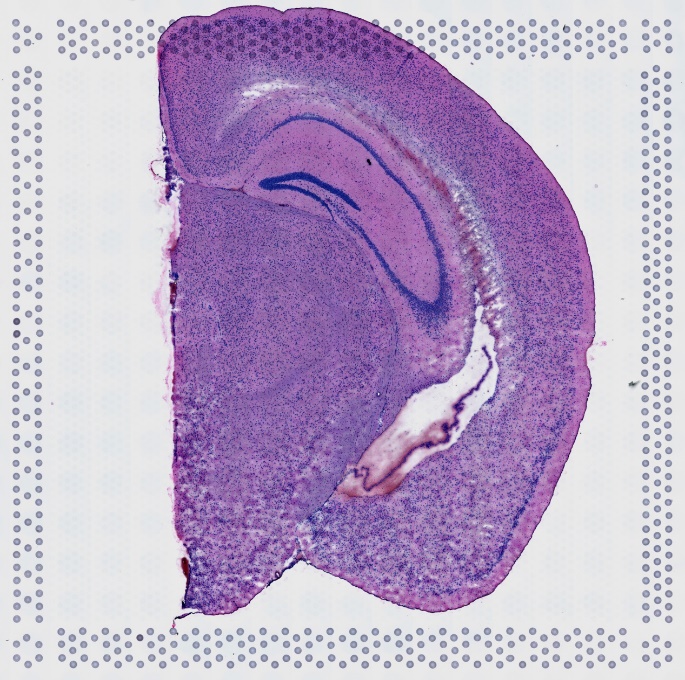

j. IVH_1d_130

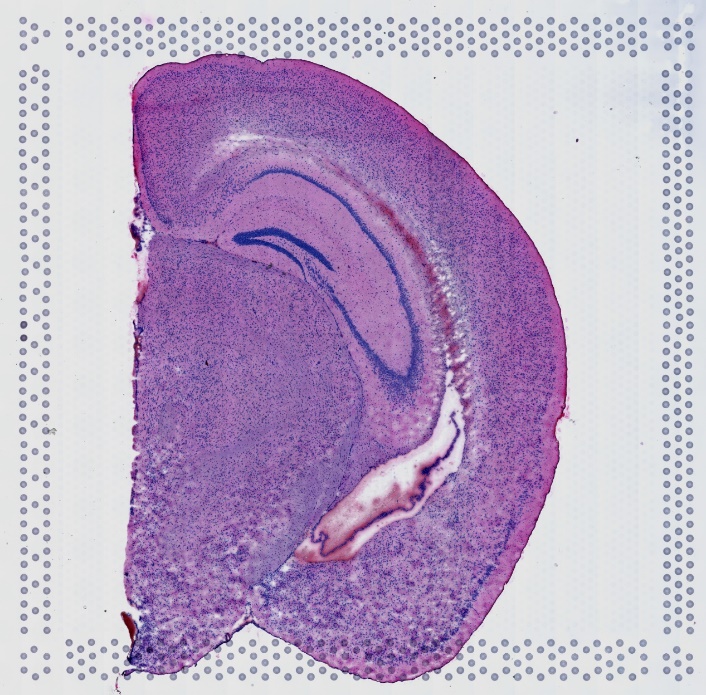

k. IVH_1d_150

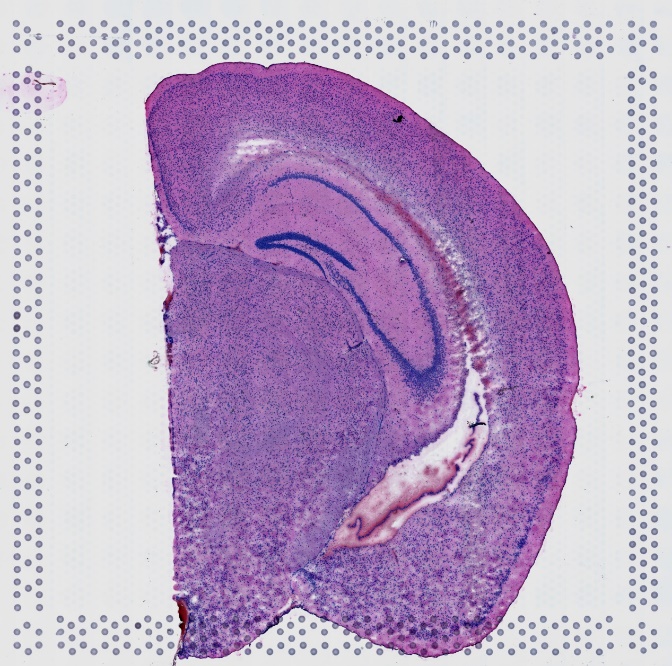

l. IVH_3d_20

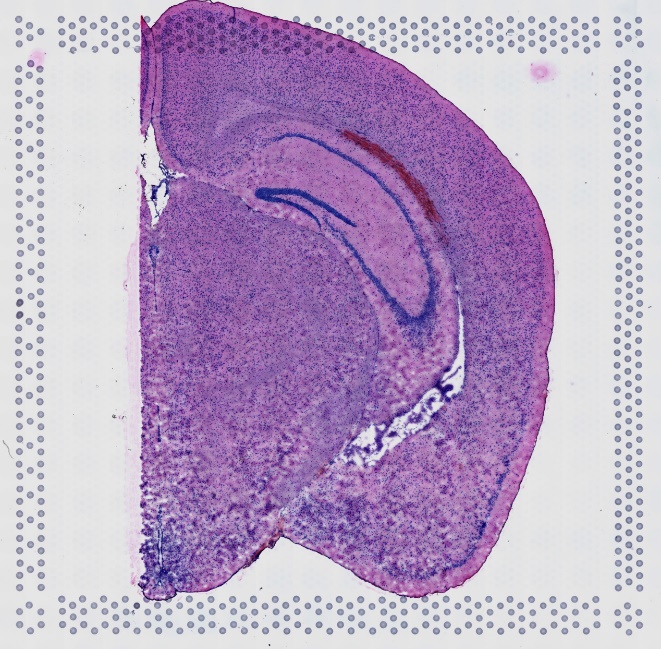

m. IVH_3d_50

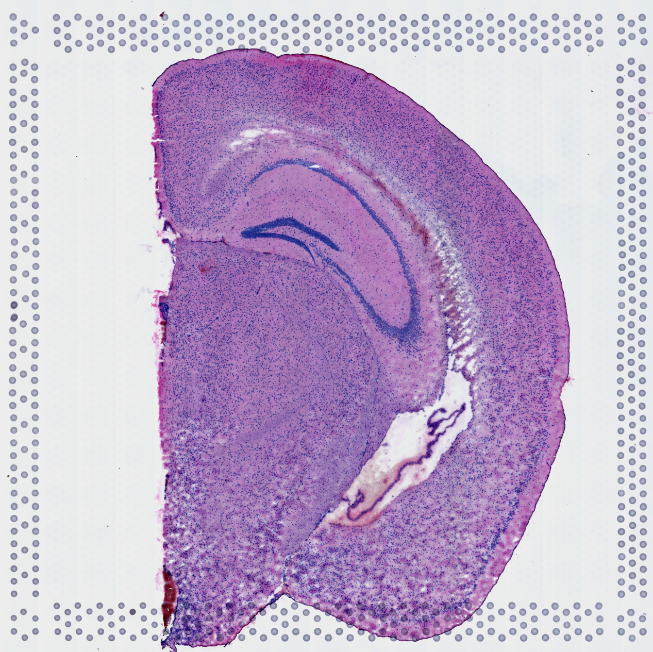

n. IVH_3d_110

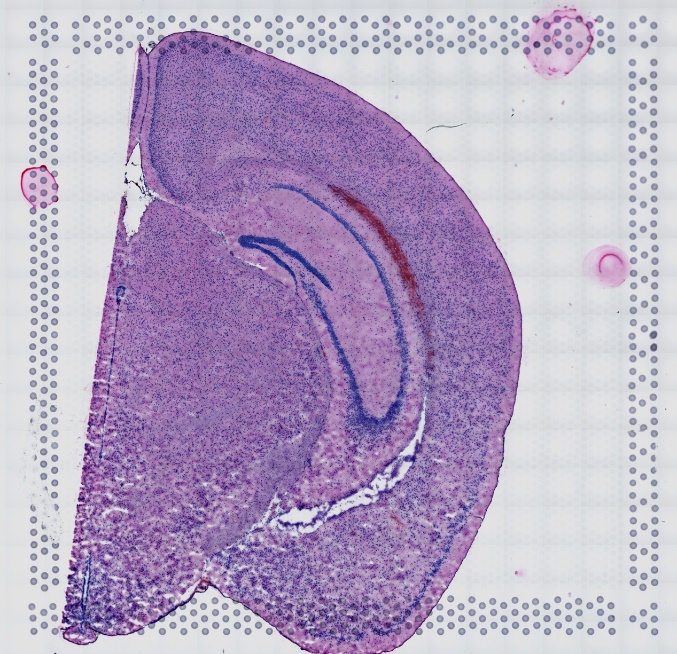

o. IVH_3d_130

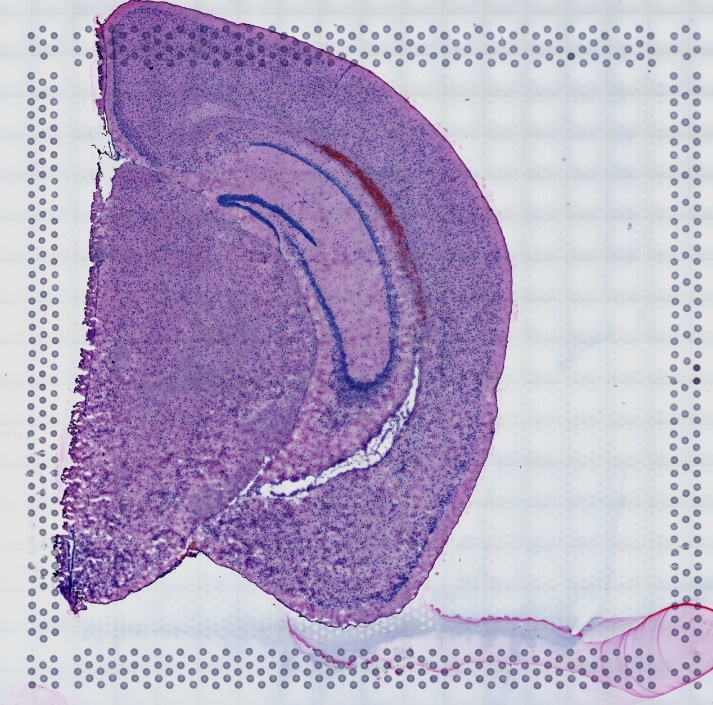

p. IVH_3d_150

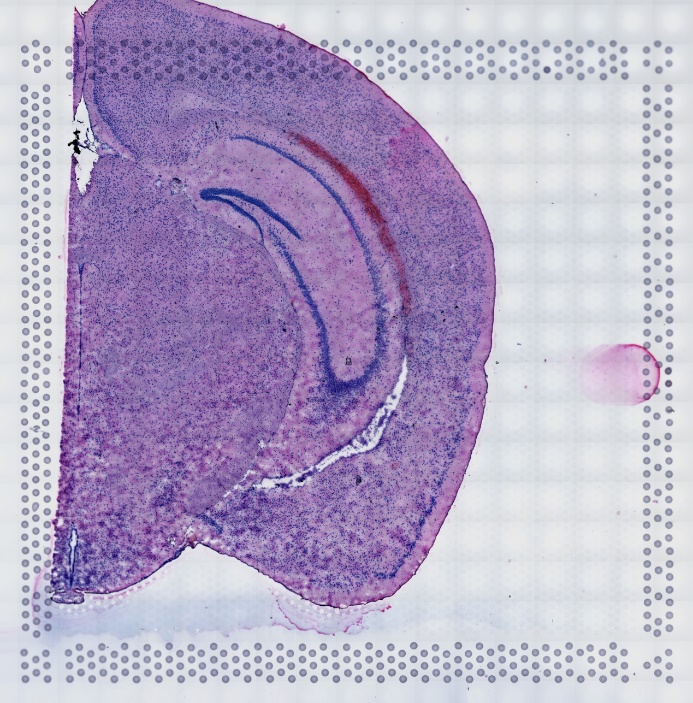

q. IVH_7d_50

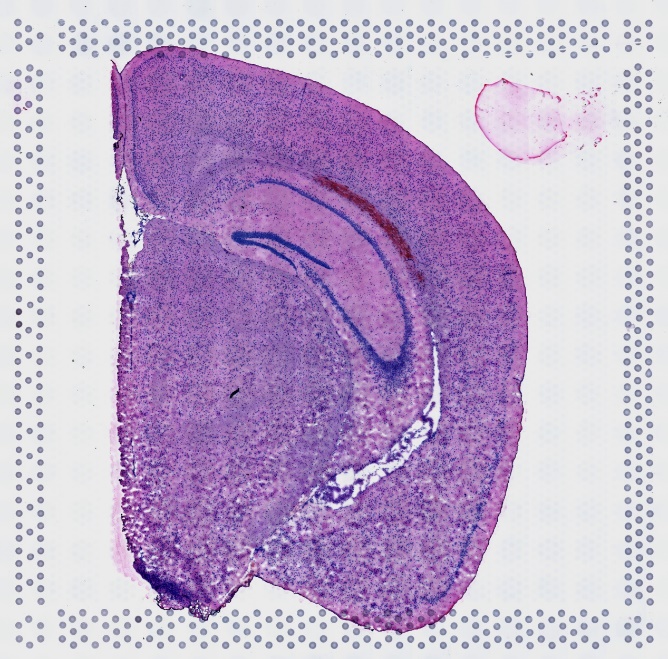

r. IVH_7d_80

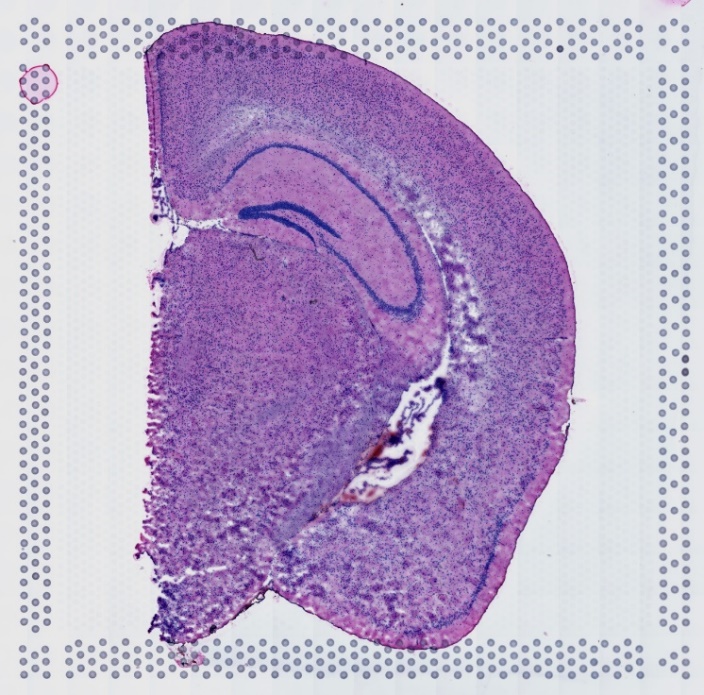

s. IVH_7d_110

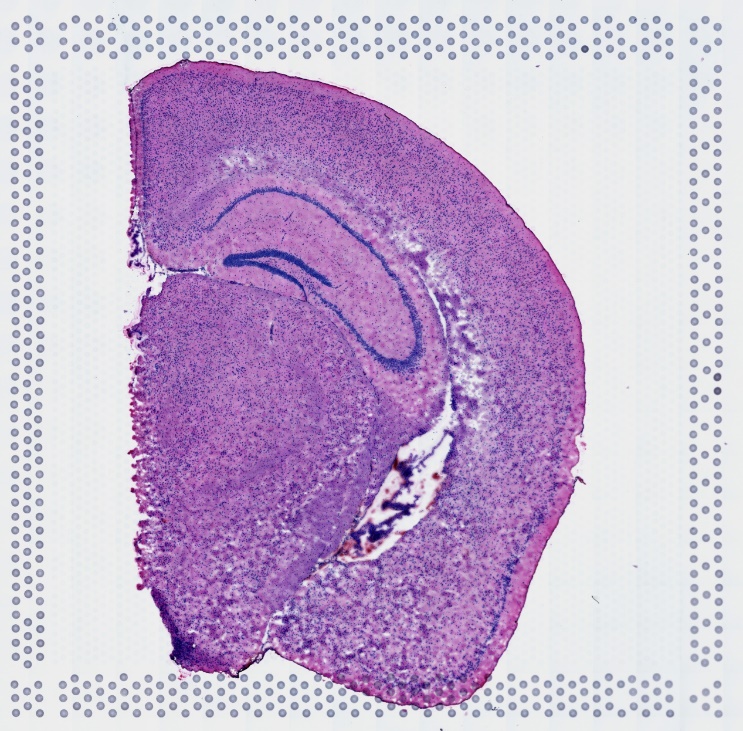

It is noted that a. IVH_C1_50 represents the H&E slices of mouse brain at position 50 for control group.

Figure 2 Cell type clustering for mouse brain

a. IVH_C1_50

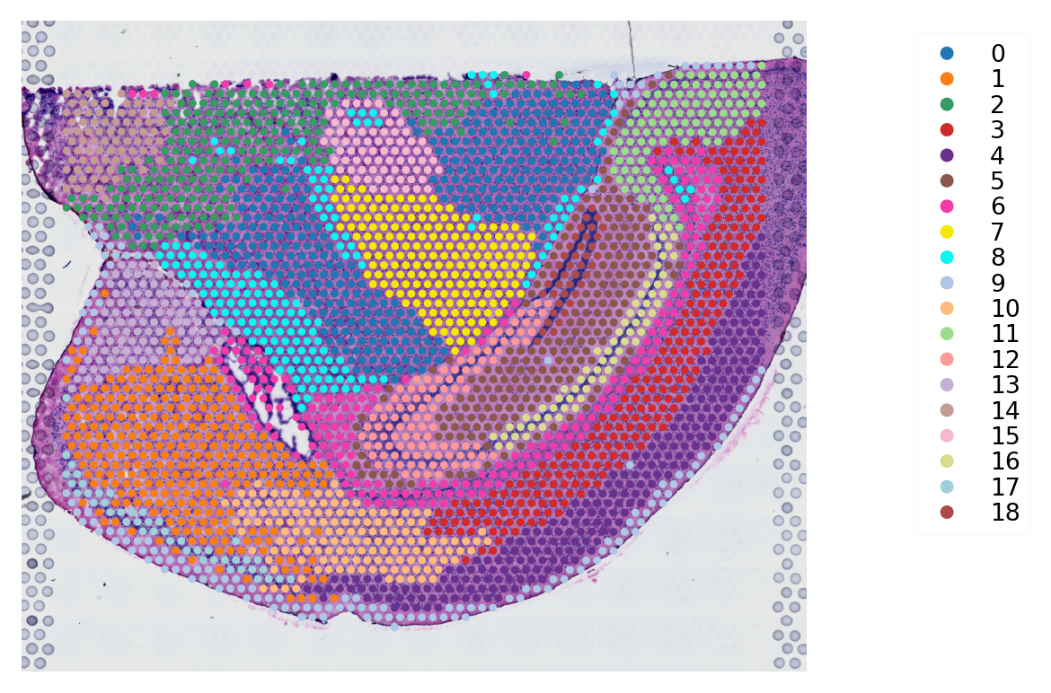

b. IVH_C1_110

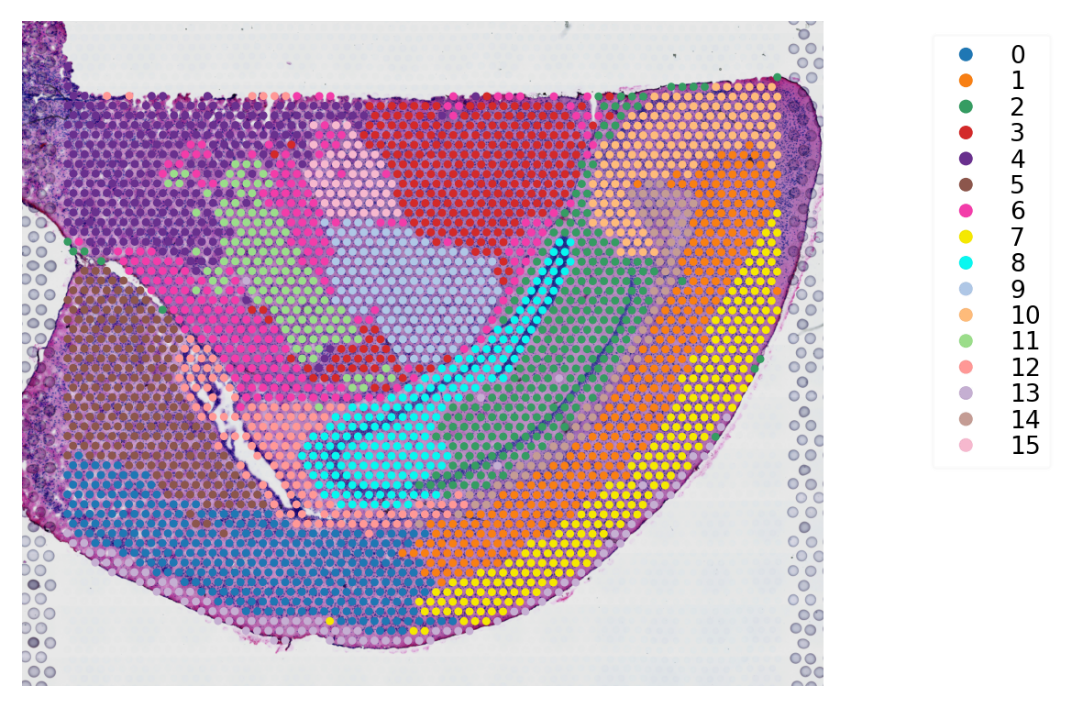

c. IVH_S1_50

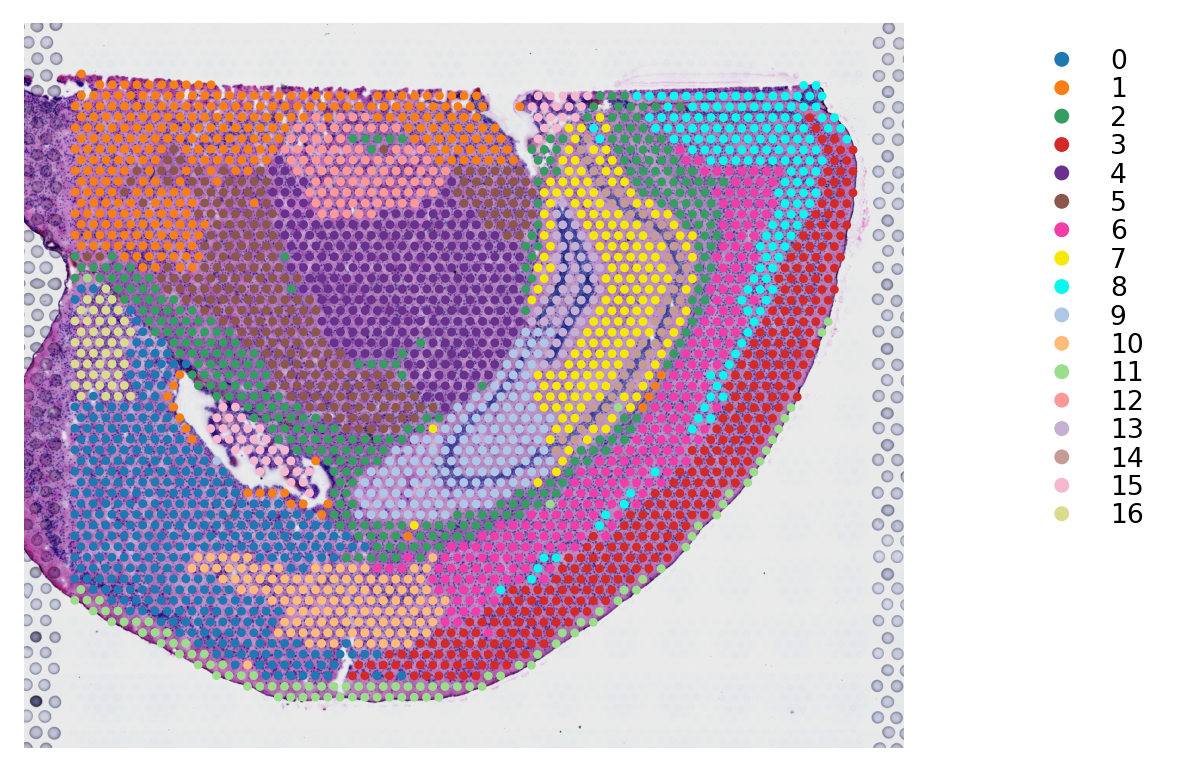

d. IVH_S1_80

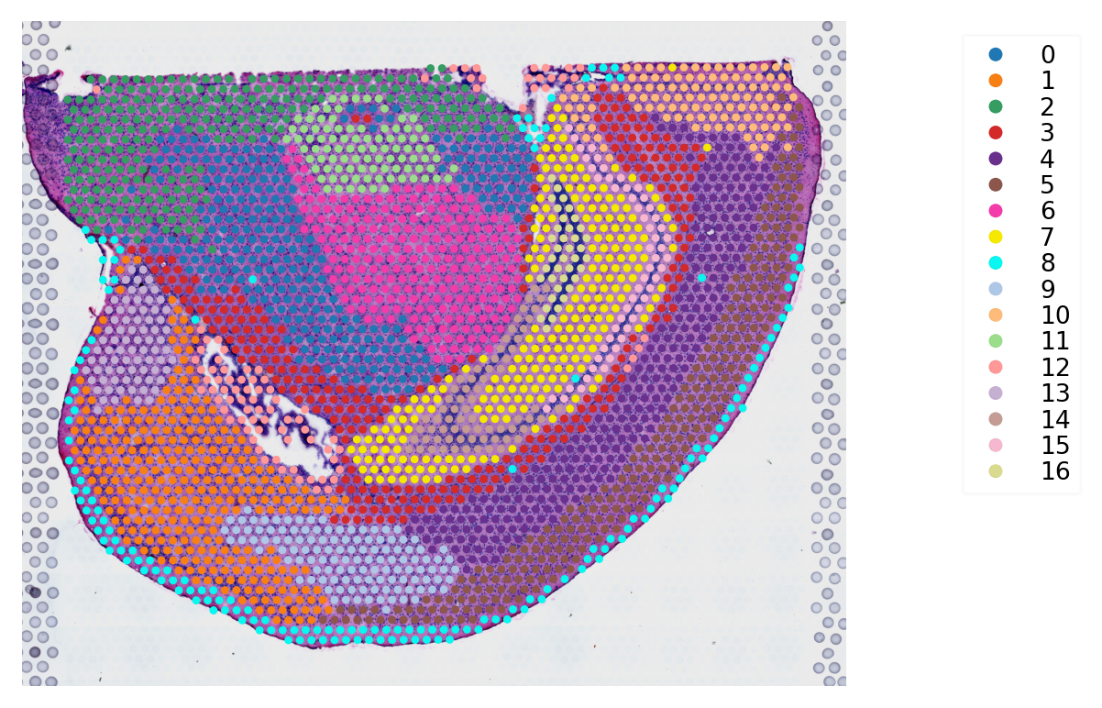

e. IVH_S1_110

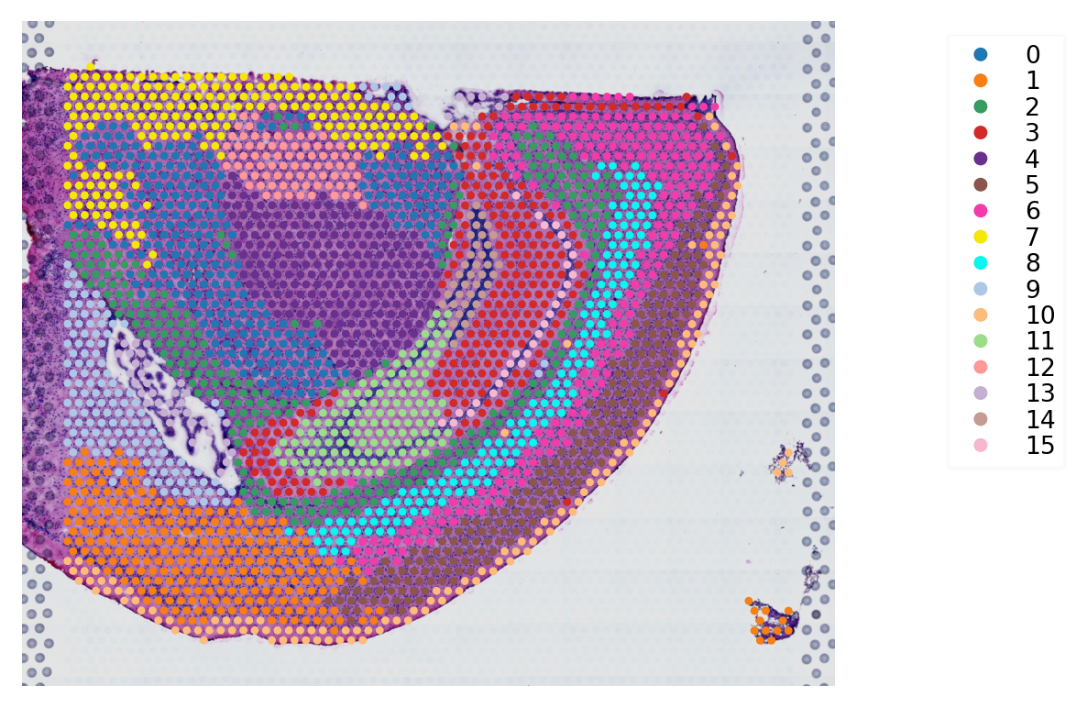

f. IVH_1d_1_20

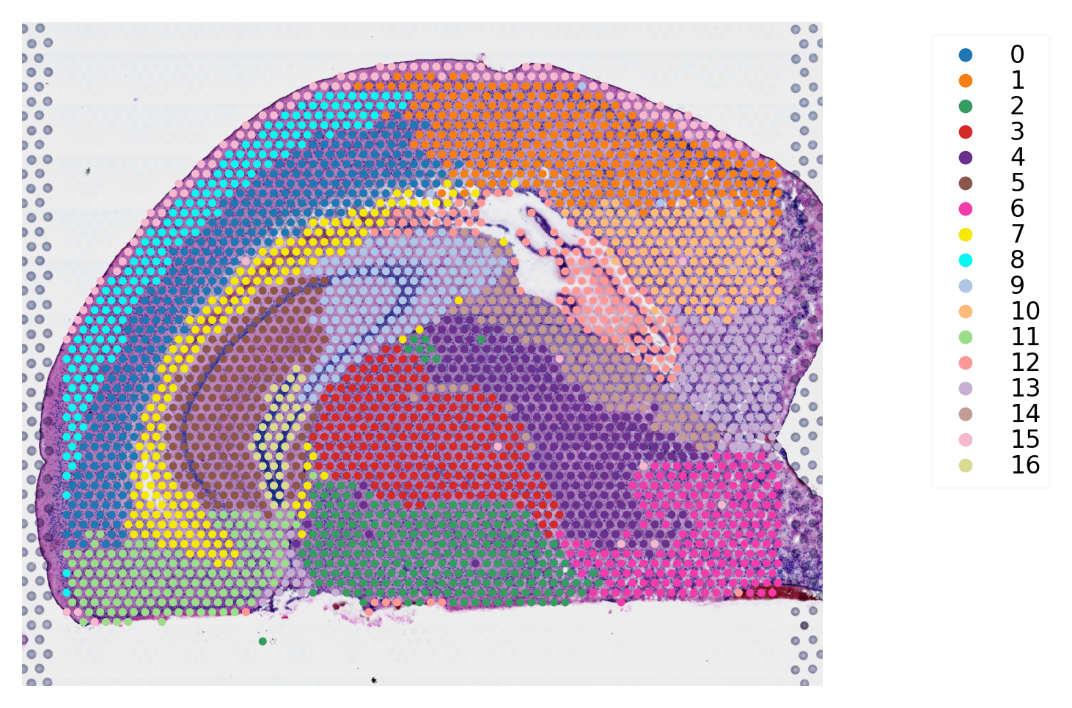

g. IVH_1d_1_50

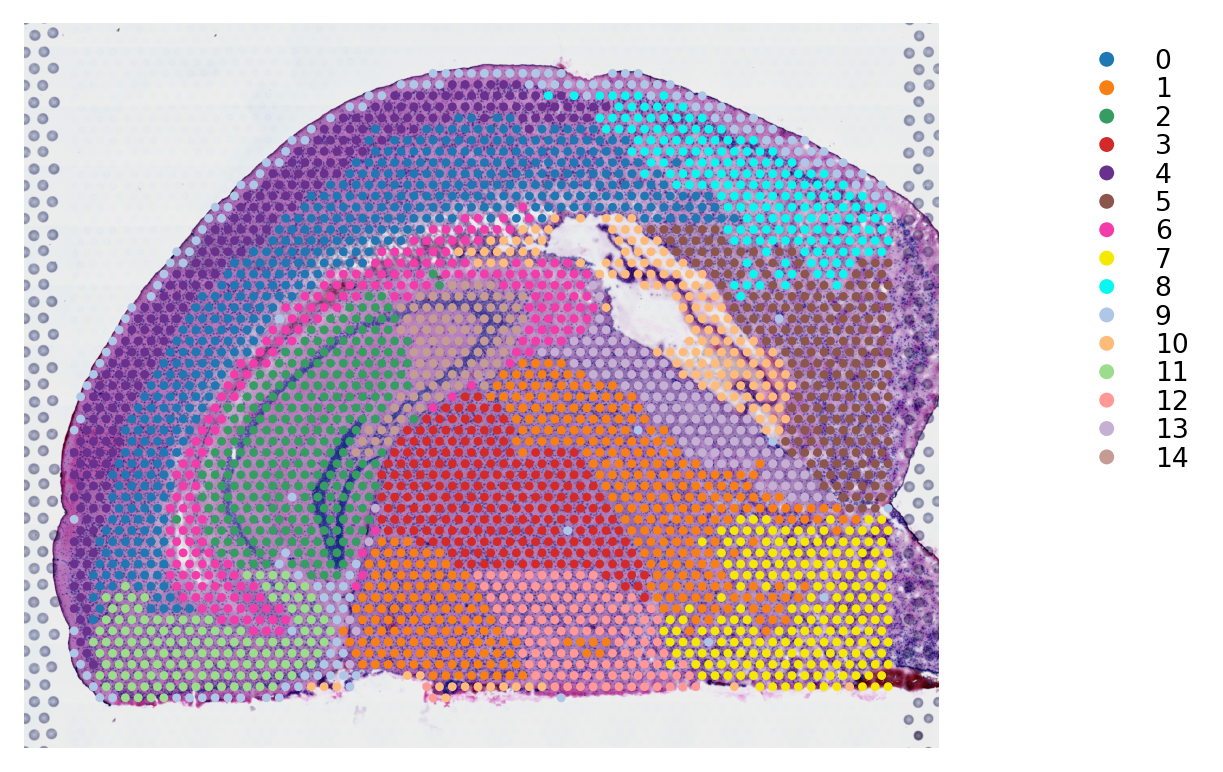

h. IVH_1d_1_80

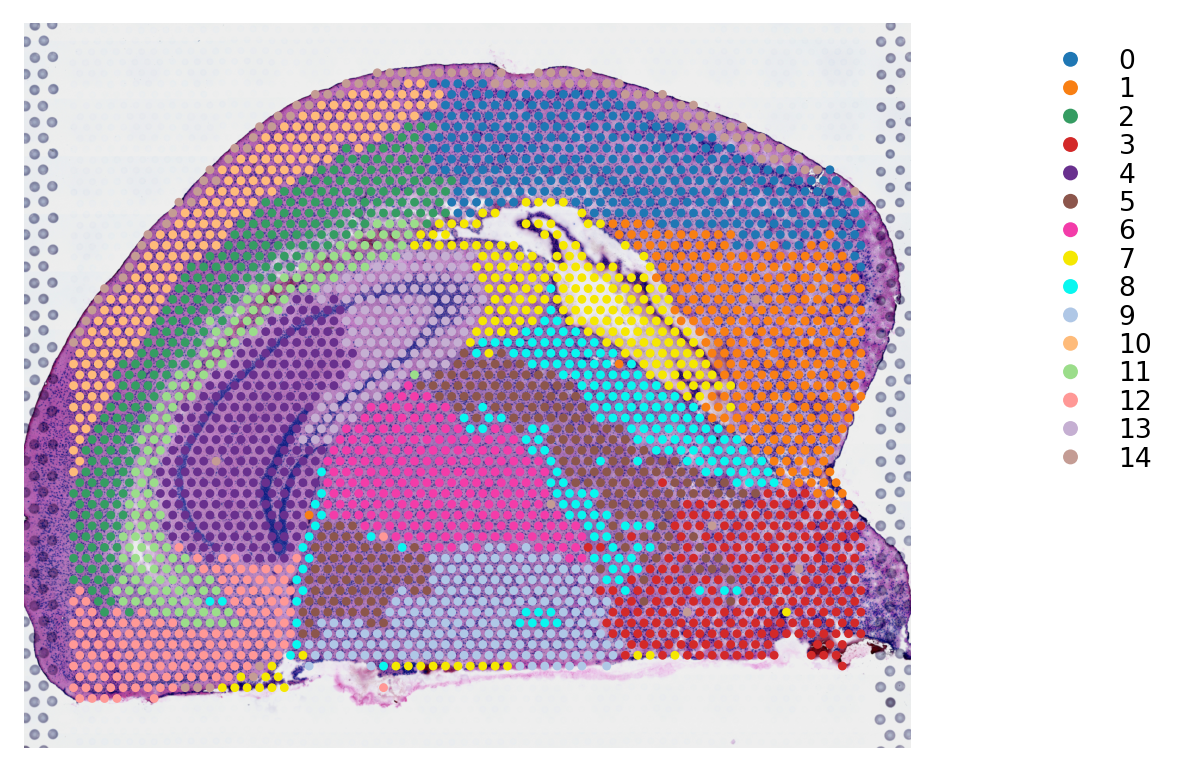

i. IVH_1d_1_110

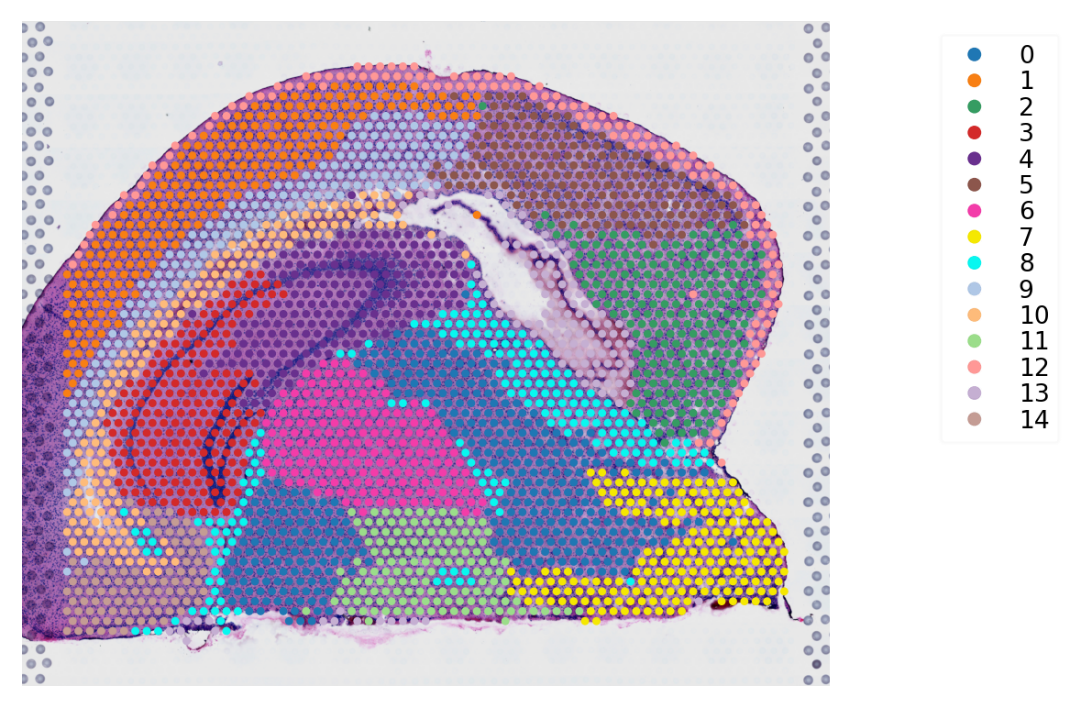

j. IVH_1d_1_130

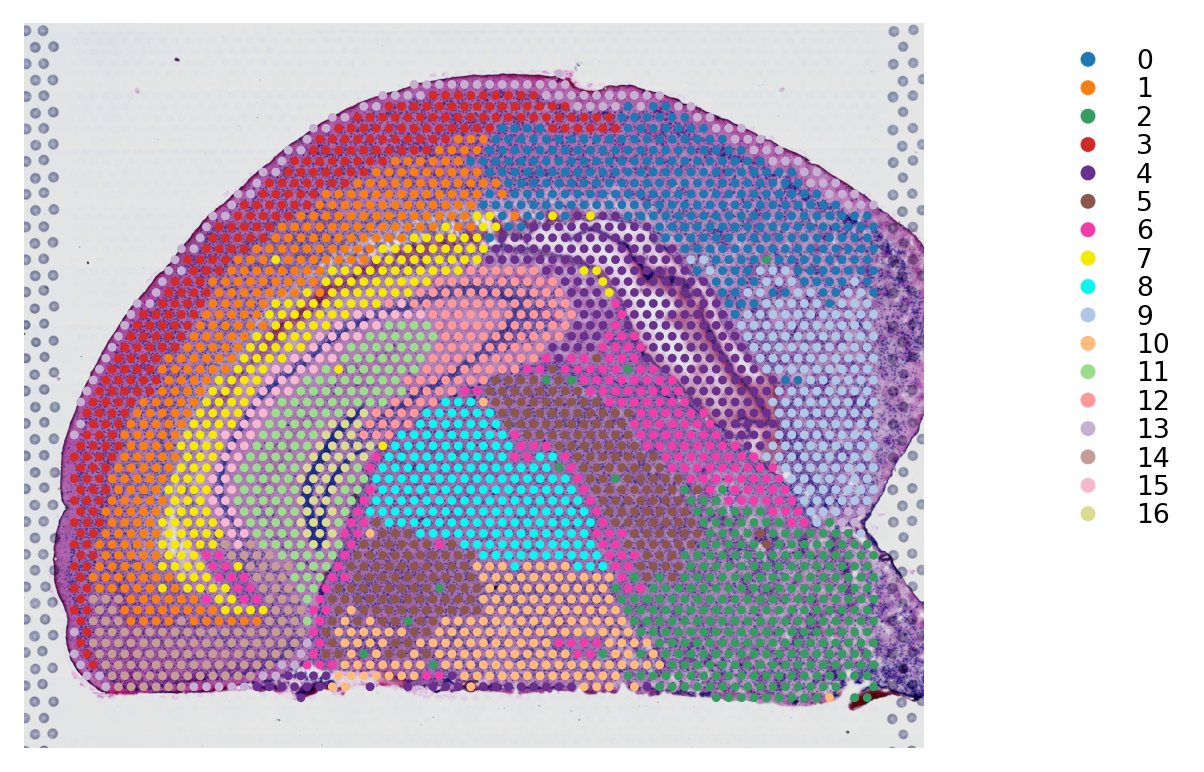

k. IVH_1d_1_150

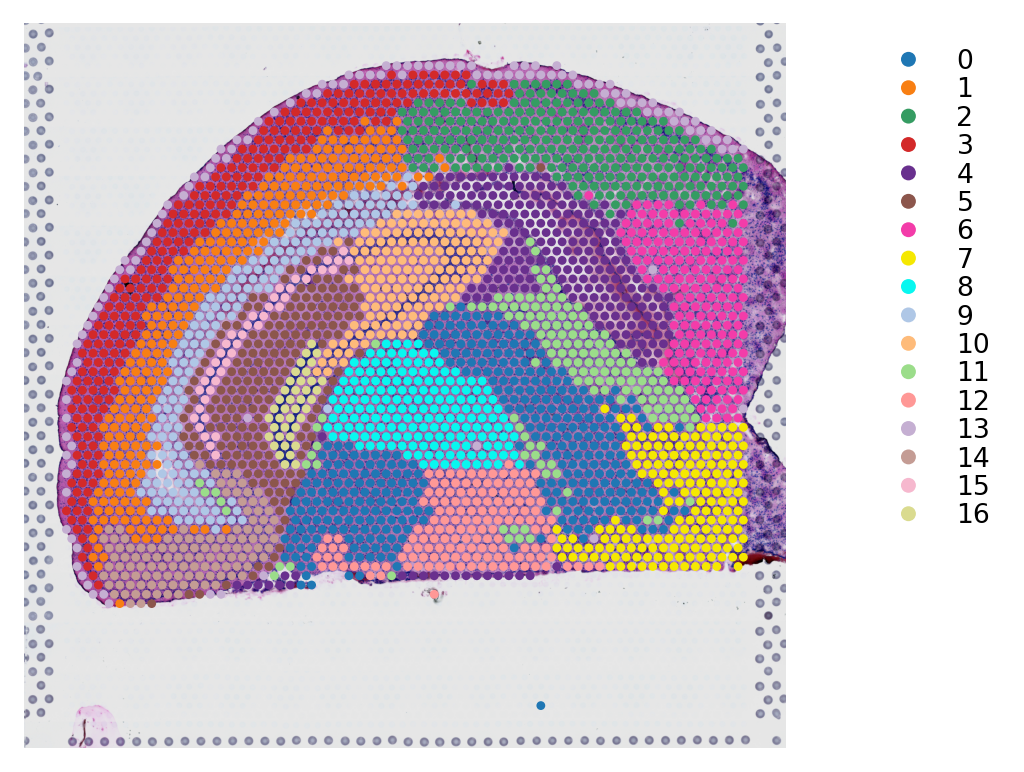

l. IVH_3d_3_20

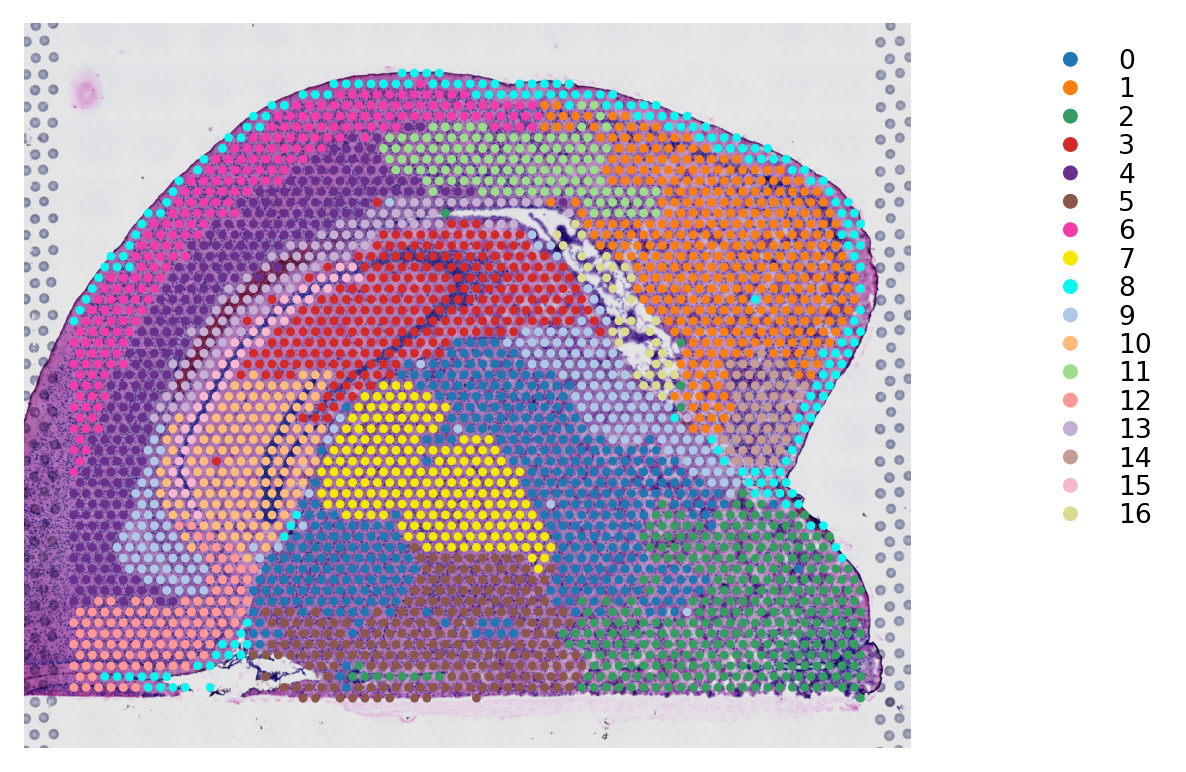

m. IVH_3d_3_50

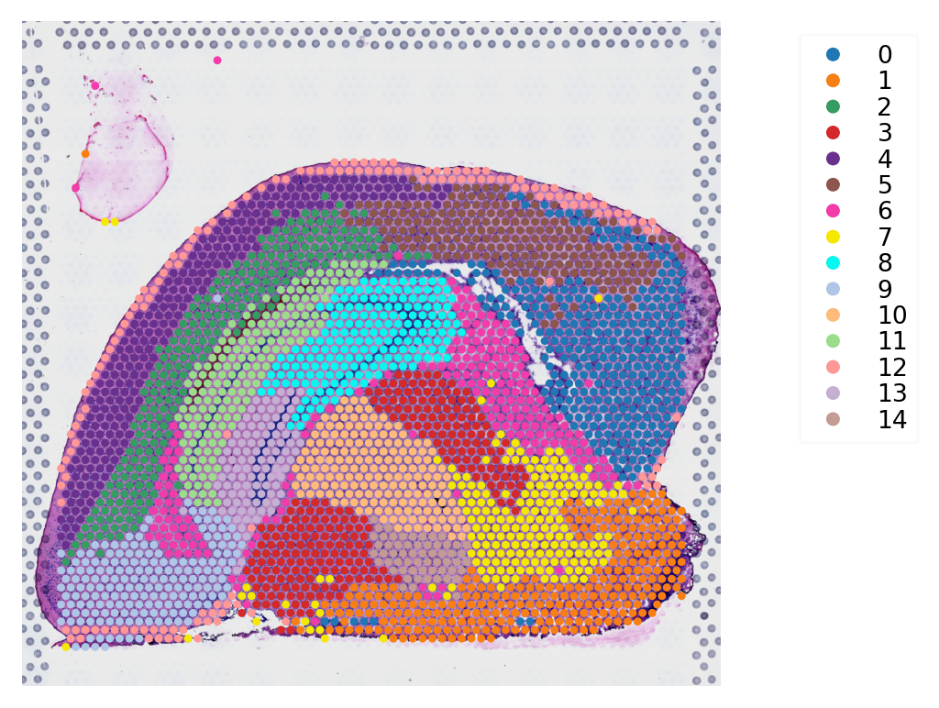

n. IVH_3d_3_110

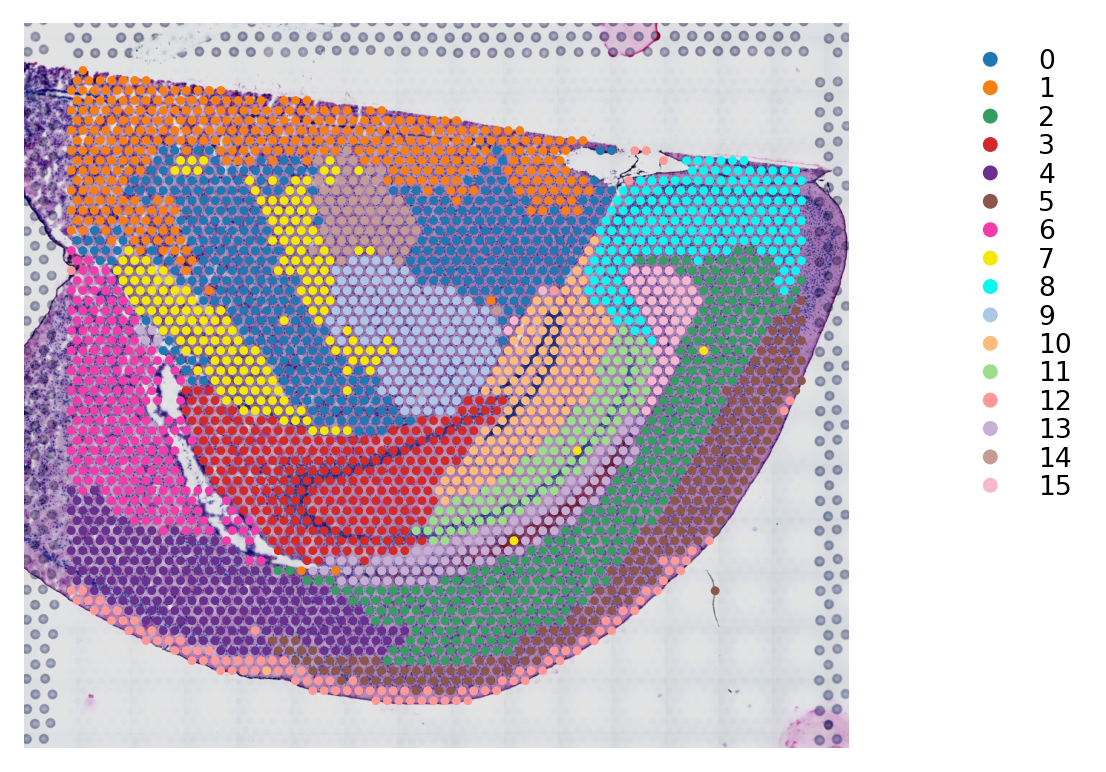

o. IVH_3d_3_130

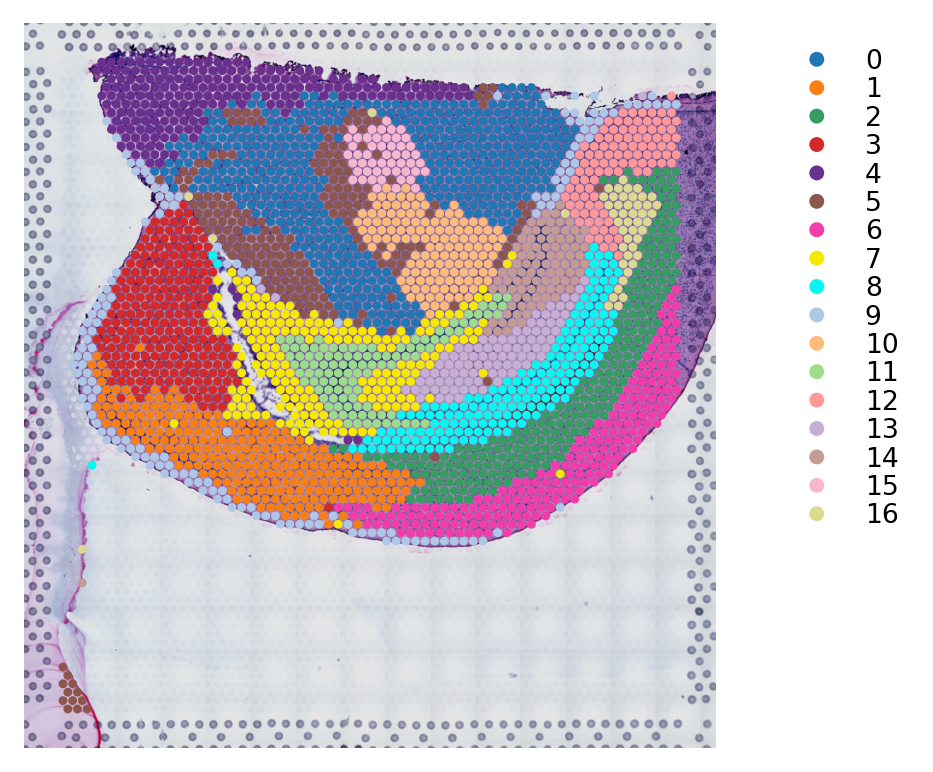

p. IVH_3d_3_150

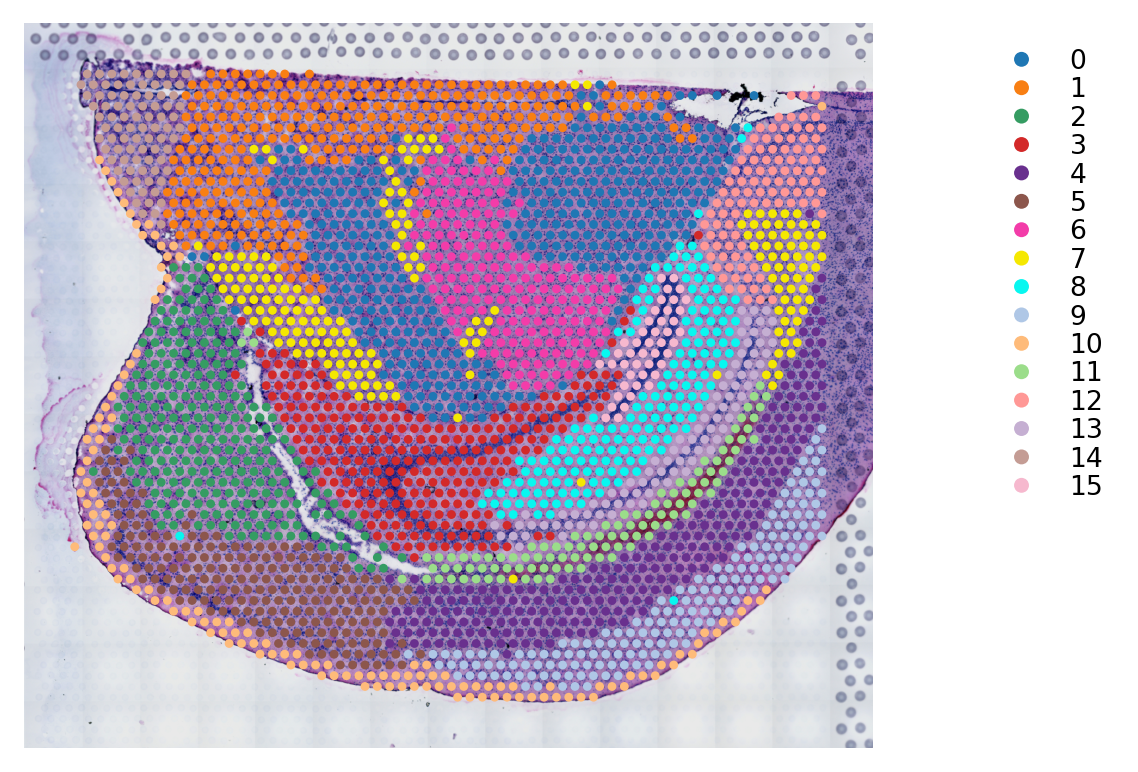

q. IVH_7d_2_50

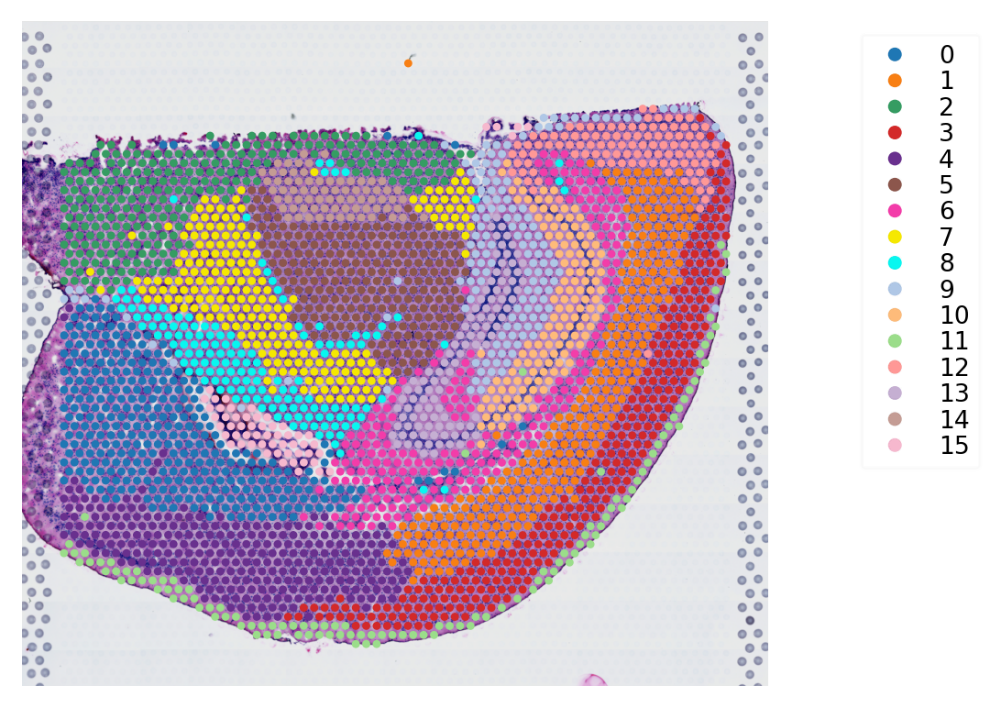

r. IVH_7d_2_80

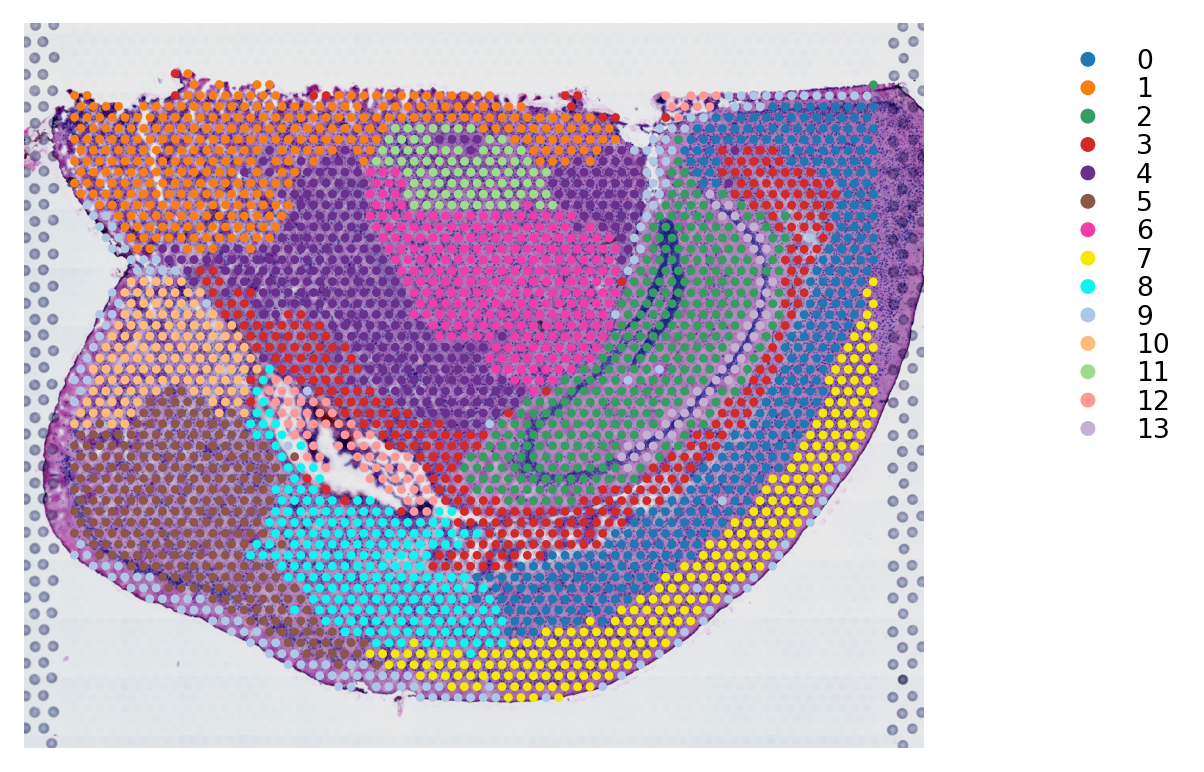

s. IVH_7d_2_110

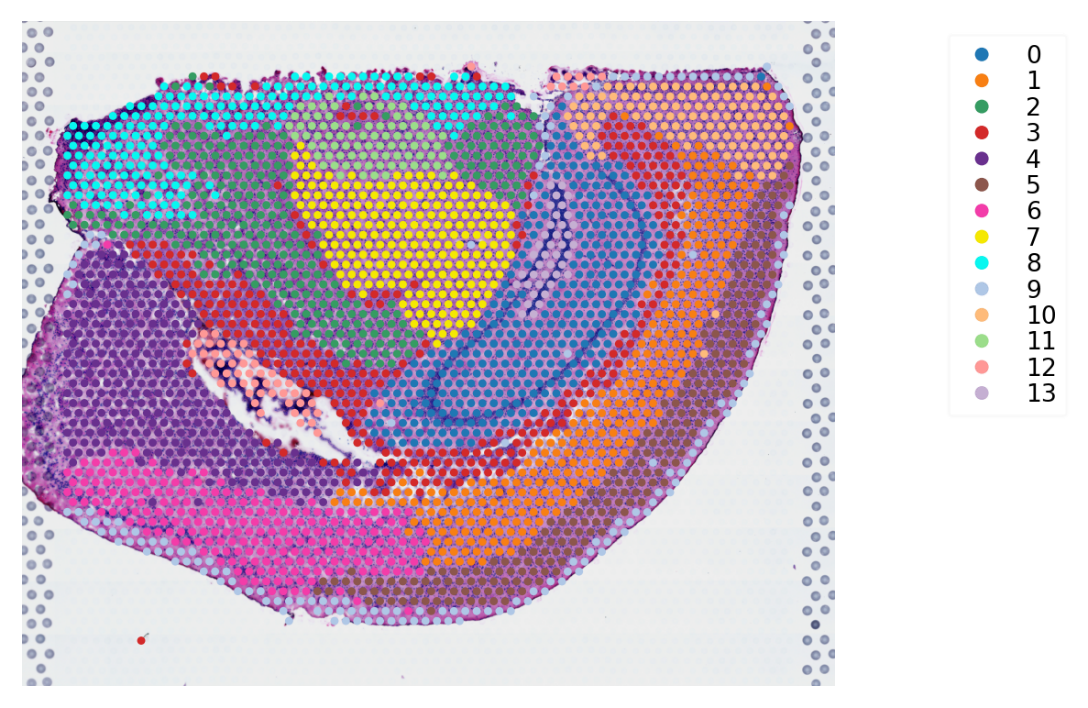

Figure 3 The annotation atlas of cell type clusters for mouse brain

a. IVH_C1_50

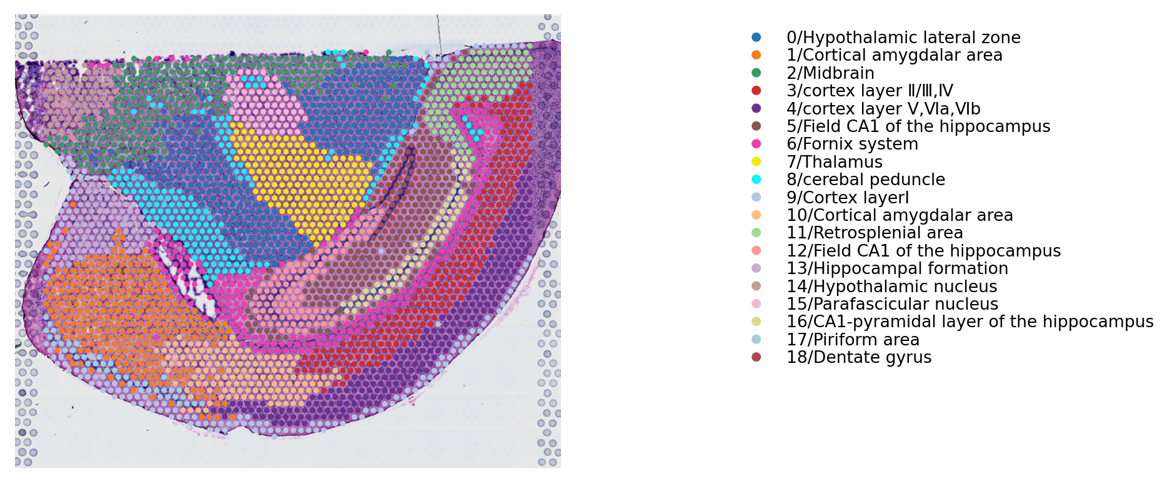

b. IVH_C1_110

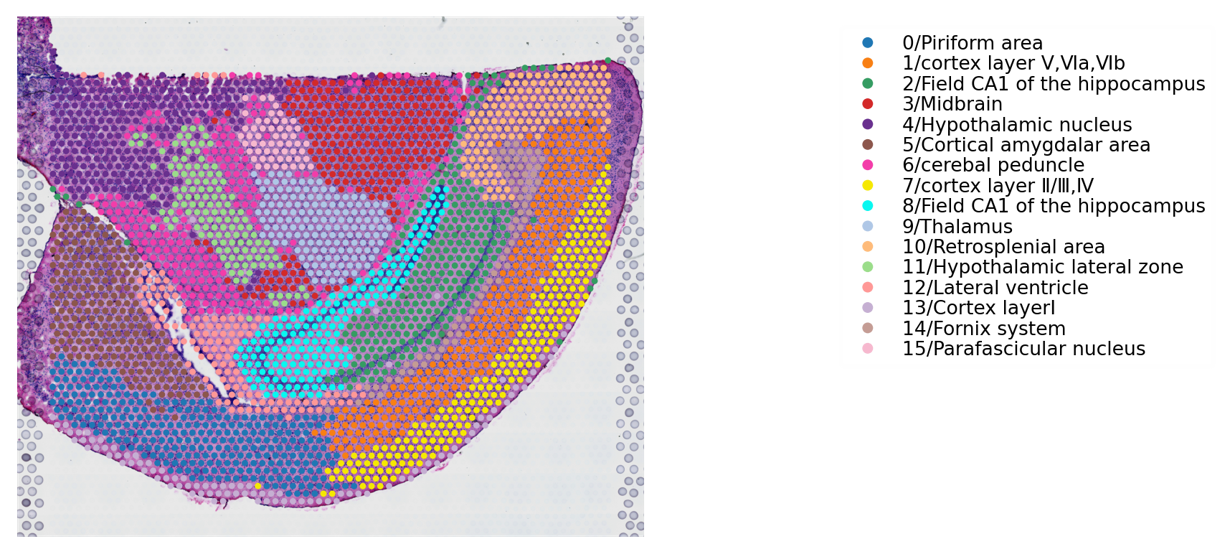

c. IVH_S1_50

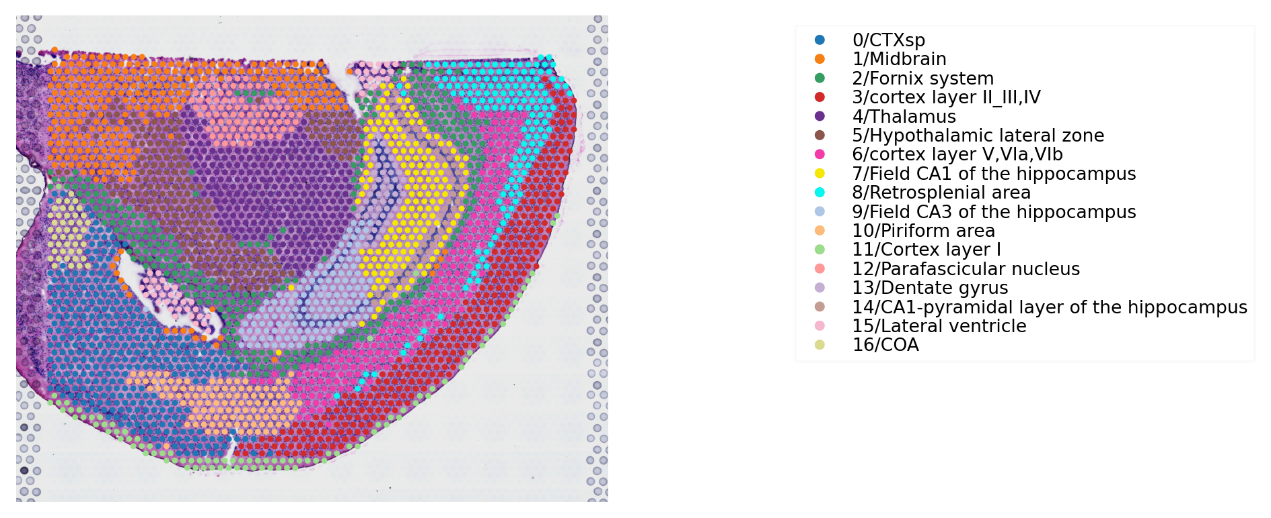

d. IVH_S1_80

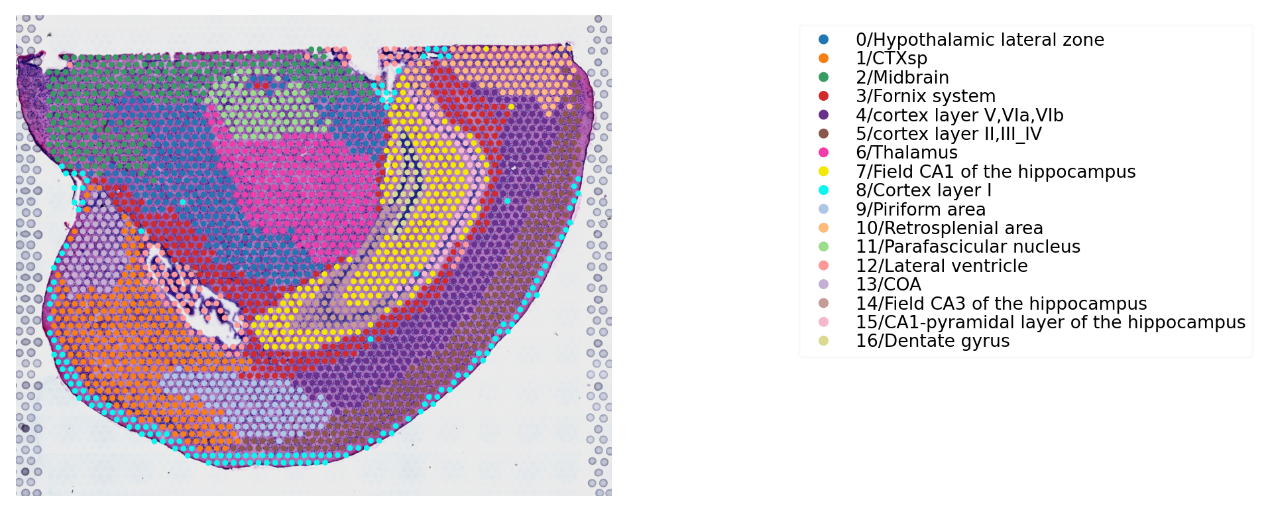

e. IVH_S1_110

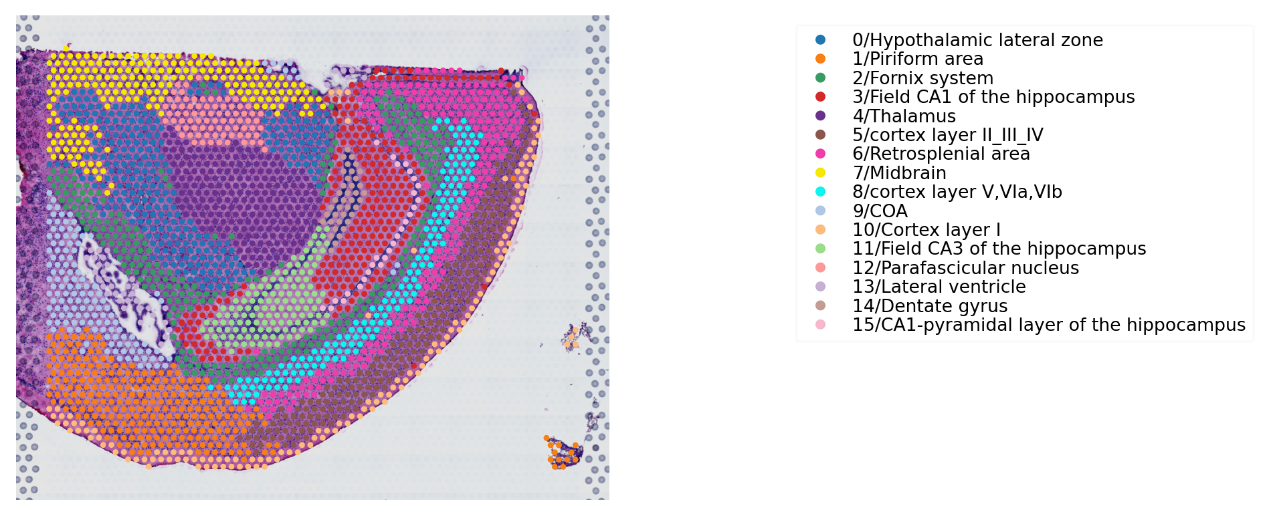

f. IVH_1d_1_20

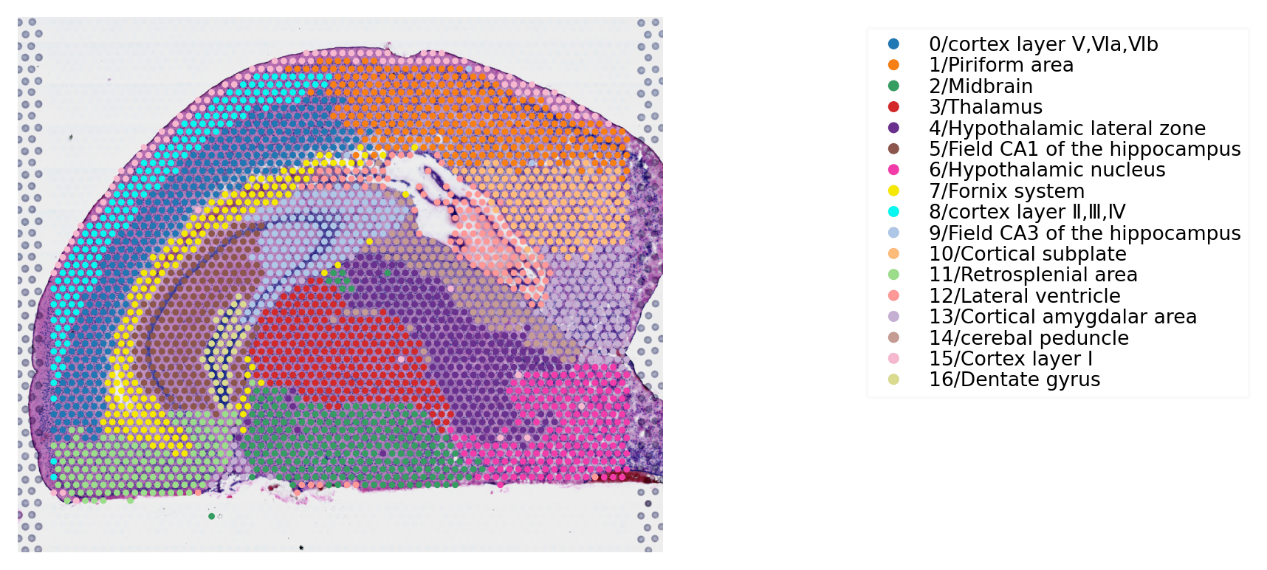

g. IVH_1d_1_50

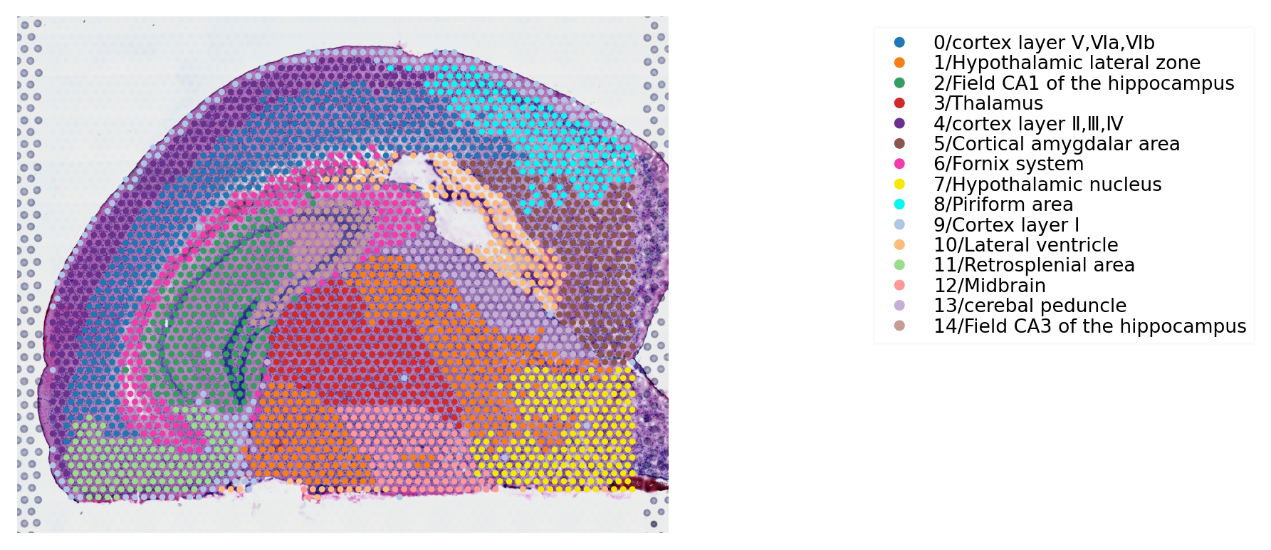

h. IVH_1d_1_80

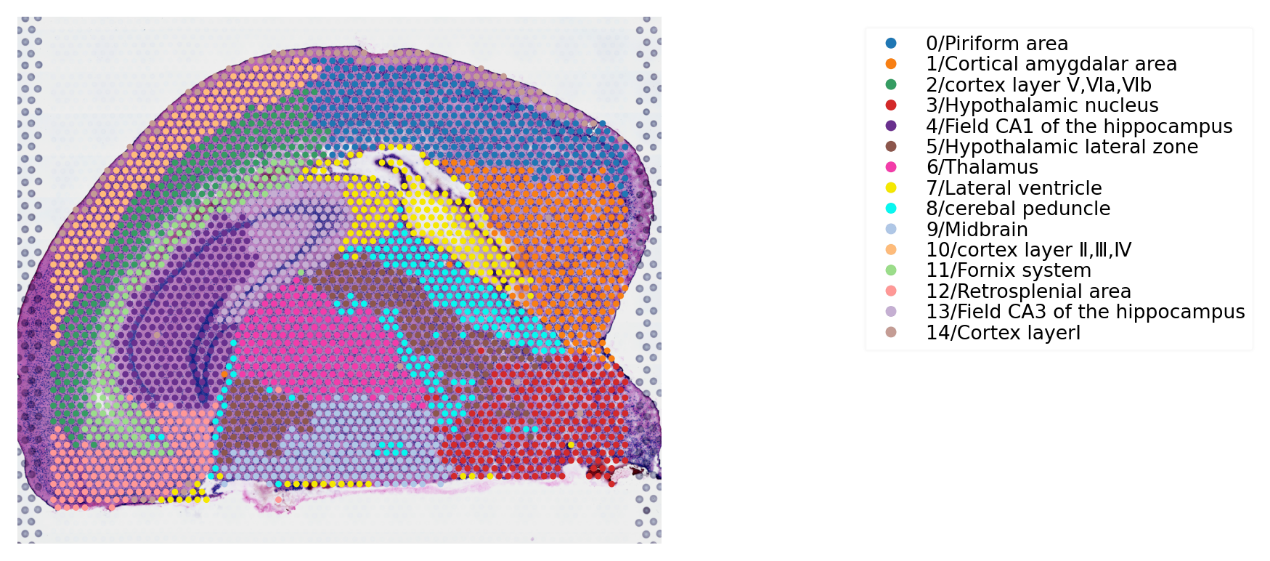

i. IVH_1d_1_110

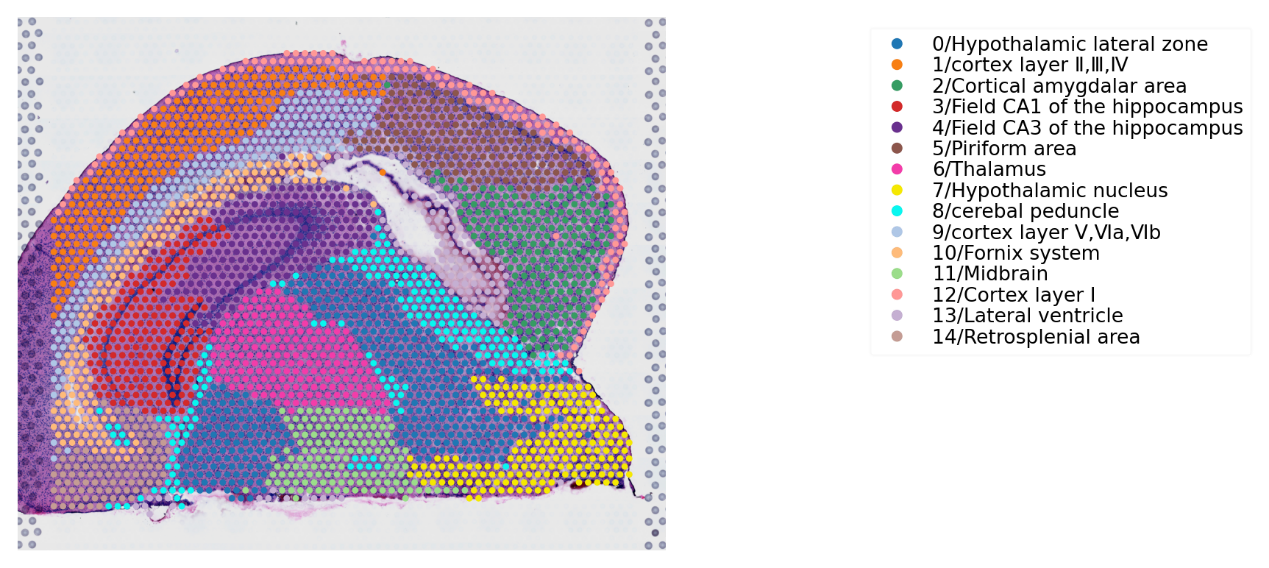

j. IVH_1d_1_130

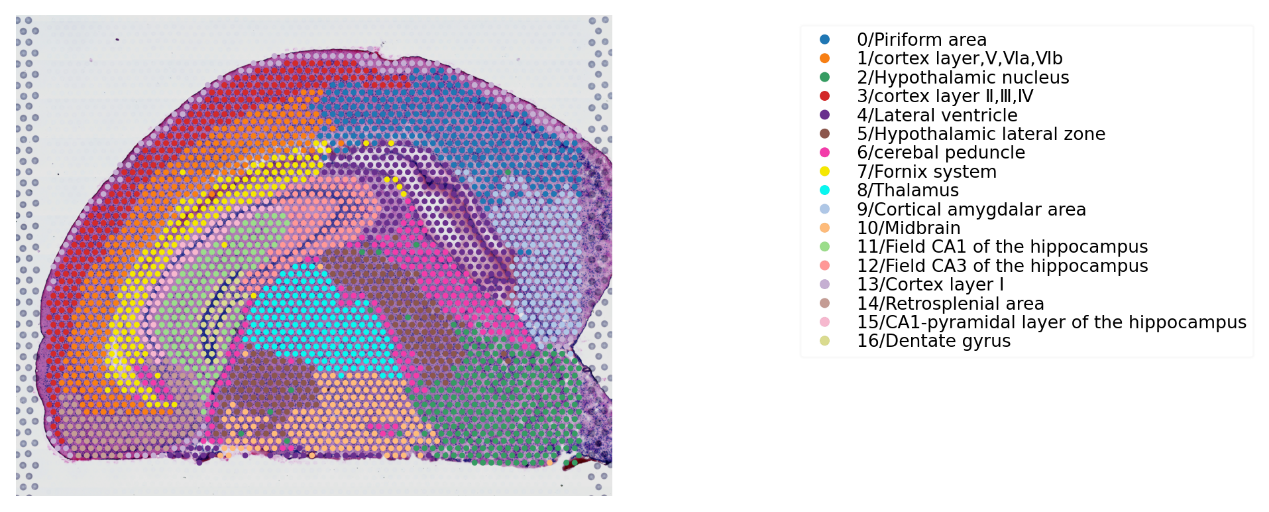

k. IVH_1d_1_150

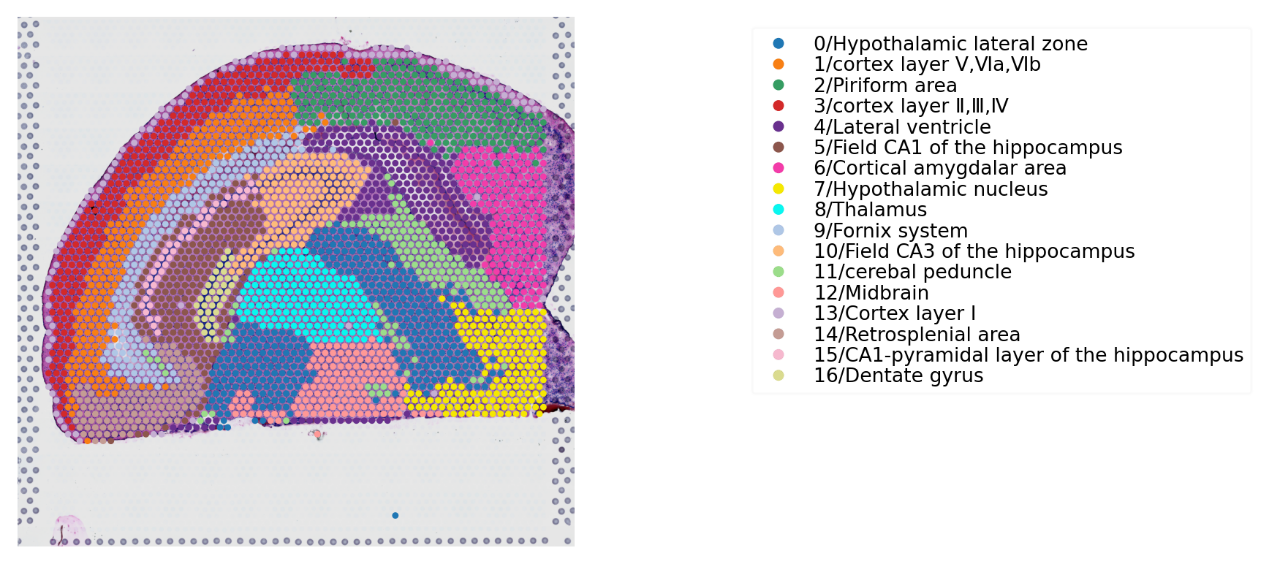

l. IVH_3d_3_20

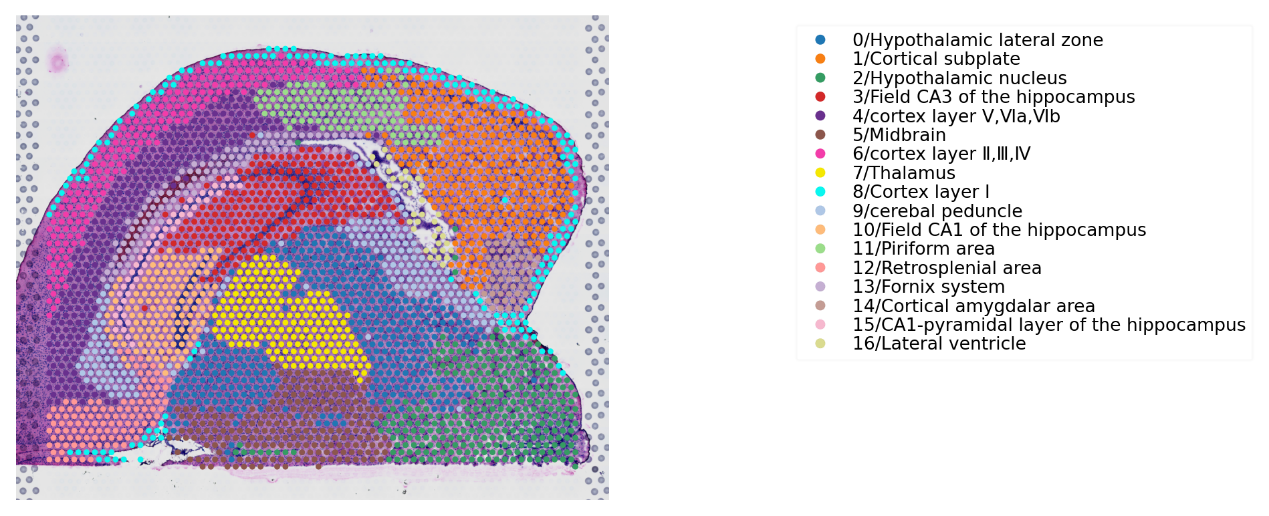

m. IVH_3d_3_50

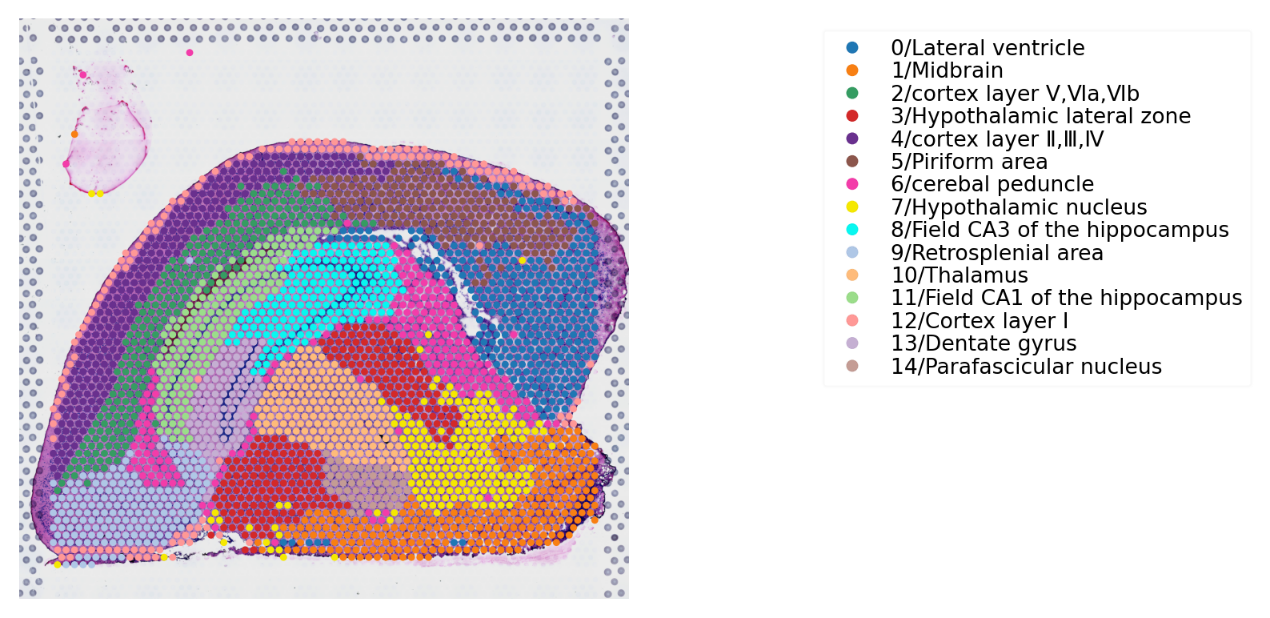

n. IVH_3d_3_110

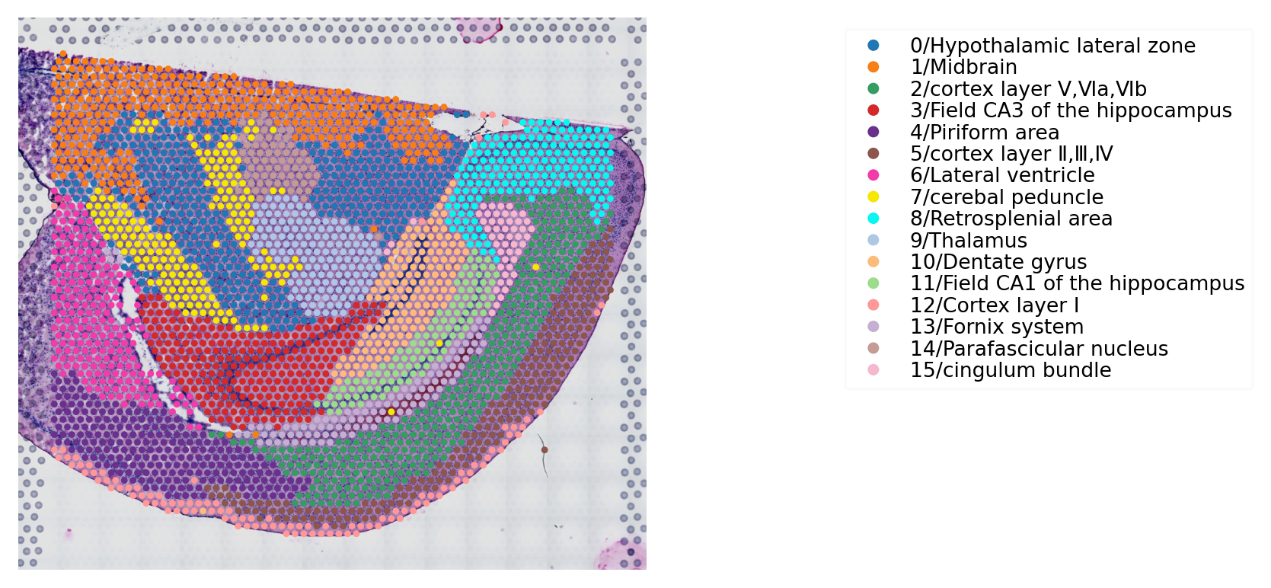

o. IVH_3d_3_130

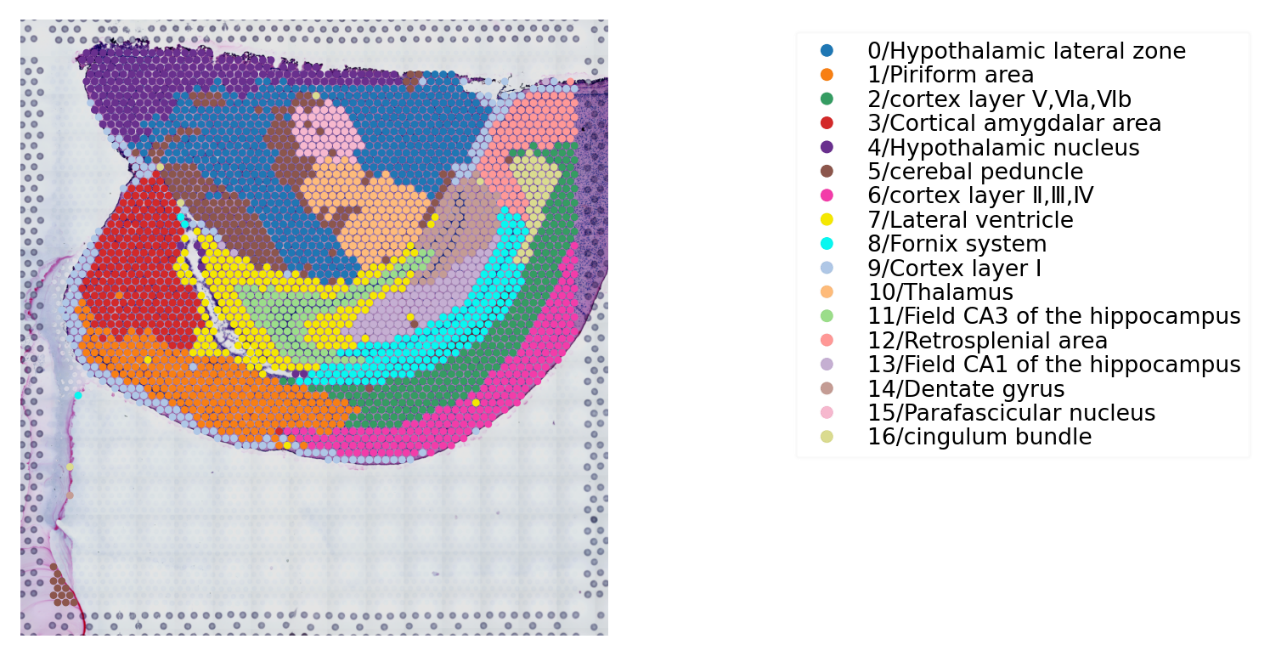

p. IVH_3d_3_150

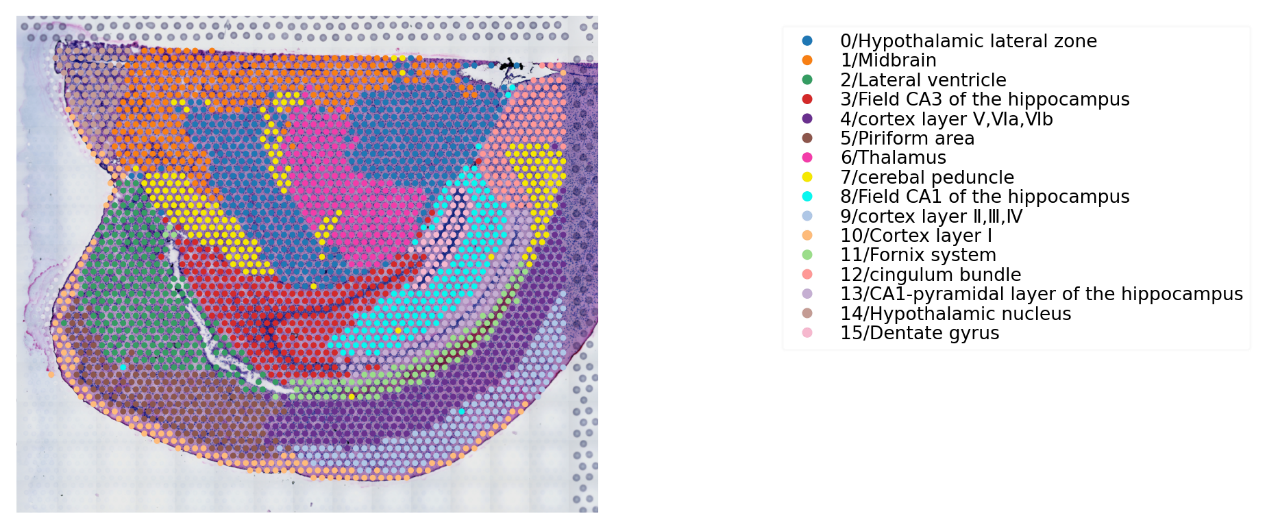

q. IVH_7d_2_50

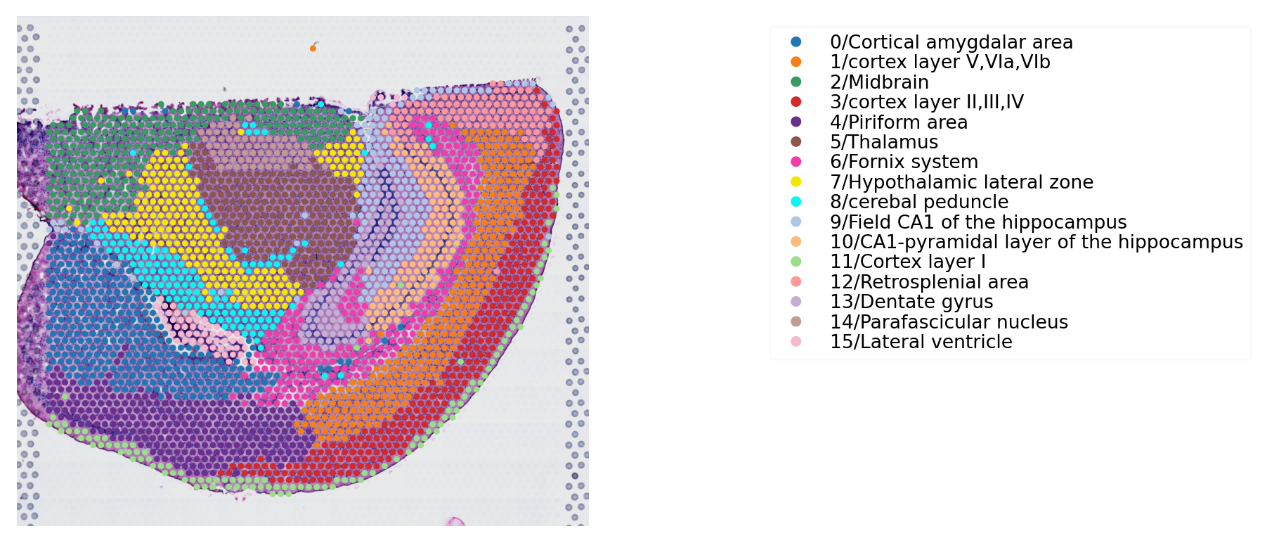

r. IVH_7d_2_80

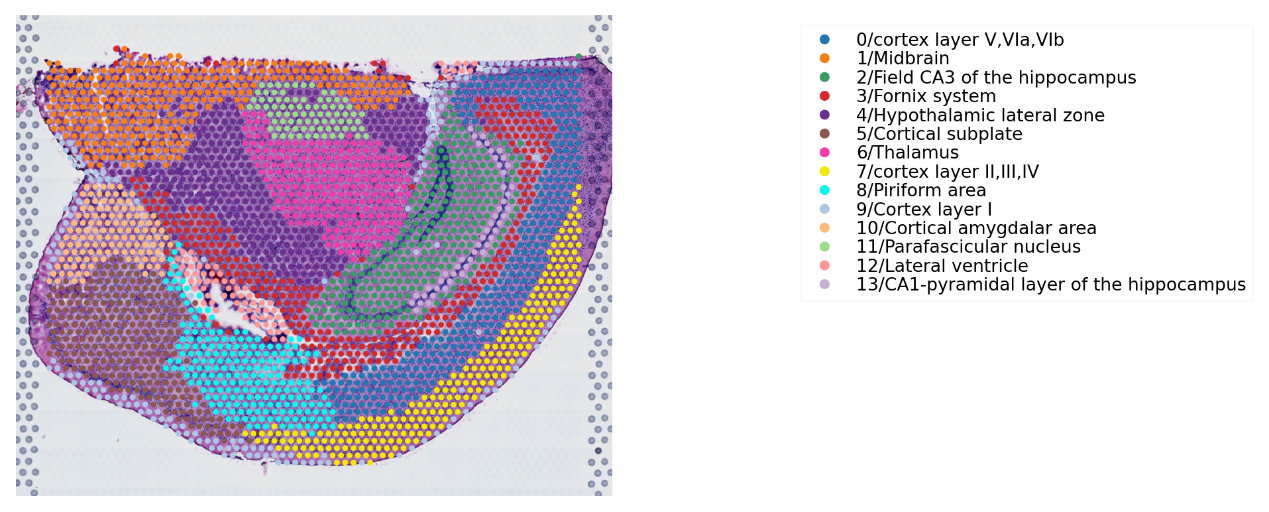

s. IVH_7d_2_110

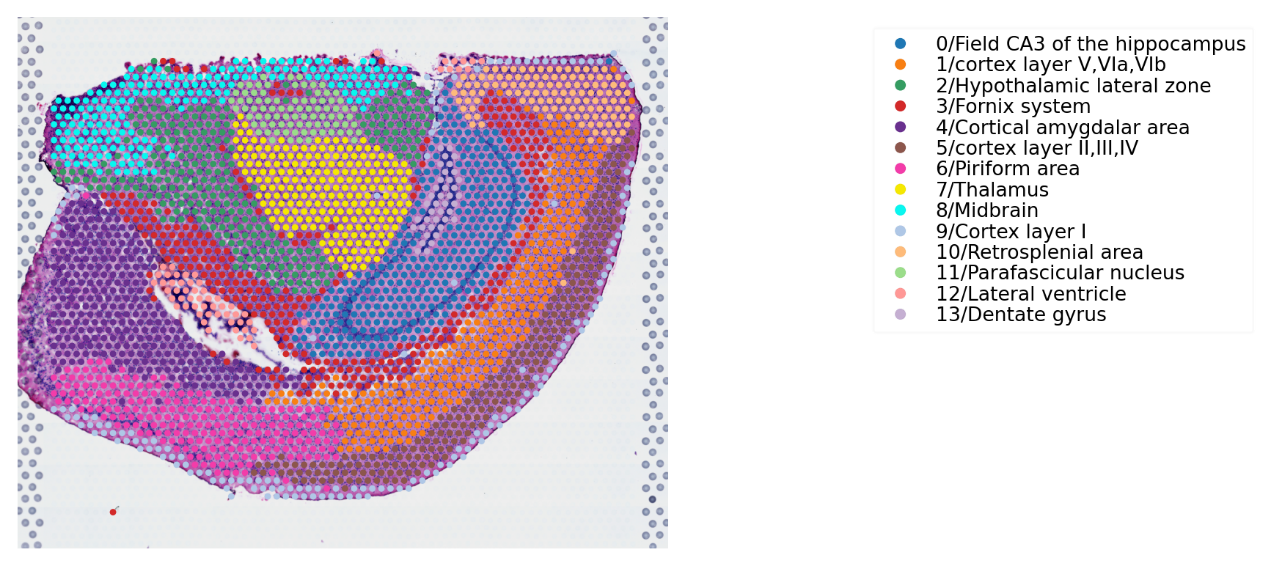

Table 1 The tissue position and spatial gene expression of mouse brain after IVH [*https://github.com/JiayidaerBadai/Spatial-transcriptome.git*](https://github.com/JiayidaerBadai/Spatial-transcriptome.git)

a. IVH_C1_50
b. IVH_C1_110
c. IVH_S1_50
d. IVH_S1_80
e. IVH_S1_110
f. IVH_1d_1_20
g. IVH_1d_1_50
h. IVH_1d_1_80
i. IVH_1d_1_110
j. IVH_1d_1_130
k. IVH_1d_1_150
l. IVH_3d_3_20
m. IVH_3d_3_50
n. IVH_3d_3_110
o. IVH_3d_3_130
p. IVH_3d_3_150
q. IVH_7d_2_50
r. IVH_7d_2_80
s. IVH_7d_2_110
2. **Supplementary of Result 2 (SR2)**

Figure 1 The planar diffusion pseudotime plots

a. IVH_S1_50

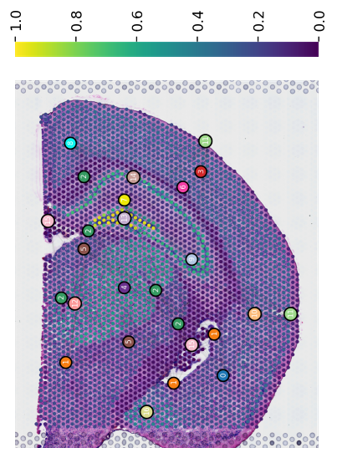

b. IVH_S1_80

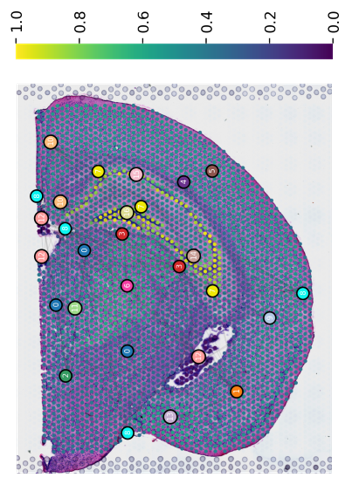

c. IVH_S1_110

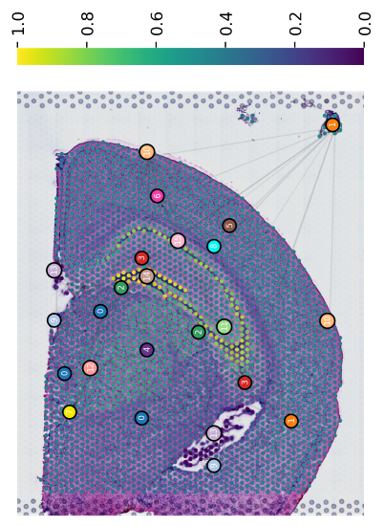

d. IVH_1d_1_20

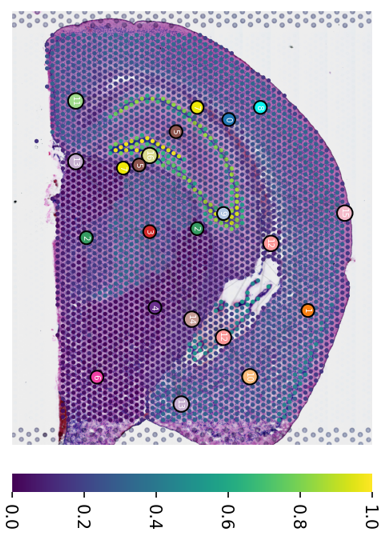

e. IVH_1d_1_50

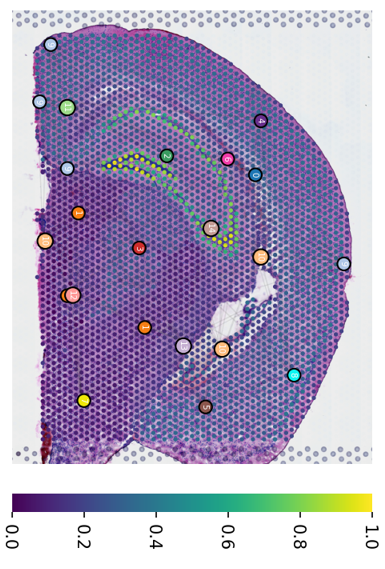

f. IVH_1d_1_80

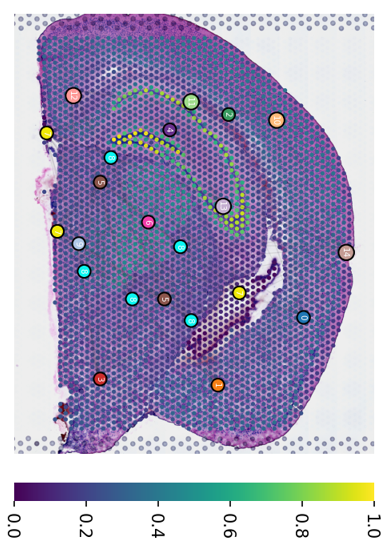

g. IVH_1d_1_110

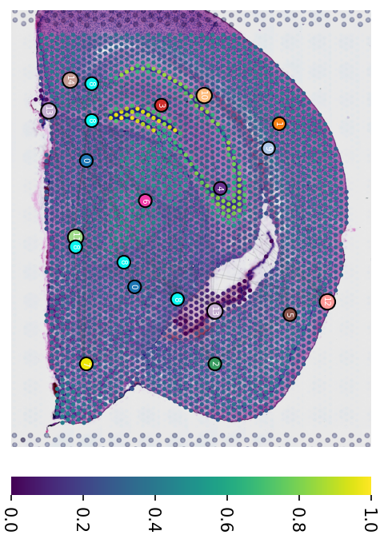

h. IVH_1d_1_130

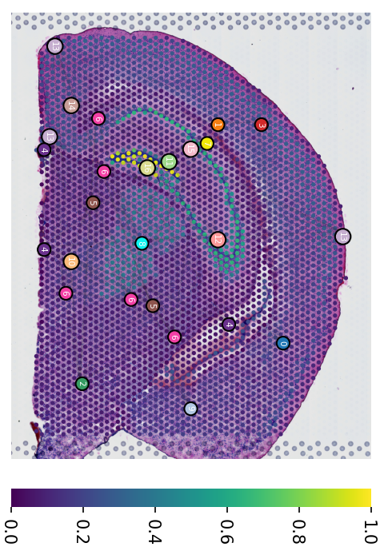

i. IVH_1d_1_150

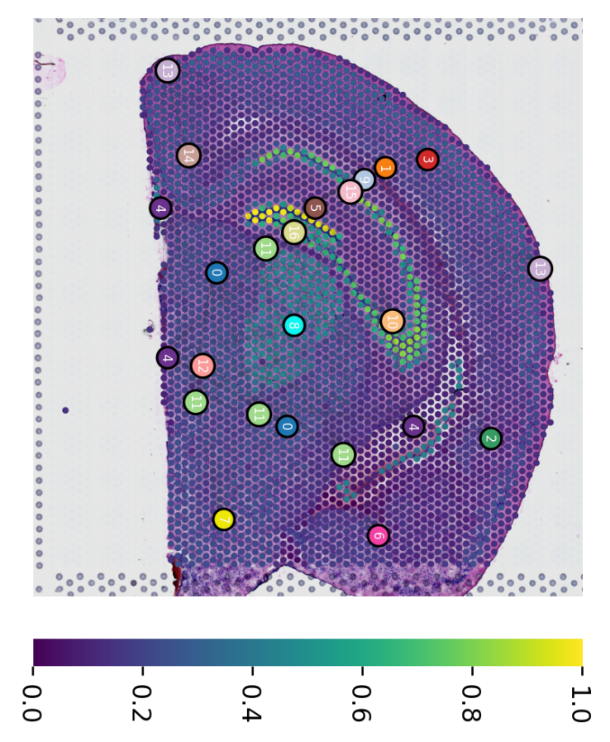

j. IVH_3d_3_20

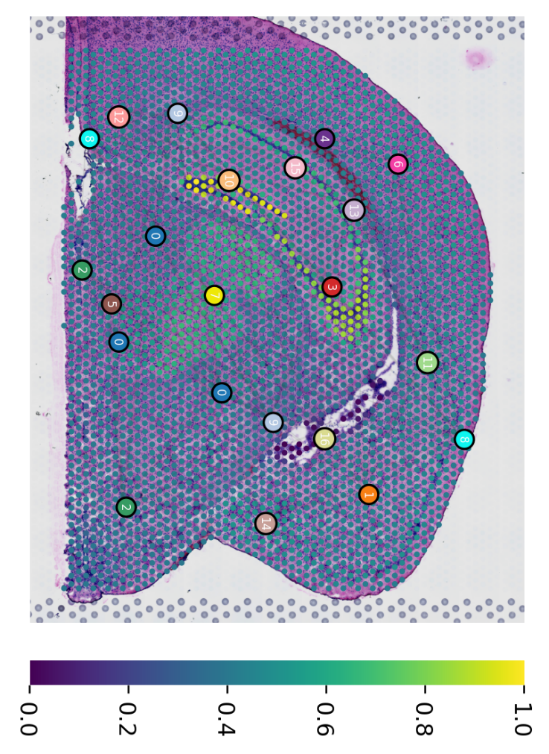

k. IVH_3d_3_50

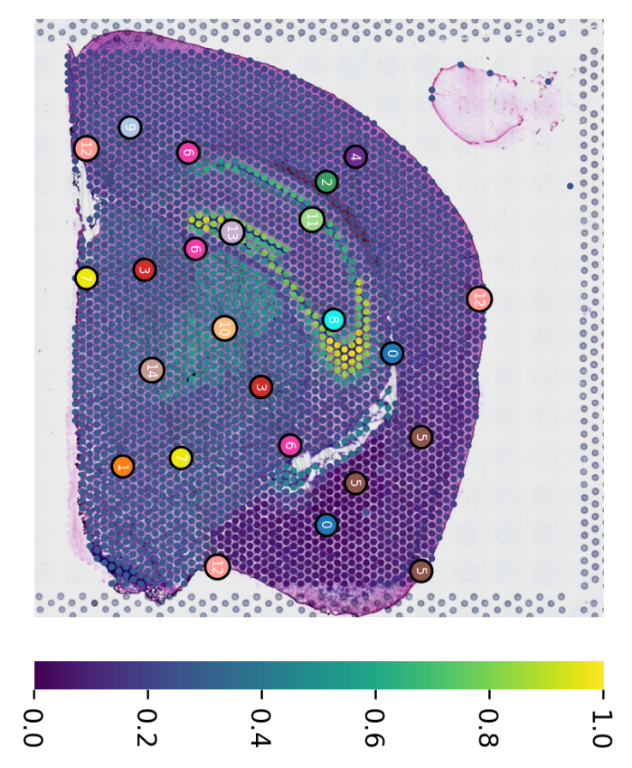

l. IVH_3d_3_110

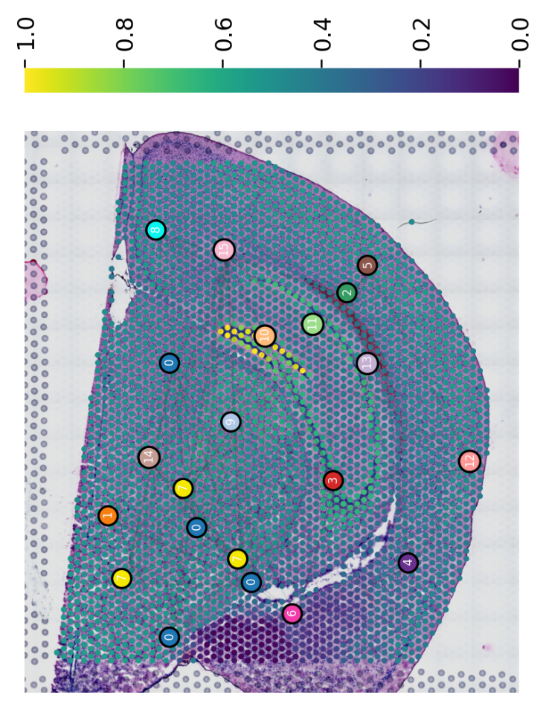

m. IVH_3d_3_130

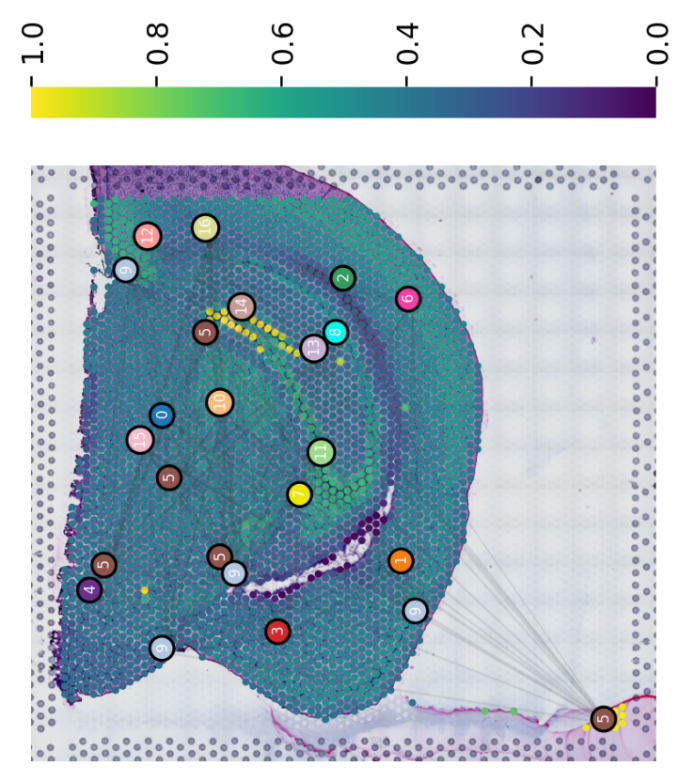

n. IVH_3d_3_150

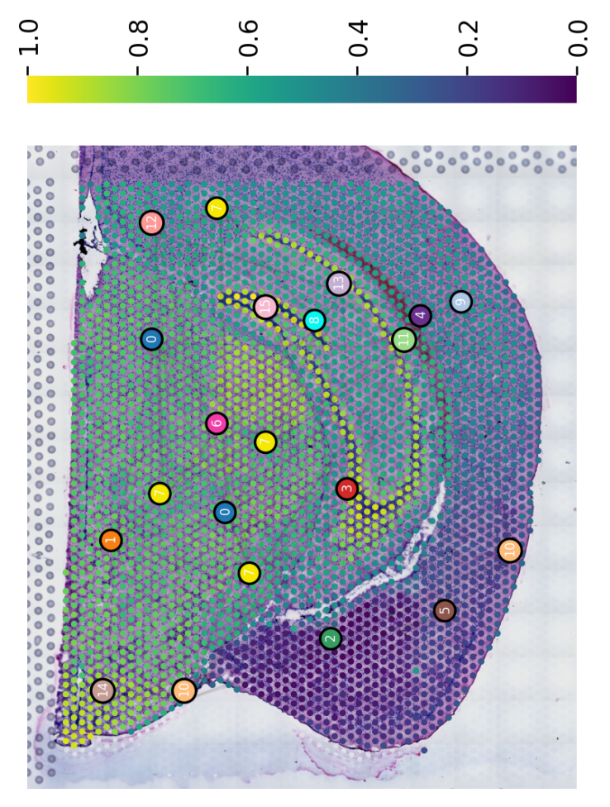

o. IVH_7d_2_50

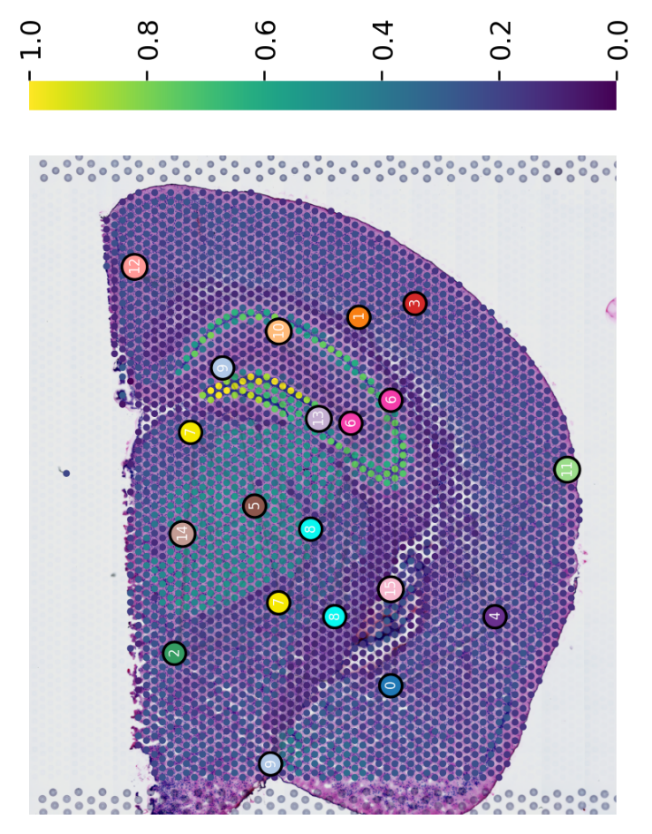

p. IVH_7d_2_80

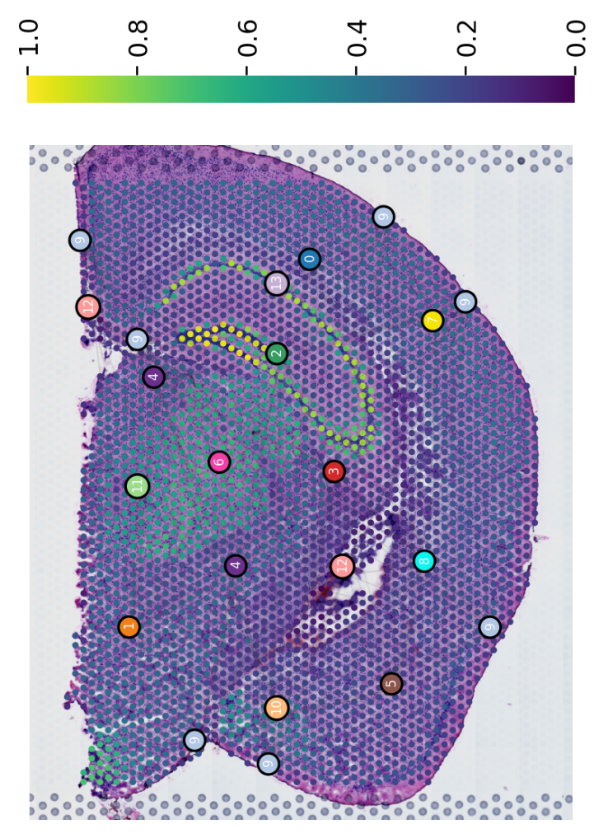

q. IVH_7d_2_110

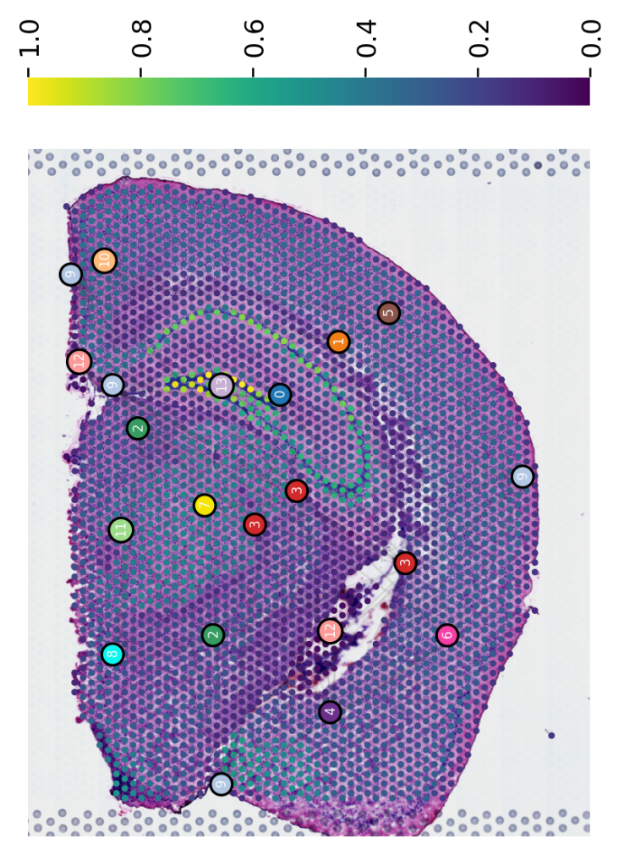

Figure 2 The planar global Pseudo-space-time trajectories

a. IVH_S1_50

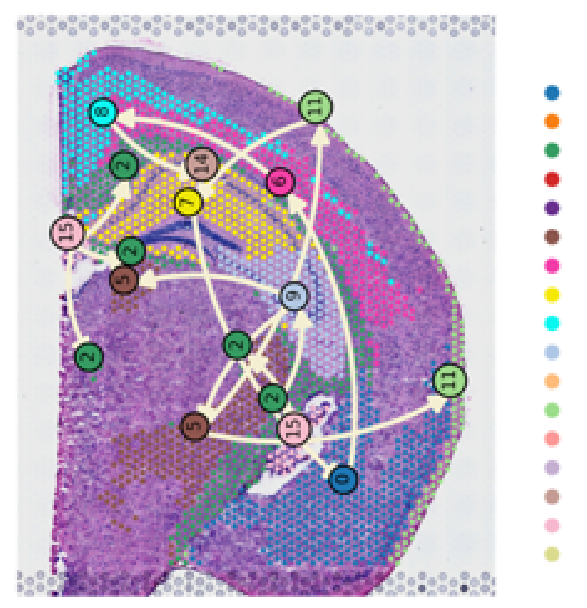

b. IVH_S1_80

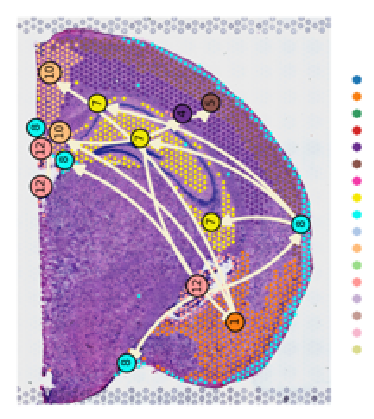

c. IVH_S1_110

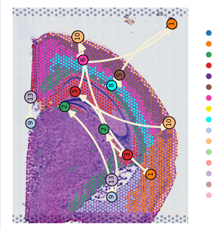

d. IVH_1d_1_20

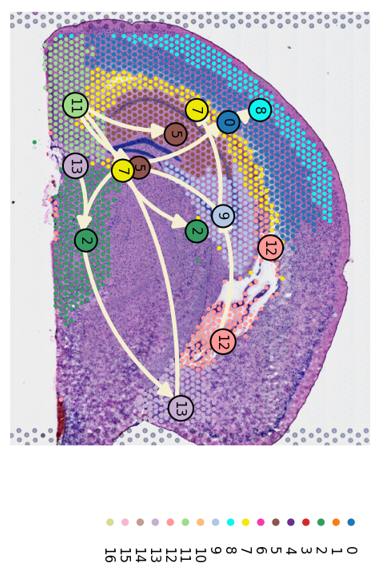

e. IVH_1d_1_50

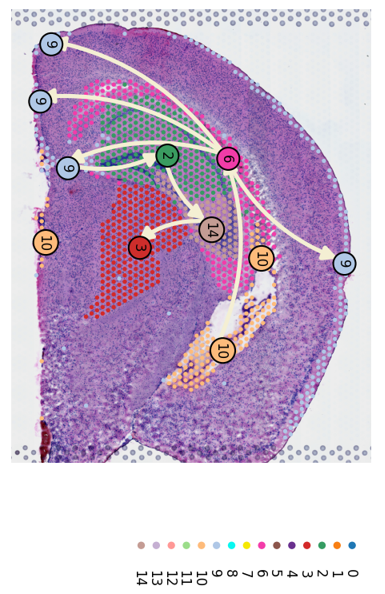

f. IVH_1d_1_80

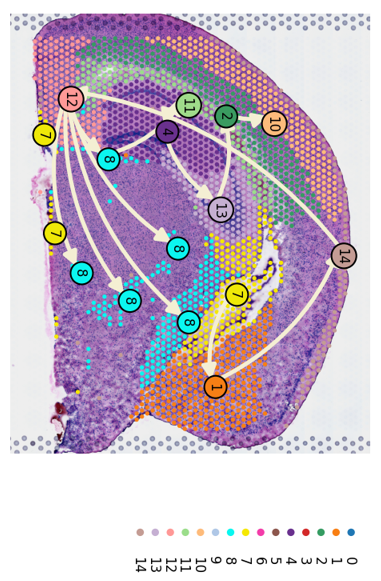

g. IVH_1d_1_110

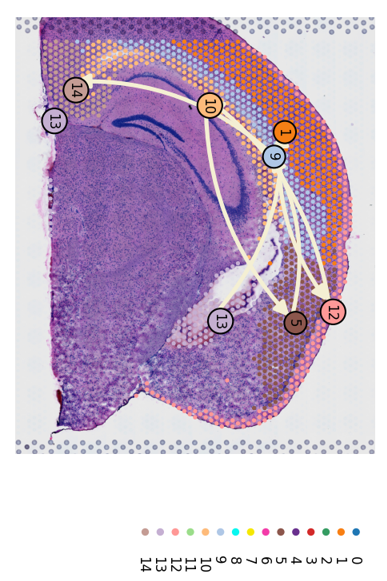

h. IVH_1d_1_130

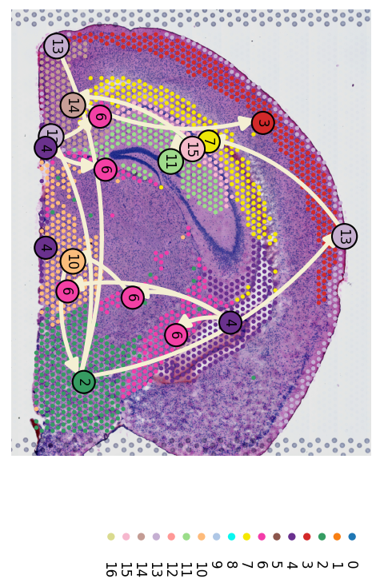

i. IVH_1d_1_150

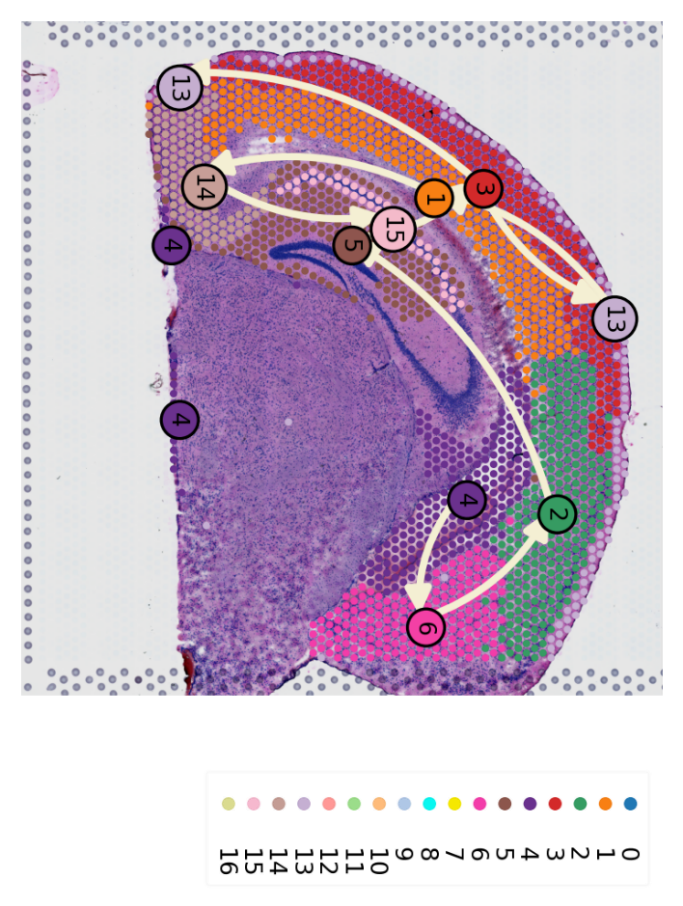

j. IVH_3d_3_20

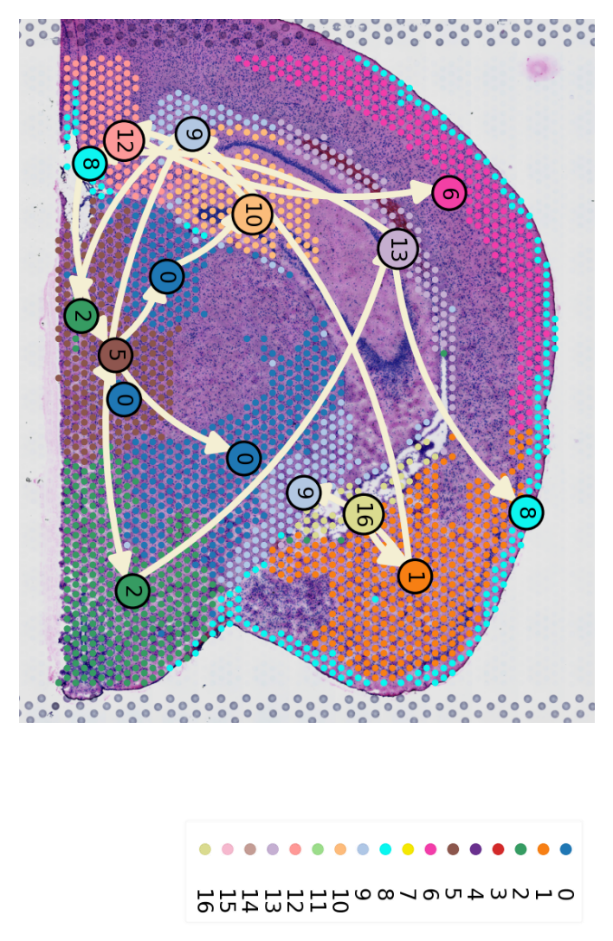

k. IVH_3d_3_50

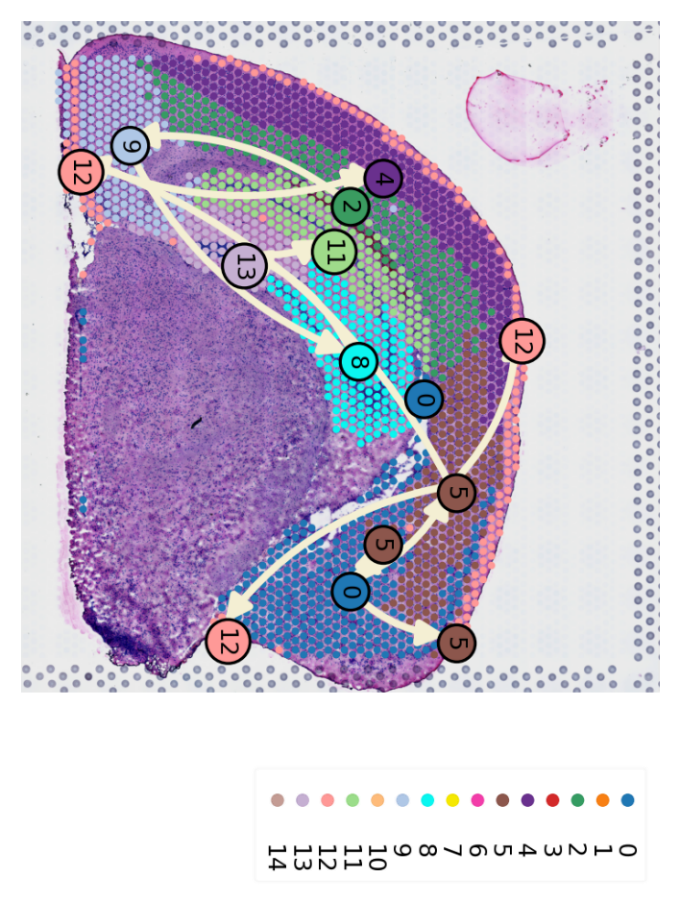

l. IVH_3d_3_110

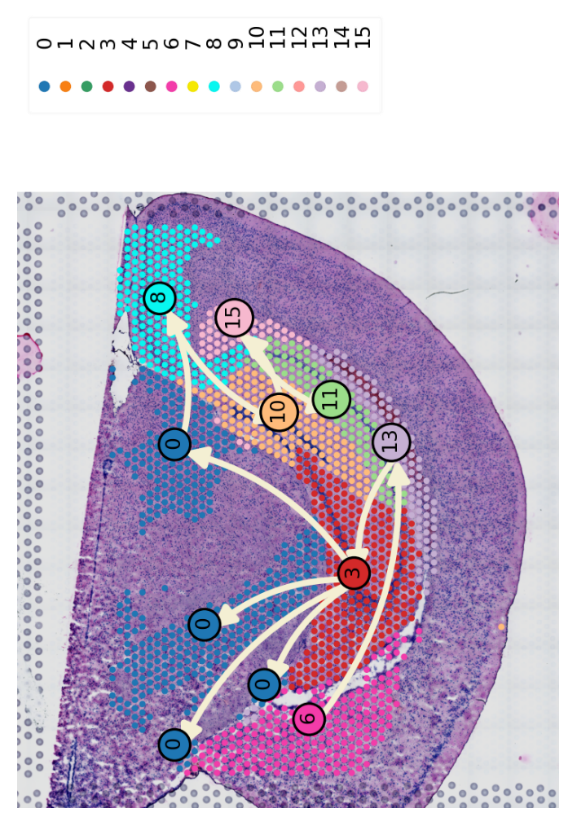

m. IVH_3d_3_130

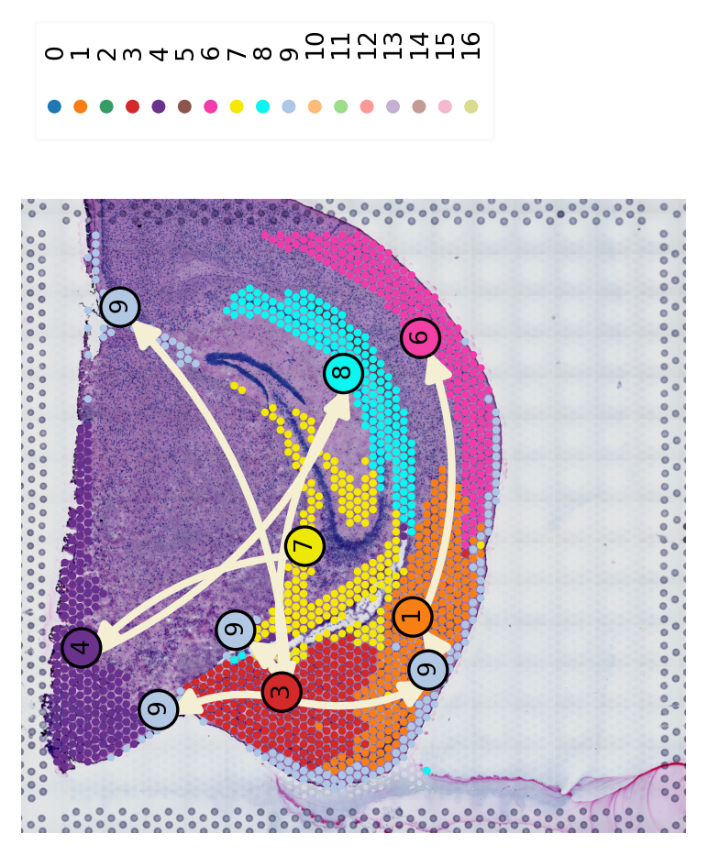

n. IVH_3d_3_150

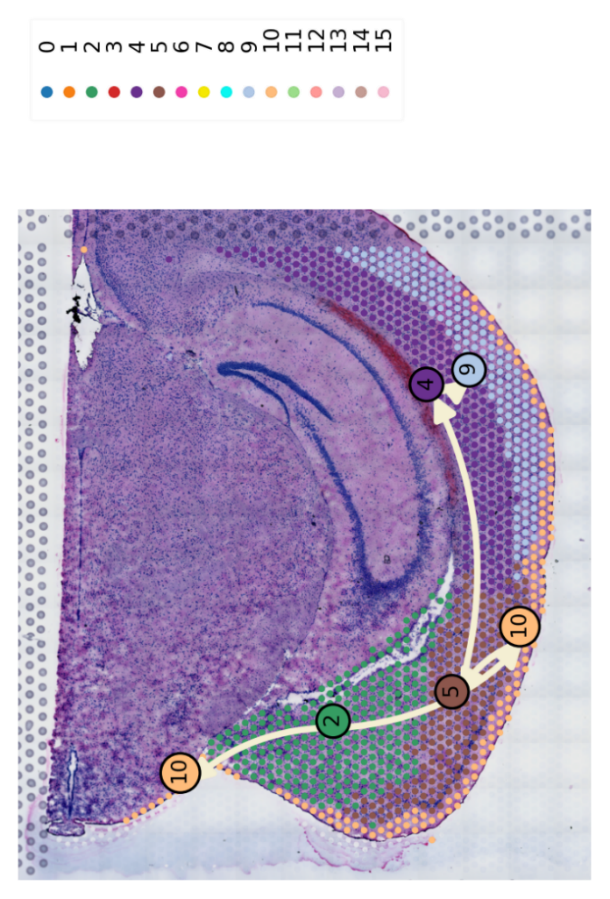

o. IVH_7d_2_50

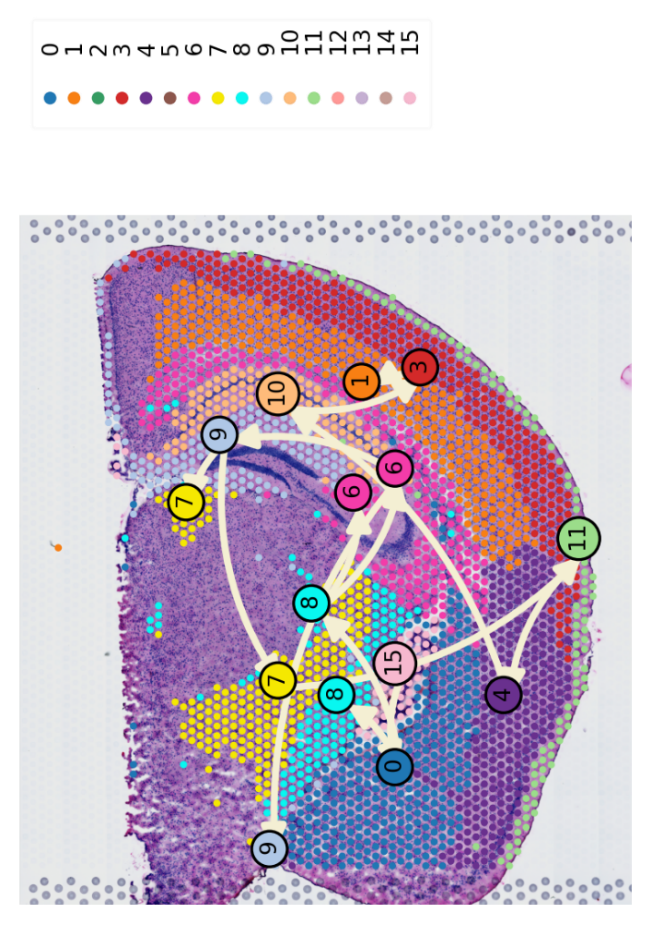

p. IVH_7d_2_80

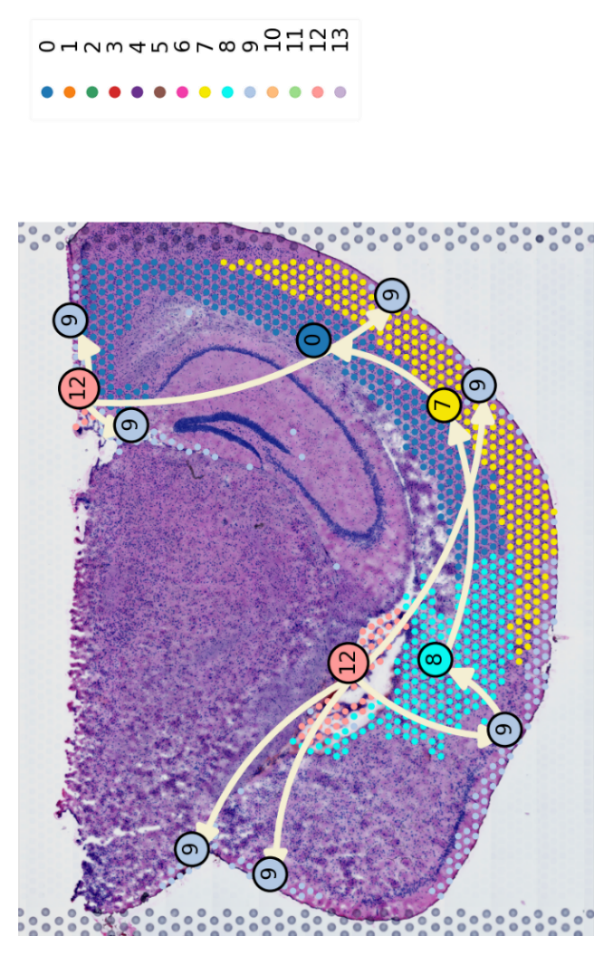

q. IVH_7d_2_110

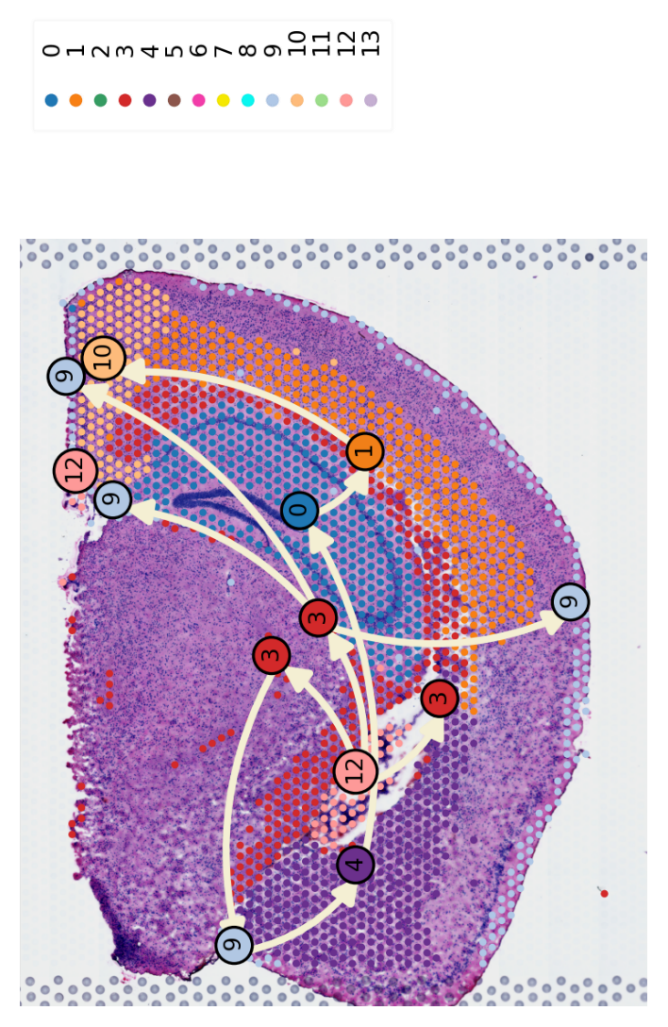

Figure 3 3D global Pseudo-space-time trajectories

a. IVH_S1

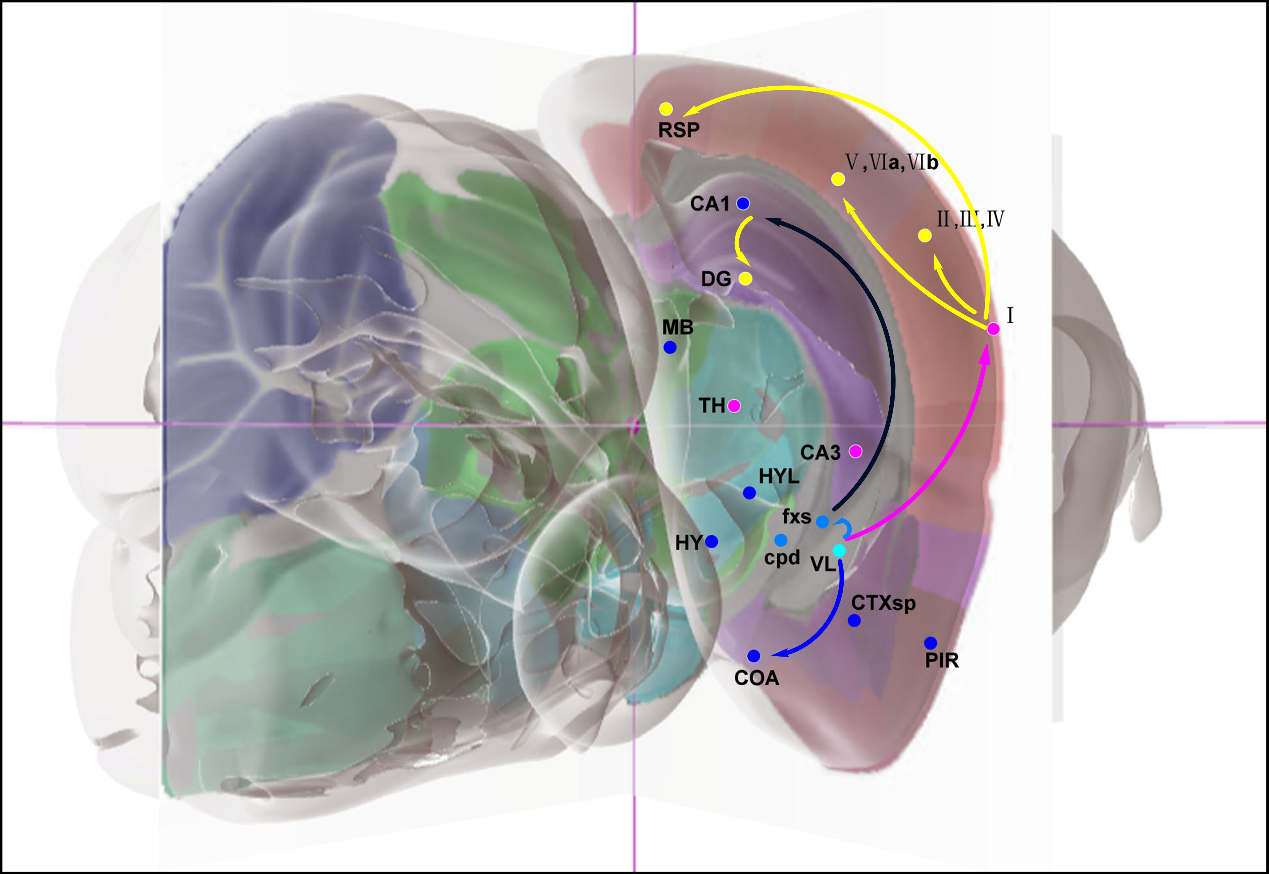

b. Trajectory bundle I of IVH_1d

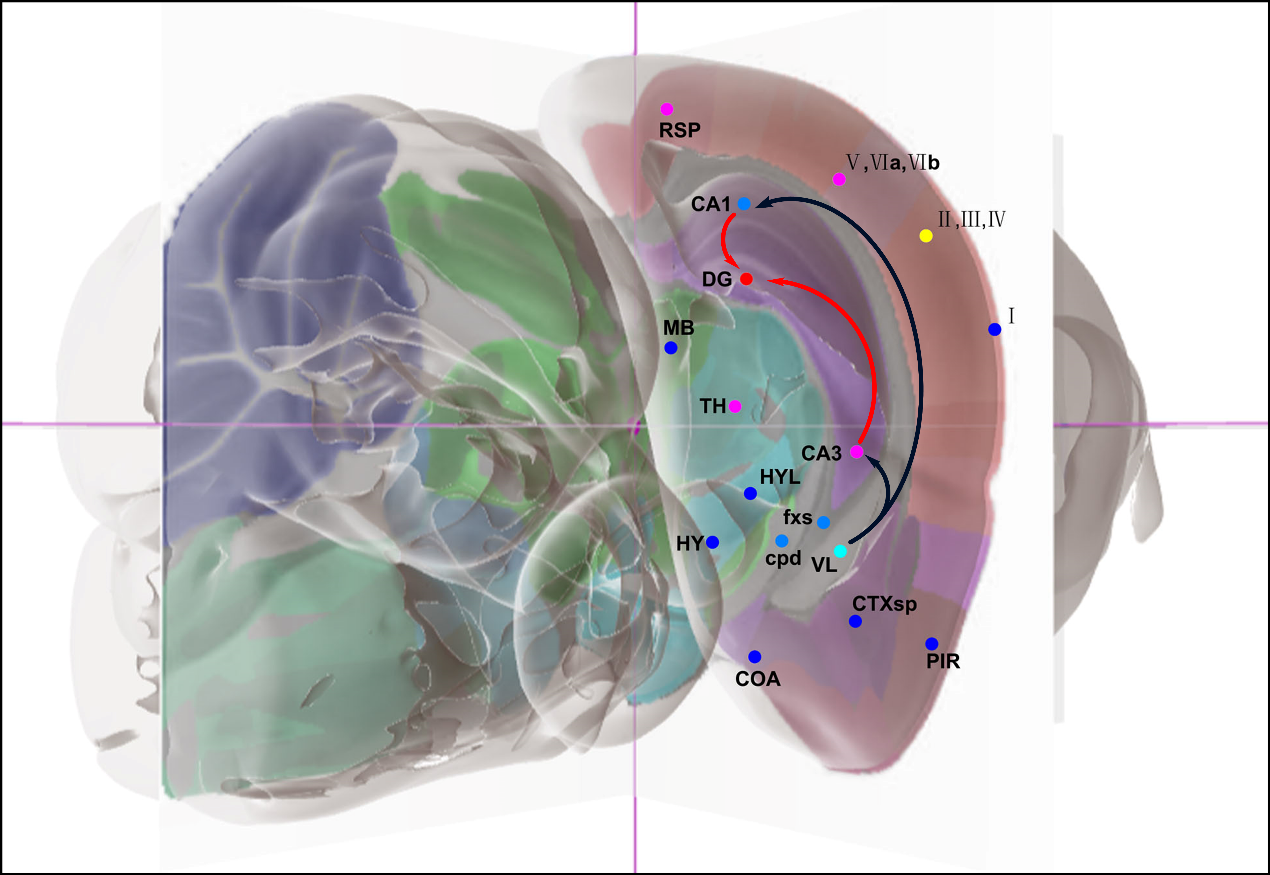

c. Trajectory bundle II of IVH_1d

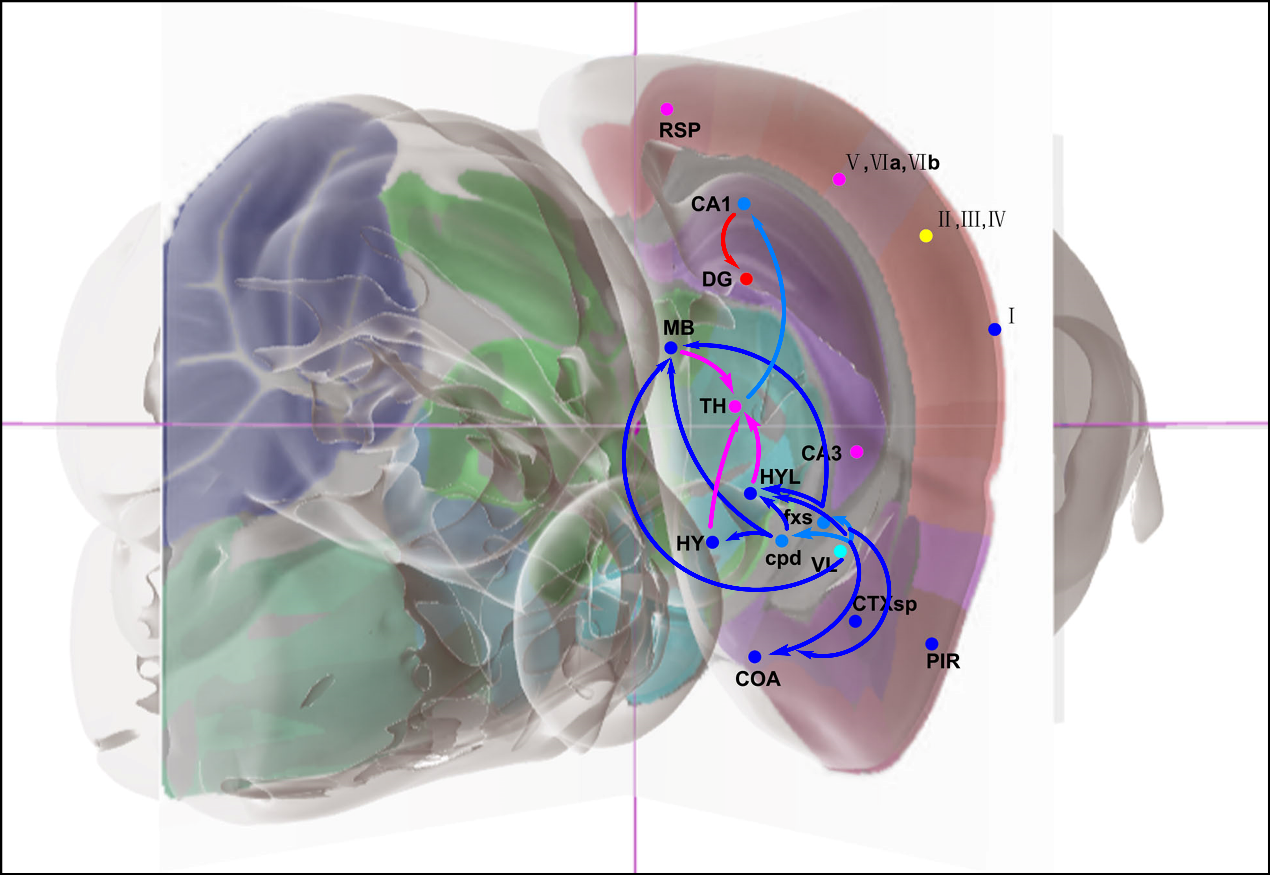

d. Trajectory bundle III of IVH_1d

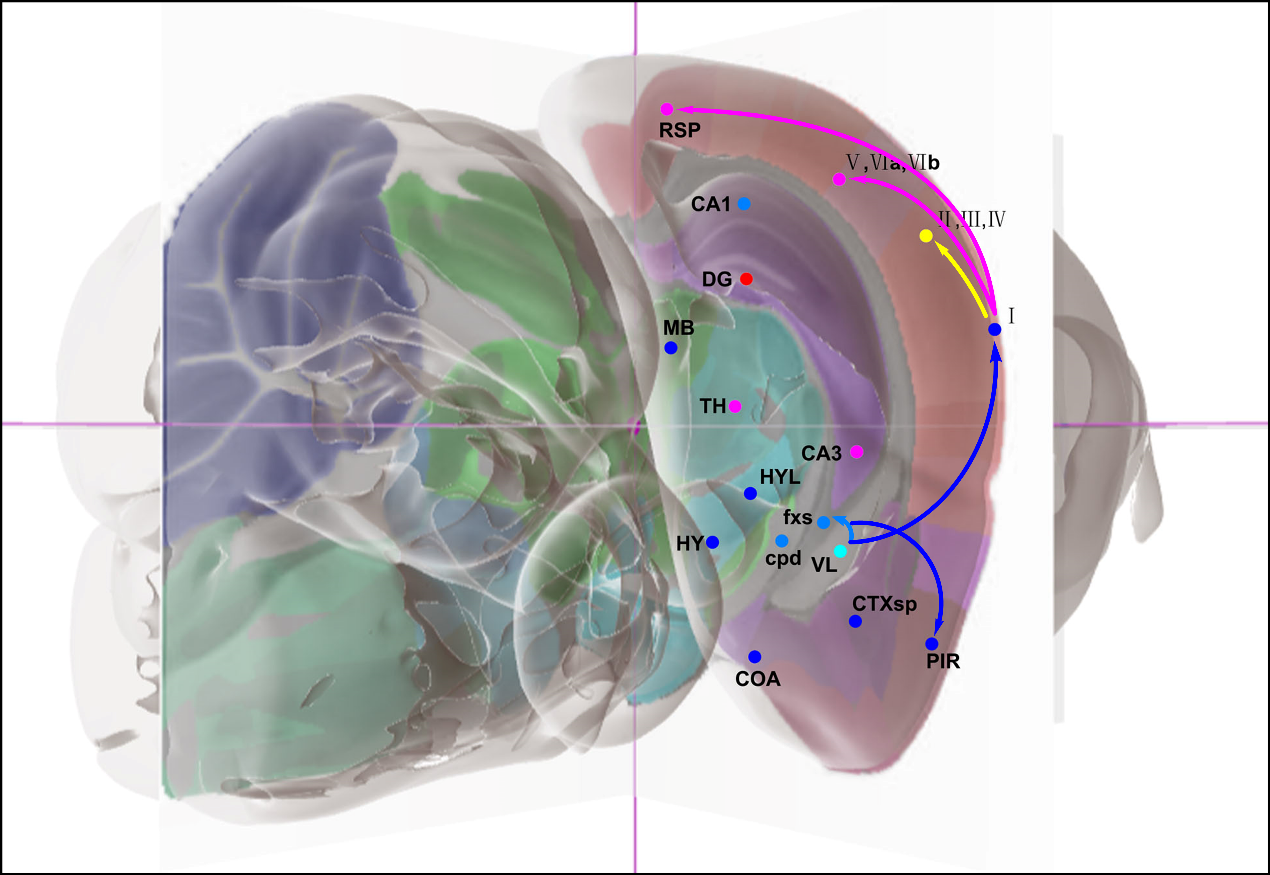

e. Trajectory bundle I of IVH_3d

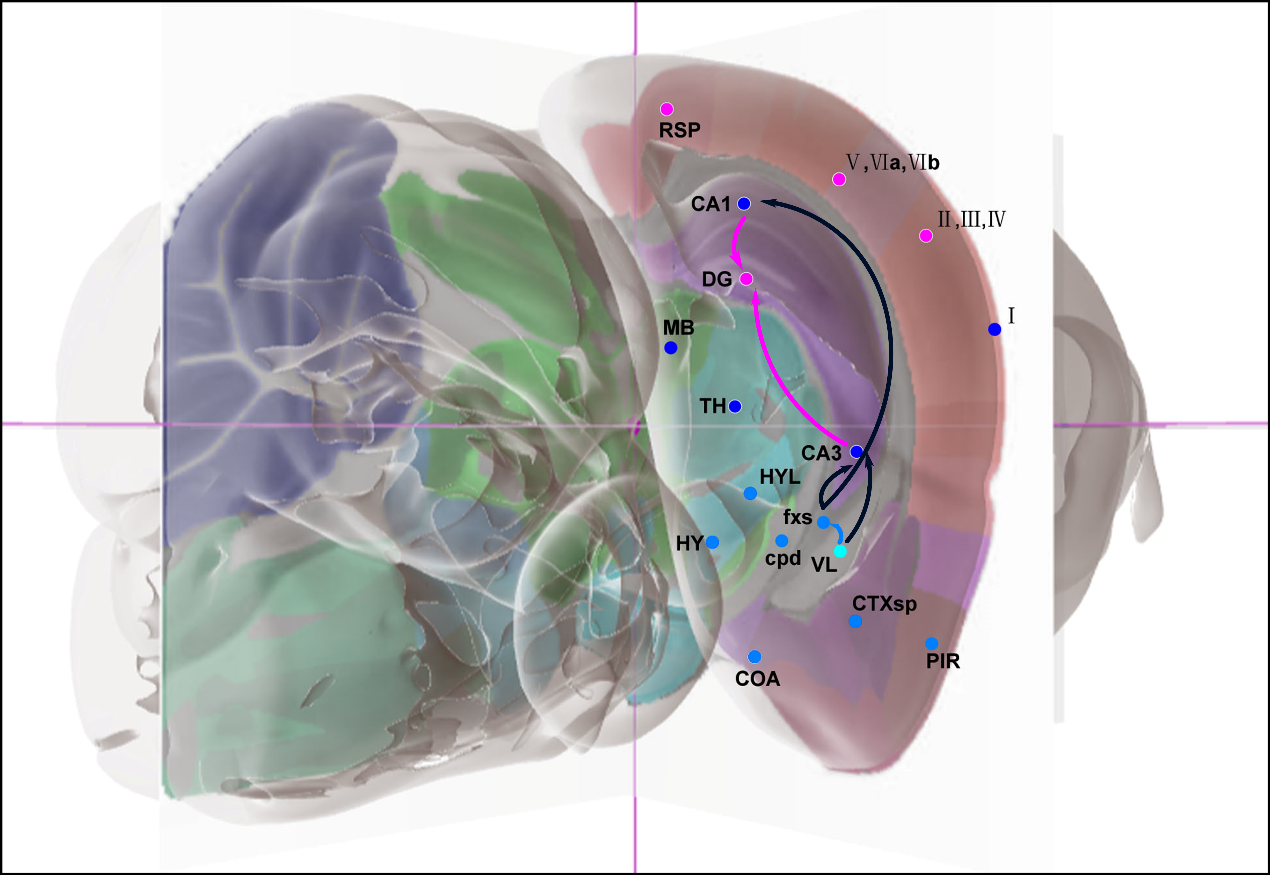

f. Trajectory bundle II of IVH_3d

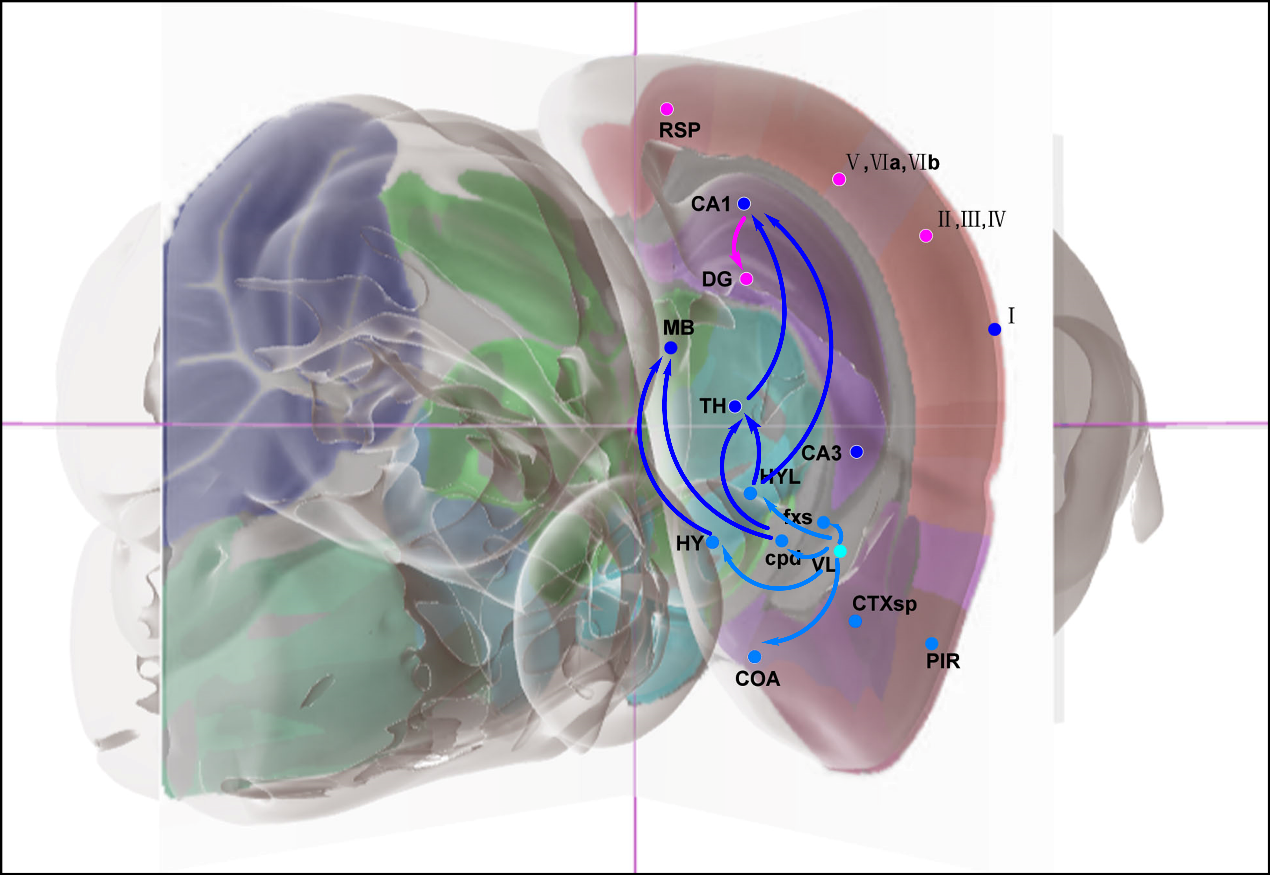

g. Trajectory bundle III of IVH_3d

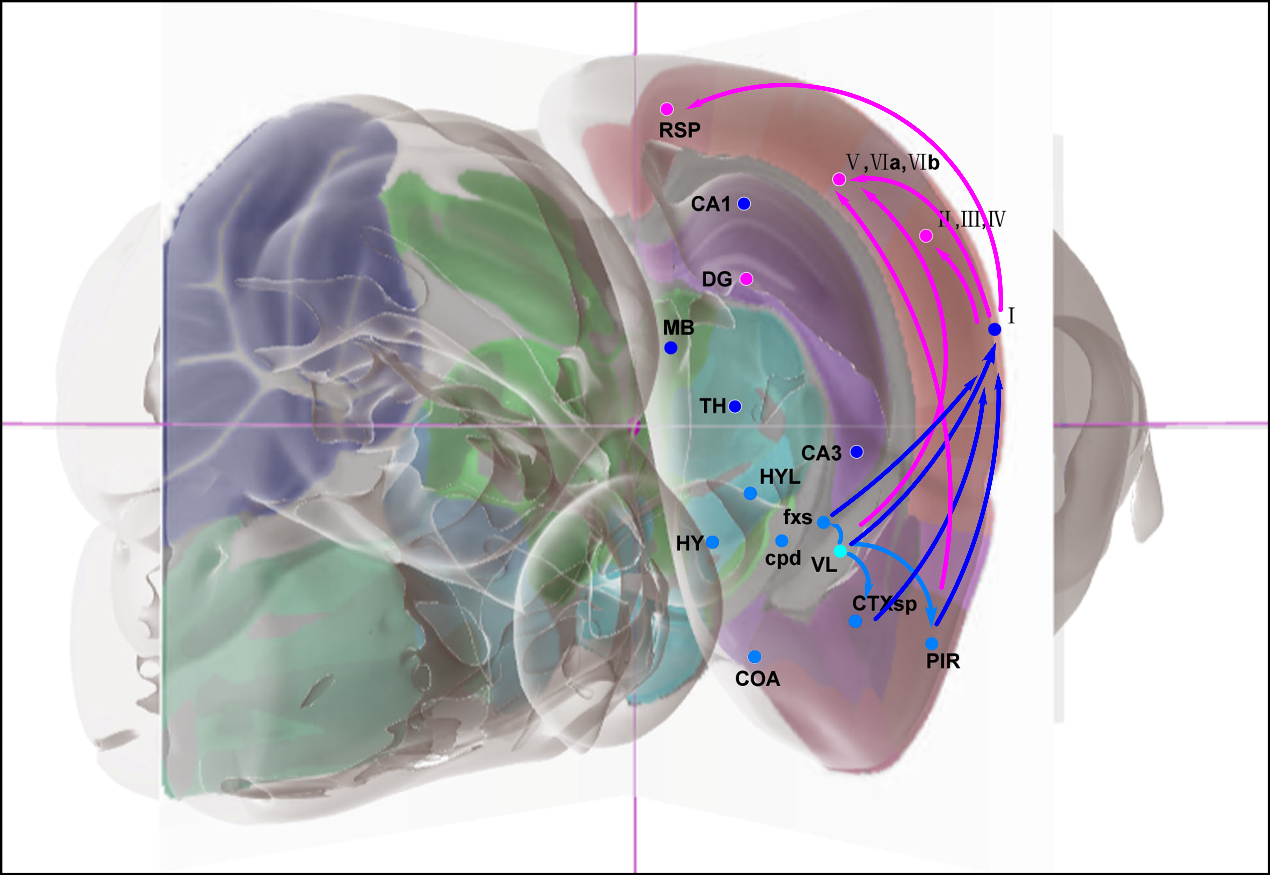

h. Trajectory bundle I of IVH_7d

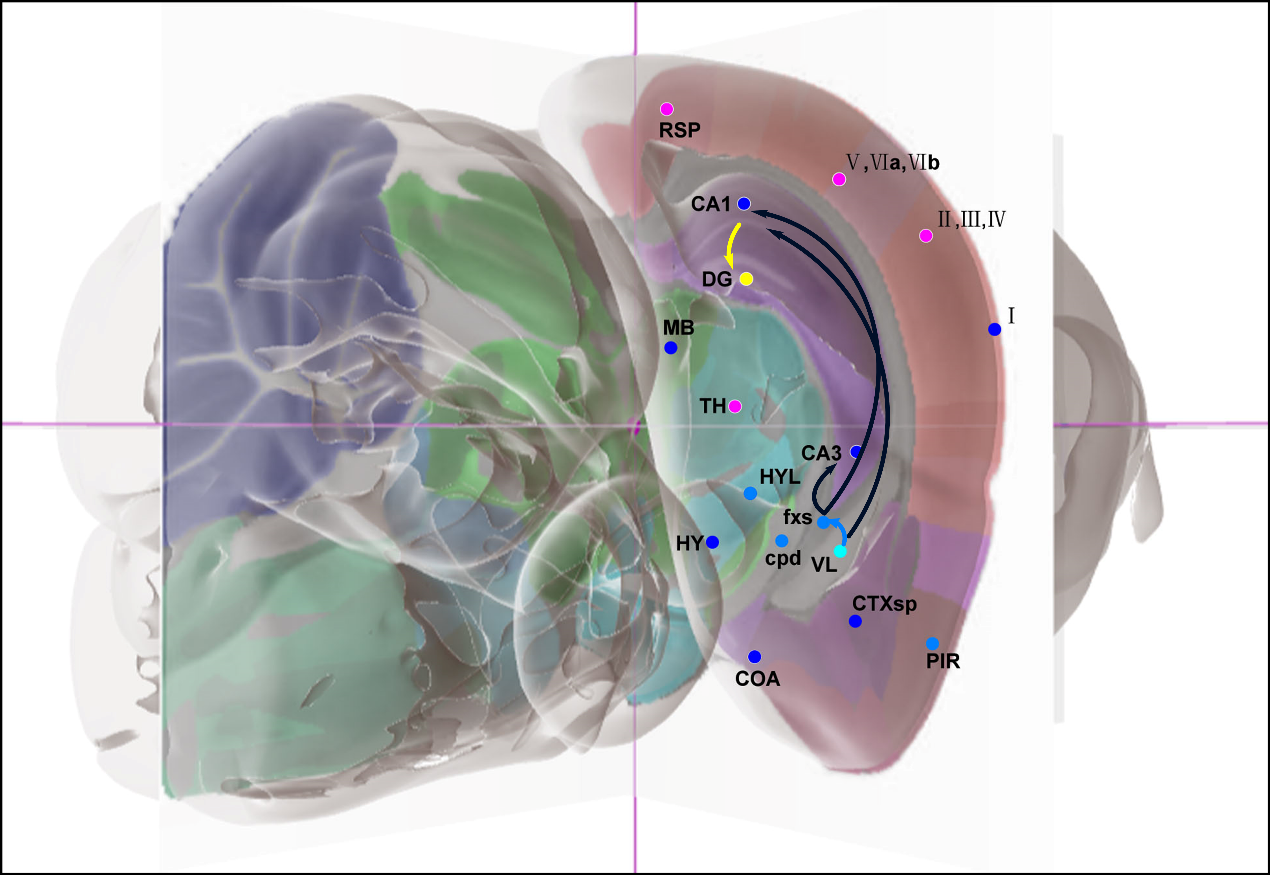

i. Trajectory bundle II of IVH_7d

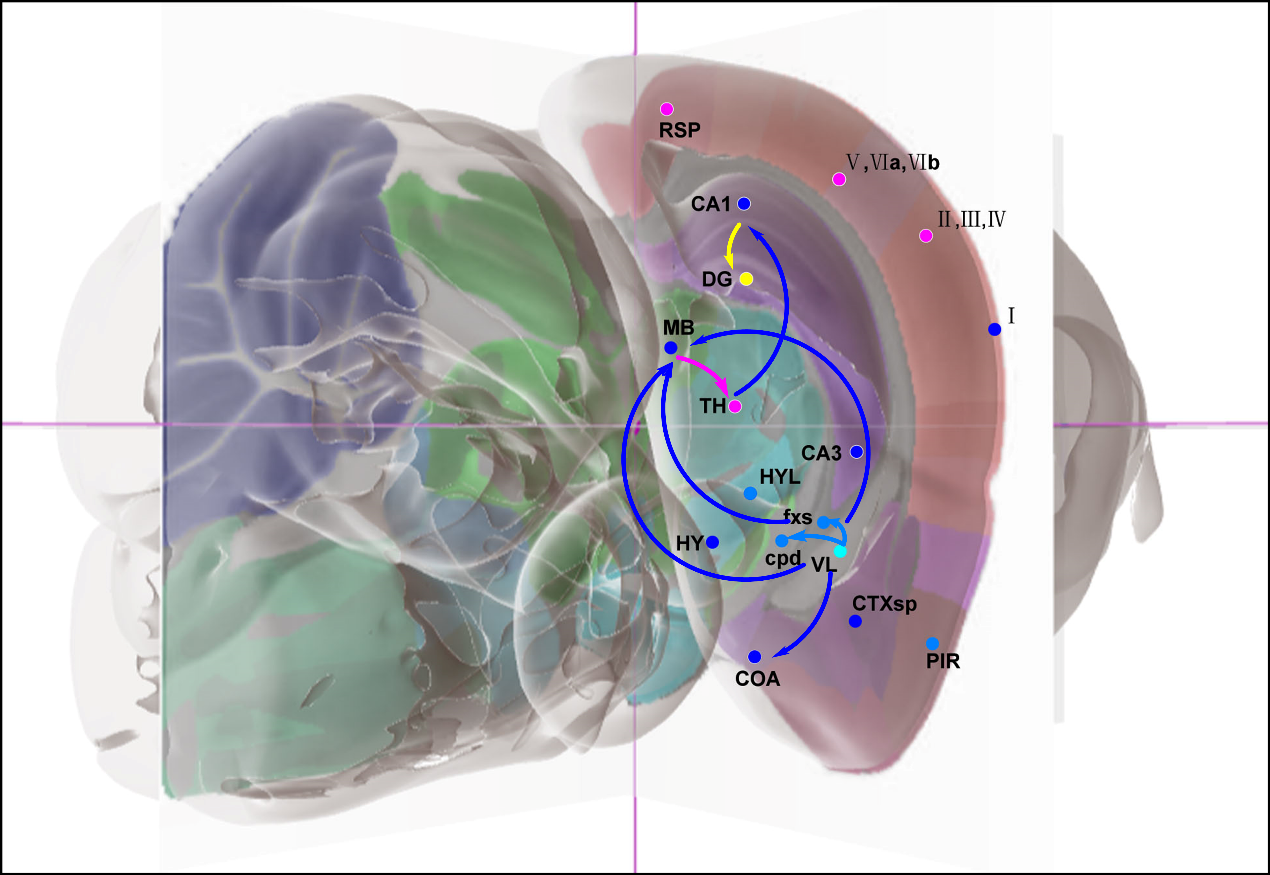

j. Trajectory bundle III of IVH_7d

3. **Supplementary for Result 3 (SR3)**

Table 1 The planar upregulated and downregulated transition genes for each planner sub-trajectory at different time

[*https://github.com/JiayidaerBadai/Spatial-transcriptome.git*](https://github.com/JiayidaerBadai/Spatial-transcriptome.git)

a. IVH_1d
b. IVH_3d
c. IVH_7d
d. IVH_S1

Table 2.1 The top 30 3D upregulated transition genes for each 3D sub-trajectory at different time

[*https://github.com/JiayidaerBadai/Spatial-transcriptome.git*](https://github.com/JiayidaerBadai/Spatial-transcriptome.git)

a. IVH_1d
b. IVH_3d
c. IVH_7d
d. IVH_S1

Table 2.2 The top 30 3D downregulated transition genes for each 3D sub-trajectory at different time

[*https://github.com/JiayidaerBadai/Spatial-transcriptome.git*](https://github.com/JiayidaerBadai/Spatial-transcriptome.git)

a. IVH_1d
b. IVH_3d
c. IVH_7d
d. IVH_S1
4. **Supplementary for Result 4 (SR4)**

Table 1 Cell subtypes corresponding the selected trajectory at different time

[*https://github.com/JiayidaerBadai/Spatial-transcriptome.git*](https://github.com/JiayidaerBadai/Spatial-transcriptome.git)

Table 2 The marker gene sets of cell subtypes corresponding the selected trajectory at different time

[*https://github.com/JiayidaerBadai/Spatial-transcriptome.git*](https://github.com/JiayidaerBadai/Spatial-transcriptome.git)

Table 3 The similarity between cell subtypes at different time on the same trajectory

[*https://github.com/JiayidaerBadai/Spatial-transcriptome.git*](https://github.com/JiayidaerBadai/Spatial-transcriptome.git)

Table 4 The cell types that are as same as cell subtypes corresponding the selected trajectory at different time

[*https://github.com/JiayidaerBadai/Spatial-transcriptome.git*](https://github.com/JiayidaerBadai/Spatial-transcriptome.git)

Table 5 The cell types shared by our identified cell subtypes at different time on the same trajectory.

[*https://github.com/JiayidaerBadai/Spatial-transcriptome.git*](https://github.com/JiayidaerBadai/Spatial-transcriptome.git)

5. **Supplementary for Result 5 (SR5)**

Table 1 Upregulated/downregulated transition ligand-receptor sets

[*https://github.com/JiayidaerBadai/Spatial-transcriptome.git*](https://github.com/JiayidaerBadai/Spatial-transcriptome.git)

1.1： IVH_1d upregulated transition ligand-receptor sets

1.2： IVH_1d downregulated transition ligand-receptor sets

1.3： IVH_3d upregulated transition ligand-receptor sets

1.4： IVH_3d downregulated transition ligand-receptor sets

1.5： IVH_7d upregulated transition ligand-receptor sets

1.6： IVH_7d downregulated transition ligand-receptor sets

1.7 ：IVH_S1 upregulated transition ligand-receptor sets

1.8 ：IVH_S1 downregulated transition ligand-receptor sets

Table 2 The interaction intensity value $Zscore$ (CCI) for the upregulated/downregulated transition ligand-receptor sets.

[*https://github.com/JiayidaerBadai/Spatial-transcriptome.git*](https://github.com/JiayidaerBadai/Spatial-transcriptome.git)

2.1： The interaction intensity value $Zscore$ (CCI) for the IVH_1d upregulated transition ligand-receptor sets

2.2： The interaction intensity value $Zscore$ (CCI) for the IVH_1d downregulated transition ligand-receptor sets

2.3： The interaction intensity value $Zscore$ (CCI) for the IVH_3d upregulated transition ligand-receptor sets

2.4： The interaction intensity value $Zscore$ (CCI) for the IVH_3d downregulated transition ligand-receptor sets

2.5： The interaction intensity value $Zscore$ (CCI) for the IVH_7d upregulated transition ligand-receptor sets

2.6： The interaction intensity value $Zscore$ (CCI) for the IVH_7d downregulated transition ligand-receptor sets

2.7： The interaction intensity value $Zscore$ (CCI) for the IVH_S1 upregulated transition ligand-receptor sets

2.8： The interaction intensity value $Zscore$ (CCI) for the IVH_S1 downregulated transition ligand-receptor sets

Table 3 The interaction value $ZT$ converted from $Zscore$ (CCI) for the upregulated/downregulated transition ligand-receptor sets.

[*https://github.com/JiayidaerBadai/Spatial-transcriptome.git*](https://github.com/JiayidaerBadai/Spatial-transcriptome.git)

3.1： The interaction intensity value $ZT$ for the IVH_1d upregulated transition ligand-receptor sets

3.2： The interaction intensity value $ZT$ for the IVH_1d downregulated transition ligand-receptor sets

3.3： The interaction intensity value $ZT$ for the IVH_3d upregulated transition ligand-receptor sets

3.4： The interaction intensity value $ZT$ for the IVH_3d downregulated transition ligand-receptor sets

3.5： The interaction intensity value $ZT$ for the IVH_7d upregulated transition ligand-receptor sets

3.6： The interaction intensity value $ZT$ for the IVH_7d downregulated transition ligand-receptor sets

3.7： The interaction intensity value $ZT$ for the IVH_S1 upregulated transition ligand-receptor sets

3.8： The interaction intensity value $ZT$ for the IVH_S1 downregulated transition ligand-receptor sets

Table 4 The $Density$ for each ligand-receptor of cell subtypes

[*https://github.com/JiayidaerBadai/Spatial-transcriptome.git*](https://github.com/JiayidaerBadai/Spatial-transcriptome.git)

4.1： The average $Density$ for each ligand-receptor of IVH_1d upregulated transition ligand-receptor sets

4.2：The average $Density$ for each ligand-receptor of IVH_1d downregulated transition ligand-receptor sets

4.3： The average $Density$ for each ligand-receptor of IVH_3d upregulated transition ligand-receptor sets

4.4：The average $Density$ for each ligand-receptor of IVH_3d downregulated transition ligand-receptor sets

4.5： The average $Density$ for each ligand-receptor of IVH_7d upregulated transition ligand-receptor sets

4.6：The average $Density$ for each ligand-receptor of IVH_7d downregulated transition ligand-receptor sets

4.7： The average $Density$ for each ligand-receptor of IVH_S1 upregulated transition ligand-receptor sets

4.8：The average $Density$ for each ligand-receptor of IVH_S1 downregulated transition ligand-receptor sets

Table 5 The Kruskal-Wallis test for Density among these cell subtypes respectively on trajectory $VL\to fxs\to CA1\to DG$ , $TH\to CA1\to DG$ and $VL\to I\to(V,VIa,VIb)$.

[*https://github.com/JiayidaerBadai/Spatial-transcriptome.git*](https://github.com/JiayidaerBadai/Spatial-transcriptome.git)

6. **Supplementary for Result 6 (SR6)**

Figure 1 The heatmap of mutual pathway sets for our identified cell subtypes

1.1 The heatmap of mutual pathway sets for trajectory VL→fxs→CA1→DG

1.2 The heatmap of mutual pathway sets for trajectory TH→CA1→DG

1.3 The heatmap of mutual pathway sets for trajectory VL→I→(V,VIa,VIb)

Figure 2 The heatmap of mutual pathway sets for similar cell types.

2.1 The heatmap of mutual pathway sets for similar cell types of trajectory VL→fxs→CA1→DG

2.2 The heatmap of mutual pathway sets for similar cell types of trajectory TH→CA1→DG

2.3 The heatmap of mutual pathway sets for similar cell types of trajectory VL→I→(V,VIa,VIb)

Table 1 Marker genes of similar cell types for our identified cell subtypes

[*https://github.com/JiayidaerBadai/Spatial-transcriptome.git*](https://github.com/JiayidaerBadai/Spatial-transcriptome.git)

Table 2 The mutual pathway sets for our identified cell subtypes

[*https://github.com/JiayidaerBadai/Spatial-transcriptome.git*](https://github.com/JiayidaerBadai/Spatial-transcriptome.git)

2.1 The mutual pathway sets for trajectory VL→fxs→CA1→DG

2.2 The mutual pathway sets for trajectory TH→CA1→DG

2.3 The mutual pathway sets for trajectory VL→I→(V,VIa,VIb)

Table 3 The mutual pathway sets for similar cell types

*<https://github.com/JiayidaerBadai/Spatial-transcriptome.git>*

3.1 The mutual pathway sets for similar cell types of trajectory VL→fxs→CA1→DG

3.2 The mutual pathway sets for similar cell types of trajectory TH→CA1→DG

3.3 The mutual pathway sets for similar cell types of trajectory VL→I→(V,VIa,VIb)

Table 4 The mutual pathway set between our identified cell subtypes and similar cell types [*https://github.com/JiayidaerBadai/Spatial-transcriptome.git*](https://github.com/JiayidaerBadai/Spatial-transcriptome.git)

Supplementary of Methods

This supplementary of Methods section consists of 11 supplementary files, which is listed as below.

Figure1 IVH model

Table1 IVH sample grouping, library patching strategy and permeabilization time（min）

| group | sample | Permeabilization time（min） |
| --- | --- | --- |
| 1. IVH_1d | IVH_1d_1_20 | 25.5min |
|  | IVH_1d_1_50 | 25.5min |
|  | IVH_1d_1_80 | 25.5min |
|  | IVH_1d_1_110 | 25.5min |
|  | IVH_1d_1_130 | 25.5min |
|  | IVH_1d_1_150 | 25.5min |
| 2. IVH_3d | IVH_3d_3_20 | 19min |
|  | IVH_3d_3_50 | 19min |
|  | IVH_3d_3_110 | 19min |
|  | IVH_3d_3_130 | 19min |
|  | IVH_3d_3_150 | 19min |
| 3. IVH_7d | IVH_7d_2_50 | 25min |
|  | IVH_7d_2_80 | 25min |
|  | IVH_7d_2_110 | 25min |
| 4. IVH_C1 | IVH_C1_50 | 25min |
|  | IVH_C1_110 | 25min |
| 5. IVH_S1 | IVH_S1_50 | 25min |
|  | IVH_S1_80 | 25min |
|  | IVH_S1_110 | 25min |

Table 2 Parameter Definition for 3D global Pseudo-space-time trajectory reconstruction algorithm

| No. | Definition | Description |
| --- | --- | --- |
| 1. | ${Time}_{t} ,t\in[1,5]$ | Time t after IVH |
| 2. | ${slide}_{n} ,n\in[1,N]$ | Slide n. |
| 3. | ${Trajectory}^{n}\equiv G^{n}=(V^{n},E^{n})$, $n\in[1,N]$ | The set of planar pseudo-space-time trajectories on the slide n, equivalent to a Directed Acyclic Graph $G^{n}$ on the slide n which is composed of nodes $V^{n}$ and directed edges $E^{n}$. |
| 4. | $V^{n}$($v_{i}^{n}\in V^{n},i\in[1,N])$ | Nodes $V^{n}$ on the graph $G^{n}$ represents brain regions on the slide n, $v_{i}^{n}$ represents brain region I on the slide n. |
| 5. | $E^{n}(e_{ij}^{n}\in E^{n}, i\in[1,N], j\in[1,N])$ | Directed edges $E^{n}$ on the graph $G^{n}$ represents the set of directed edge $e_{ij}^{n}$ from the brain region I to the brain region j on the slide n. |
| 6. | ${DPT}_{v_{i}^{n}}^{m}$, $n\in\left[ 1,N \right]$*, m* $\in\left[ 1,N \right]$*,* $i\in[1,N]$*,* ${DPT}_{v_{i}^{n}}^{m}\in[0,1]$ | ${DPT}_{v_{i}^{n}}^{m}$ represents the diffusion pseudotime of each spot in node (brain region) *i.* |
| 7. | $Avg {DPT}_{v_{i}^{n}}=\frac{\sum_{m=1}^{N} {DPT}_{v_{i}^{n}}^{m}}{m}$ (1.1) | The average diffusion pseudotime for each planar brain region. |
| 8. | $3D\_ {dpt}_{v_{i}}=\frac{\sum_{n=1}^{N} Avg {DPT}_{v_{i}^{n}}}{N}$ (1.2) | 3D diffusion pseudotime of brain region *i.* |
| 9. | imagerow ($x_{v_{i}^{n}}^{m}$), imagecol ($y_{v_{i}^{n}}^{m}$) | Planar coordinates of each spot in node (brain region) *i.* |
| 10. | $z_{v_{i}^{n}}$=$\left\{ \begin{aligned} n=1 \left( {Time}_{t}\_20 \right), 1 \\ n=2 \left( {Time}_{t}\_50 \right), 2 \\ n=3 \left( {Time}_{t}\_80 \right), 3 \\ n=4 \left( {Time}_{t}\_110 \right), 4 \\ n=5 \left( {Time}_{t}\_130 \right), 5 \\ n=6\left( {Time}_{t}\_150 \right), 6 \end{aligned} \right\}$ | Height coordinates of node (brain region) *I* of each slide*.* |
| 11. | ${(Avg\_x}_{v_{i}^{n}}^{m},{Avg\_y}_{v_{i}^{n}}^{m})=$($\frac{{(x}_{v_{i}^{n}}^{1}+x_{v_{i}^{n}}^{2}+\ldots+x_{v_{i}^{n}}^{m})}{m},\frac{{(y}_{v_{i}^{n}}^{1}+y_{v_{i}^{n}}^{2}+\ldots+y_{v_{i}^{n}}^{m})}{m}$) (1.3) | Planar Centroid coordinates of node $v_{i}$ on the slide n with m spots. |
| 12. | $v_{1}^{n}$ | $v_{1}^{n}$ represents the starting node (brain region) of the set of planar pseudo-space-time trajectories on the slide n. |
| 13. | ${(x}_{i},y_{i},z_{i})=$ ($\frac{{(x}_{v_{i}^{1}}^{m}+x_{v_{i}^{2}}^{m}+\ldots+x_{v_{i}^{n}}^{m})}{n},\frac{{(y}_{v_{i}^{1}}^{m}+y_{v_{i}^{2}}^{m}+\ldots+y_{v_{i}^{n}}^{m})}{n}$) (1.4) | 3D Centroid coordinates of $v_{i}$ |
| 14. | $v_{f}^{n}$，$f\in F^{n}$ | $v_{f}^{n}$ represents ending node (brain region) of planar pseudo-space-time trajectory on the slide n. $F^{n}$ represents ending node brain region set of planar pseudo-space-time trajectories. |
| 15. | $v_{i}=\left( l_{i},p_{i} \right),$  $l_{i}\in[1,5]$*,* $p_{i}\in[1,3]$ | $l_{i}\in[1,5]$ represents that the node (brain region) $v_{i}$ is divided into 5 levels according to the average diffusion pseudotime of brain region. $p_{i}\in[1,3]$ represents that according to the trajectory (path) from the starting node 1 to the ending node $f$, node $v_{i}$ on trajectory $T_{1-..-i-..-f}^{n}{\in Trajectory}^{n}$ is divided into 3 categories. $p_{1}$ contains “fxs”, “CA3”, “CA1”, “DG”.  $p_{2}$ contains “COA”, “HY”, “cpd”, “fxs”, “HYL”, “TH”, “MB”, “CA1”, “CA3”, “DG”. $p_{3}$ contains “CTXsp”, “PIR”, “I”, “II_III_IV”, “V_VIa_VIb”, “RSP”. |
| 16. | ${Trajectory}^{common}$ | A common set of 3D pseudo-space-time trajectories. |
| 17. | $B^{n}$, $n\in[1,N]$ | The starting nodes of the set of planar pseudo-space-time trajectories ${Trajectory}^{n}$. |
| 18. | $C^{n}$, $n\in[1,N]$ | Descendant node set. |
| 19. | $H^{n}$, $n\in[1,N]$ | Candidate descendant node set. |
| 20. | $D^{n}$ | Final descendant node set. |
| 21. | ${dis}_{ij}=\sqrt{{(x_{i}-x_{j})}^{2}+{(y_{i}-y_{j})}^{2}+{(z_{i}-z_{j})}^{2}} (1.5)$ | Distance between node I and node j, $i\in[1,N]$, j$\in[1,N]$ |
| 22. | ${Dis}_{i}$ | Distance set of node i. |

Table 3 3D global Pseudo-space-time trajectory reconstruction algorithm

| **Algorithm** 3D Global Pseudo-space-time Trajectory Reconstruction Algorithm | |
| --- | --- |
| 1: | **Input:** ${Trajectory}^{n}$, $v_{i}^{n}$，$e_{i}^{n}$，$p_{i}$, ${DPT}_{v_{i}^{n}}^{m}$，imagerow ($x_{v_{i}^{n}}^{m}$), imagecol ($y_{v_{i}^{n}}^{m}$) |
| 2: | **For** each pseudo-space-time trajectories index *n*： |
| 3: | **For** each brain region *I* in pseudo-space-time trajectories： |
| 4: | Calculate the the average diffusion pseudotime $Avg {DPT}_{v_{i}^{n}}$ by Eq.1.1 |
| 5: | **For** each brain region *I* in pseudo-space-time trajectories： |
| 6: | Calculate the 3D diffusion pseudotime $3D\_ {dpt}_{v_{i}}$ by Eq.1.2 |
| 7: | Divide the $3D\_ {dpt}_{i}$ by step size (0.0889104 for brain regions in $p_{3}$ and 0.1462136 for brain regions in $p_{1}$& $p_{2}$) to obtain the diffusion pseudotime level $l_{i}$ for each 3D brain region from 0 to 6 level. |
| 8: | Insert $v_{1}$ into set ${Trajectory}^{common}$ |
| 9: | **For** each pseudo-space-time trajectories index *n* in $p_{i}$： |
| 10: | Insert $v_{1}$ into set $B^{n}$ |
| 11: | **For** *length* from *1* to *max_length*： |
| 12: | $Min\_length ={\{\infty\}}^{N}$, $min\_index ={\{0\}}^{N}$,$C={\{\emptyset\}}^{N}$,$H={\{\emptyset\}}^{N}$,$D={\{\emptyset\}}^{N}$ |
| 13: | **For** each pseudo-space-time trajectories index *n*： |
| 14: | **For** each vector *I* in $B^{n}$： |
| 15: | **For** each vector *j* in $e_{i}^{n}$： |
| 16: | Insert vector *j* into set $C^{n}$ |
| 17: | **If** $l_{i}=length+n$： |
| 18: | Insert vector *j* into set $H^{n}$ |
| 19: | **If** $v_{i}== "TH"$ , $v_{j}== "CA1"$ or $v_{j}== "CA3"$ |
| 20: | Insert vector *j* into set $D^{n}$ |
| 21: | Calculate the 3D Centroid coordinates of $v_{i}$ by Eq.1.3-1.4, and distance ${dis}_{ij}^{n}$ between vector *I* and vector *j* by Eq.1.5 |
| 22: | I**f** ${dis}_{ij}^{n}<Min\_length$： |
| 23: | ${Min\_length}^{n}={dis}_{ij}^{n}$, ${min\_index}^{n}=j$ |
| 24: | **If** $\bigcup_{k=1}^{N} D^{k}=\emptyset$： |
| 25: | $D=min\_index$ |
| 26: | **If** $\bigcup_{k=1}^{N} C^{k}=\emptyset$： |
| 27: | Break |
| 28: | **Else if** $\bigcup_{k=1}^{N} H^{k}=\emptyset$： |
| 29: | Help |
| 30: | **Else**： |
| 31: | **For** each pseudo-space-time trajectories index *n*： |
| 32: | **For** each vector *I* in $B^{n}$： |
| 33: | **For** each vector *j* in $D^{n}$： |
| 34: | Insert edge $e_{ij}$ into ${Trajectory}^{common}$ |
| 35: | $B^{n}=D^{n}$ |
| 36: | **Output**: ${Trajectory}^{common}$ |

Table 4 Parameter Definition for the algorithm to identify a cell subtype and Similarity algorithm for cell subtypes

| No. | Definition | Description |
| --- | --- | --- |
| 1. | ${CellT}_{\mathrm{Trajectory}_{i}}^{{time}_{n}}$ | A cell subtype on $\mathrm{Trajectory}_{i}$ at different time, time represents the day (Sham, 1d, 3d and 7d). (SR4.Table 1) |
| 2. | ${GeneS}_{\mathrm{Trajectory}_{i}}^{{time}_{n}}=\left( GeneNames,Symbols \right)$ | A set of marker genes corresponding to  ${CellT}_{\mathrm{Trajectory}_{i}}^{{time}_{n}}$. (SR4.Table 2) |
| 3. | ${Gp}_{\mathrm{Trajectory}_{i}}^{{time}_{n}}$ | The top 30 upregulated transition genes of 3D sub-trajectory $\mathrm{Trajectory}_{i}$ at ${time}_{n}$. (SR3. Table 2.1) |
| 4. | ${Gn}_{\mathrm{Trajectory}_{i}}^{{time}_{n}}$ | The top 30 downregulated transition genes of 3D sub-trajectory $\mathrm{Trajectory}_{i}$ at ${time}_{n}$. (SR3. Table 2.2) |
| 5. | ${Gp}_{a->b}^{1d}\cup{Gp}_{b->c}^{1d}={Gp}_{a->b->c}^{1d}$ (1.1) | The upregulated transition genes of 3D trajectory $a->b->c$ at the first day. |
| 6. | ${Gn}_{a->b}^{1d}\cup{Gn}_{b->c}^{1d}={Gn}_{a->b->c}^{1d}$ (1.2) | The downregulated transition genes of 3D trajectory $a->b->c$ at the first day. |
| 7. | ${Gp}_{a->b->c}^{1d}\cup{Gn}_{a->b->c}^{1d}=G_{a->b->c}^{1d}=\{g1+,g2-,g3+, \ldots gi+\}$ (1.3) | $G_{a->b->c}^{1d}$ describes a cell subtype in the trajectory (a->b->c) at the first day. $gi$ Represents the gene, sign describes upregulation or downregulation. |
| 8. | $GeneNames=\{g_{0}，g_{1}，g_{2}\ldots g_{j}\}$ (2.1) | Represents the set of gene names, the element of the set is represented by $g_{j}$. |
| 9. | $Symbols=\{s_{0}，s_{1}，s_{2}\ldots s_{j}\}$ (2.2) | Represents the set of symbols, the element of the set is represented by $s_{j}$. The positive and negative of $s_{j}$ represents whether it is positively or negatively correlated with the key trajectory. |
| 10. | ${Similarity}_{\mathrm{Trajectory}_{i}}^{{time}_{m\_n}}=\left（ \frac{{GeneS}_{\mathrm{Trajectory}_{i}}^{{time}_{m}}\cdot{GeneS}_{\mathrm{Trajectory}_{i}}^{{time}_{n}}}{\left\vert{GeneS}_{\mathrm{Trajectory}_{i}}^{{time}_{m}}\cup{GeneS}_{\mathrm{Trajectory}_{i}}^{{time}_{n}} \right\vert}+1 \right）/2$ (2.3)  ${GeneS}_{\mathrm{Trajectory}_{i}}^{{time}_{m}}\cdot{GeneS}_{\mathrm{Trajectory}_{i}}^{{time}_{n}}=\sum\left( g_{j_{m}},s_{j_{m}} \right)\cdot\left( g_{j_{n}},s_{j_{n}} \right)$ (2.4)  $\left( g_{j_{m}},s_{j_{m}} \right)\cdot\left( g_{j_{n}},s_{j_{n}} \right)=\left\{ \begin{aligned} &\begin{matrix} -1, & \text{ (}s_{j_{m}}!=s_{j_{n}} \end{matrix})\text{\&\&(}g_{j_{m}}\text{=}g_{j_{n}}\text{)} \\ &\begin{matrix} 0, & {(g}_{j_{m}}\notin\left( g_{j_{m}}\cap g_{j_{n}} \right) \end{matrix}\text{\&\&}{(g}_{j_{n}}\notin\left( g_{j_{m}}\cap g_{j_{n}} \right) \\ &\begin{matrix} 1, & \text{ (}s_{j_{m}}=s_{j_{n}} \end{matrix})\text{\&\&(}g_{j_{m}}\text{=}g_{j_{n}}\text{)} \end{aligned} \right.)$ (2.5) | Eq. 2.3-2.5 calculate the similarity between different cell subtypes ${CellT}_{\mathrm{Trajectory}_{i}}^{{time}_{n}}$.  (Figure 4 and SR4.Table 3) |

Table 5 The algorithm to identify a cell subtype

| **Algorithm** The algorithm to identify a cell subtype | |
| --- | --- |
| 1: | **Input:** $\mathrm{Trajectory}_{i}$ (Trajectory a->b->c),${Gp}_{\mathrm{Trajectory}_{i}}^{{time}_{n}}$, ${Gn}_{\mathrm{Trajectory}_{i}}^{{time}_{n}}$ |
| 2: | **For** each sub trajectory ($a->b, b->c)$ in $\mathrm{Trajectory}_{i}$ , $index n$： |
| 3: | Calculate the upregulated transition genes ${Gp}_{\mathrm{Trajectory}_{i}}^{{time}_{n}}$ (${Gp}_{a->b->c}^{1d}$) by Eq.1.1 |
| 4: | Calculate the downregulated transition genes ${Gn}_{\mathrm{Trajectory}_{i}}^{{time}_{n}}$ (${Gn}_{a->b->c}^{1d})$ by Eq.1.2 |
| 5: | **For** ${Gp}_{\mathrm{Trajectory}_{i}}^{{time}_{n}}$ (${Gp}_{a->b->c}^{1d}$) and ${Gn}_{\mathrm{Trajectory}_{i}}^{{time}_{n}}$ (${Gn}_{a->b->c}^{1d})$ ： |
| 6: | Calculate the union of these two sets${GeneS}_{\mathrm{Trajectory}_{i}}^{{time}_{n}} (G_{a->b->c}^{1d})$ by Eq.1.3, if different plane slices had the same gene with a different sign (upregulated or downregulated), we chose the gene whose absolute value was the greatest |
| ,7: | **Output**: ${GeneS}_{\mathrm{Trajectory}_{i}}^{{time}_{n}}$ |

Table 6 Similarity algorithm for cell subtypes

| **Algorithm** Similarity algorithm for cell subtypes | |
| --- | --- |
| 1: | **Input:** ${GeneS}_{\mathrm{Trajectory}_{i}}^{{time}_{n}} ,$ $GeneNames=\left\{ g_{0}，g_{1}，g_{2}\ldots g_{j} \right\}, Symbols=\left\{ s_{0}，s_{1}，s_{2}\ldots s_{j} \right\}$ $Eq.2.1-2.2$ |
| 2: | **For** each ${GeneS}_{\mathrm{Trajectory}_{i}}^{{time}_{n}} ,index i,n$： |
| 3: | Calculate the similarity ${Similarity}_{\mathrm{Trajectory}_{i}}^{{time}_{m\_n}}$ between cell subtypes ${CellT}_{\mathrm{Trajectory}_{i}}^{{time}_{n}}$ at different time on the same trajectory by Eq.2.3-2.5 |
| 4: | **Output**: ${Similarity}_{\mathrm{Trajectory}_{i}}^{{time}_{m\_n}}$ |

Table 7 Parameter Definition for Cell-cell communication strength (Density) algorithm

| No. | Definition | Description |
| --- | --- | --- |
| 1. | ${LR}_{n}$ | Ligand-receptor n. (SR5 Table 1.1 – 1.8) |
| 2. | ${Zscore}_{i}^{{LR}_{n}},i\in(1,N)$ | Represents the interaction intensity of ligand-receptor (${LR}_{n}$) at each point $i$ on the section. (SR5 Table 2.1 - 2.8) |
| 3. | ${ZT}_{i}^{{LR}_{n}}$ | Represents the discrete interaction intensity after conversion of the continuous interaction intensity ${Zscore}_{i}^{{LR}_{n}}$. (SR5 Table 3.1-3.8) |
| 4. | $ZTTotal$ | Represents the sum of the interaction intensity ${ZT}_{i}^{{LR}_{n}}$ of N points in the corresponding position. (SR5 Table 3.1-3.8) |
| 5. | $A$ | Represents the number of points for each brain region pair. |
| 6. | $B$ | Represents the number of points for all brain regions. |
| 7. | ${Density}^{{LR}_{n}}$ | Represents the density of the interaction intensity of the ligand-receptor (${LR}_{n}$) of cell type over the interaction intensity of all brain regions. (SR5 Table 4.1-4.8) |
| 8. | ${ZT}_{i}^{{LR}_{n}} =\left\{ \begin{aligned} {Zscore}_{i}^{{LR}_{n}}<-3 0 \\ \begin{aligned} -3\leq{Zscore}_{i}^{{LR}_{n}}<-2 1 \\ -2\leq{Zscore}_{i}^{{LR}_{n}}<-1 2 \\ -1\leq{Zscore}_{i}^{{LR}_{n}}<0 3 \\ 0\leq{Zscore}_{i}^{{LR}_{n}}<1 4 \\ 1\leq{Zscore}_{i}^{{LR}_{n}}<2 5 \\ 2\leq{Zscore}_{i}^{{LR}_{n}}<3 6 \\ 3\leq{Zscore}_{i}^{{LR}_{n}}<4 7 \\ 4\leq{Zscore}_{i}^{{LR}_{n}}<5 8 \\ 5\leq{Zscore}_{i}^{{LR}_{n}}<6 9 \\ \ldots\ldots\ldots\ldots\\ n\leq{Zscore}_{i}^{{LR}_{n}}< n+1 (n\geq-3) n+4 \end{aligned} \end{aligned} \right\}$ (1.1) | Converting the ${Zscore}_{i}^{{LR}_{n}}$ into  ${ZT}_{i}^{{LR}_{n}}$. |
| 9. | $ZTTotal$ $=\sum_{i=1}^{N} {ZT}_{i}^{{LR}_{n}}$ (1.2) | Calculating the total interaction intensity $ZTTotal$ for each ligand-receptor (${LR}_{n}$) of cell subtypes and all the brain regions, respectively. |
| 10. | ${Density}^{{LR}_{n}}=\frac{ZTTotalA/A}{ZTTotalB/B}$ (1.3) | Calculating the average ${Density}^{{LR}_{n}}$ for each ligand-receptor (${LR}_{n}$) of cell subtypes. |

Table 8 Cell-cell communication strength (Density) algorithm

| **Algorithm** Cell-cell communication strength (Density) algorithm | |
| --- | --- |
| 1: | **Input:** ${Zscore}_{i}^{{LR}_{n}}$ |
| 2: | **For** each ${Zscore}_{i}^{{LR}_{n}}$ of each ligand-receptor (${LR}_{n}$) at each point $i$ on the cell subtypes (A points) and all the brain regions section (B points), index *n*： |
| 3: | Convert these two ${Zscore}_{i}^{{LR}_{n}}$ into ${ZT\_A}_{i}$ and ${ZT\_B}_{i}$ by Eq.1.1 |
| 4: | **For** each ${ZT\_A}_{i}$ and ${ZT\_B}_{i}$ of ${LR}_{n}$, index *i*： |
| 5: | Calculate the total interaction intensity $ZTTotalA$ and $ZTTotalB$ by Eq.1.2 |
| 6: | **For** $ZTTotalA$ and $ZTTotalB$ of ${LR}_{n}$： |
| 7: | Calculate the average ${Density}^{{LR}_{n}}$ for ${LR}_{n}$ of cell subtypes by Eq.1.3 |
| 8: | **Output**: ${Density}^{{LR}_{n}}$ |

Table 9 Parameter Definition for Similarity algorithm for mutual pathway sets

| No. | Definition | Description |
| --- | --- | --- |
| 1. | $P{\_A}_{{Trajectory}_{n}}$, ($n=1\ldots N)$ | Mutual pathway sets for our identified cell subtypes (SR6. Table 2.1-2.3 and SR6. Figure 1.1-1.3). |
| 2. | ${P\_B}_{{Trajectory}_{n}}$, ($n=1\ldots N)$ | Mutual pathway sets for similar cell types (SR6. Table 3.1-3.3 and SR6. Figure 2.1-2.3). |
| 3. | ${P\_inter}_{{Trajectory}_{n}}=P{\_A}_{{Trajectory}_{n}}\cap{P\_B}_{{Trajectory}_{n}}$ (1.1) | Mutual pathway set of our identified cell subtypes and similar cell types (SR6. Table 4). |
| 4. | ${Similarity}_{{Trajectory}_{n}}^{P_{A\_B}}$ $=\frac{{P\_inter}_{{Trajectory}_{n}}}{P{\_A}_{{Trajectory}_{n}}}$ (1.2) | The similarity between $P{\_A}_{{Trajectory}_{n}}$ and ${P\_B}_{{Trajectory}_{n}}$(SR6. Figure 3). |

Table 10 Similarity algorithm for mutual pathway sets

| **Algorithm** Similarity algorithm for mutual pathway sets | |
| --- | --- |
| 1: | **Input:** $P{\_A}_{{Trajectory}_{n}}, {P\_B}_{{Trajectory}_{n}}$, ($n=1\ldots N)$ |
| 2: | **For** each $P{\_A}_{{Trajectory}_{n}}$, ${P\_B}_{{Trajectory}_{n}}$ , index *n*： |
| 3: | Calculate the mutual pathway set ${P\_inter}_{{Trajectory}_{n}}$ between $P{\_A}_{{Trajectory}_{n}}$ and ${P\_B}_{{Trajectory}_{n}}$ by Eq.1.1 |
| 4: | **For** each ${P\_inter}_{{Trajectory}_{n}}$ , index *n*： |
| 5: | Calculate the similarity ${Similarity}_{{Trajectory}_{n}}^{P_{A\_B}}$ by Eq.1.2 |
| 6: | **Output**: ${P\_inter}_{{Trajectory}_{n}}$, ${Similarity}_{{Trajectory}_{n}}^{P_{A\_B}}$ |
